# Rat models of human diseases and related phenotypes: a systematic inventory of the causative genes

**DOI:** 10.1101/2020.03.23.003392

**Authors:** Claude Szpirer

**Affiliations:** Université Libre de Bruxelles, B-6041 Gosselies, Belgium (retired) Current address: Avenue Jassogne, 27; B-1410 Waterloo, Belgium

## Abstract

The rat has been used for a long time as the model of choice in several biomedical disciplines. Numerous inbred strains have been isolated, displaying a wide range of phenotypes and providing many models of human traits and diseases. Rat genome mapping and genomics was considerably developed in the last decades. The availability of these resources has stimulated numerous studies aimed at discovering disease genes by positional identification. Numerous rat genes have now been identified that underlie monogenic or complex diseases and remarkably, these results have been translated to the human in a significant proportion of cases, leading to the identification of novel human disease susceptibility genes, helping in studying the mechanisms underlying the pathological abnormalities and also suggesting new therapeutic approaches. In addition, reverse genetic tools have been developed. Several genome-editing methods were introduced to generate targeted mutations in genes the function of which could be clarified in this manner [generally these are knockout (KO) mutations]. Furthermore, even when the human gene causing a disease is identified, mutated rat strains (in particular KO strains) were created to analyze the gene function and the disease pathogenesis. Today, about 300 rat genes have been identified as underlying diseases or playing a key role in critical biological processes that are altered in diseases. This article provides the reader with an inventory of these genes.

---

Why map and identify genes for rat disease phenotypes or related traits? The rat is more than a bigger mouse, a species which has been the mammalian genetic model of choice for a long time, with an initial focus on monogenic traits [1–4]. Rat models of monogenic traits and diseases have also been isolated but the rat has essentially been a key model for studies of complex traits in fields such as physiology, including cardiovascular and diabetes research, arthritis, pharmacology, toxicology, oncology and neurosciences. The intermediate size of the rat allows one to carry out experiments and measurements that are difficult if not impossible in the mouse and the rat exhibits more sophisticated neurobehavioral traits; it is an important animal model in neuropsychiatric and behavioral studies; in some scientific fields, the rat thus provides one with particularly reliable models of human traits or diseases [5–9].

Consequently, many rat strains have been created by selective breeding of animals expressing a desired phenotype, generating a large collection of genetic models of pathological complex, polygenic traits, most of which are quantitative. Interestingly, these strains also provide one with additional phenotypes, which were not selected for. Just as the traits that were selected for, most of these phenotypes are polygenic. All these phenotypes can be used as models of human traits or diseases [10], implying that the genes underlying these traits or diseases should be identified. Information on rat strains and rat disease models, can be found at the Rat Genome Database (RGD, https://rgd.mcw.edu/) [11].

In order to give the rat the status of a valuable genetic model, and in particular to identify the genes underlying complex traits by forward genetic approaches and to analyze the relevant biological mechanisms, several tools had to be developed. This has been accomplished. Genetic and chromosome maps have been developed; the genomic sequence of several rat strains has been established; a number of resources have been created to provide investigators with access to genetic, genomic, phenotype and disease-relevant data as well as software tools necessary for their research [3, 12]. Thanks to these resources, positional identification of numerous genes underlying monogenic or complex diseases and related traits could be achieved. On the other hand, reverse genetic tools have also been developed. Efficient methods to generate mutant rats became available; sperm N-ethyl-N-nitrosourea (ENU) mutagenesis followed by gene-targeted screening methods lead to the isolation of several mutants, including knockout (KO) strains [13 and references therein]. Rat ES were successfully derived and could be used for targeted mutations by homologous recombination; more importantly, several methods not relying on the use of ES cells were introduced to generated targeted mutations (often these are KO mutations), namely gene editing by zinc finger nucleases, by transcription activator-like effector nucleases and finally by the clustered regularly interspaced short palindromic repeat (CRISPR/Cas) system [for a review, see 14]. Transgenic rats can also be generated, including humanized rats carrying large chromosomic fragments (“transchromosomic humanized” rats) [15]. Development of these technologies provides the researcher with all the tools required to take advantage of the unique opportunities offered by the rat as leading model for studies different areas of biomedical research [3, 8]. In this review I made an inventory of the rat genes identified as responsible for monogenic or polygenic diseases and related traits. I took into account the rat genes identified by forward genetic methods as well as those inactivated by ENU-mutagenesis and by targeted mutations, the inactivation of which generated a disease or an abnormal phenotype. This inventory shows that a considerable number of conserved genes have similar effects on biological traits in rats and humans.

## Materials and methods

The data were collected by regular and systematic screening of the biomedical literature, PubMed searches (https://www.ncbi.nlm.nih.gov/) and Google Scholar alerts based on the terms “knockout”, “mutation”, “rat”. In addition, relevant data were retrieved from the RGD, thanks to advices from Jennifer Smith. The official gene symbols are used in this article and were obtained from the National Center for Biotechnology Information (https://www.ncbi.nlm.nih.gov/), Gene section. In several instances the original publications did not use the official gene symbol; in these cases, the non-official symbol is indicated in parenthesis in the footnote to the table, where the full name of each gene is described. The position of every gene was also obtained from the NCBI.

## Results and conclusions

The core of this article is a list of the diseases and related traits or phenotypes the causal gene of which was identified in the rat (Table 1). The genes identified by forward genetic methods or, in a few instances, by direct molecular characterization are labeled by asterisks (see legend to table). Also listed are the phenotypes uncovered by reverse genetics methods, either by ENU-mutagenesis followed by selection of the desired mutated gene (these genes are labeled by the symbol ^ENU^), or by targeted gene editing (these genes are labeled by ^T^).

**Table 1:**
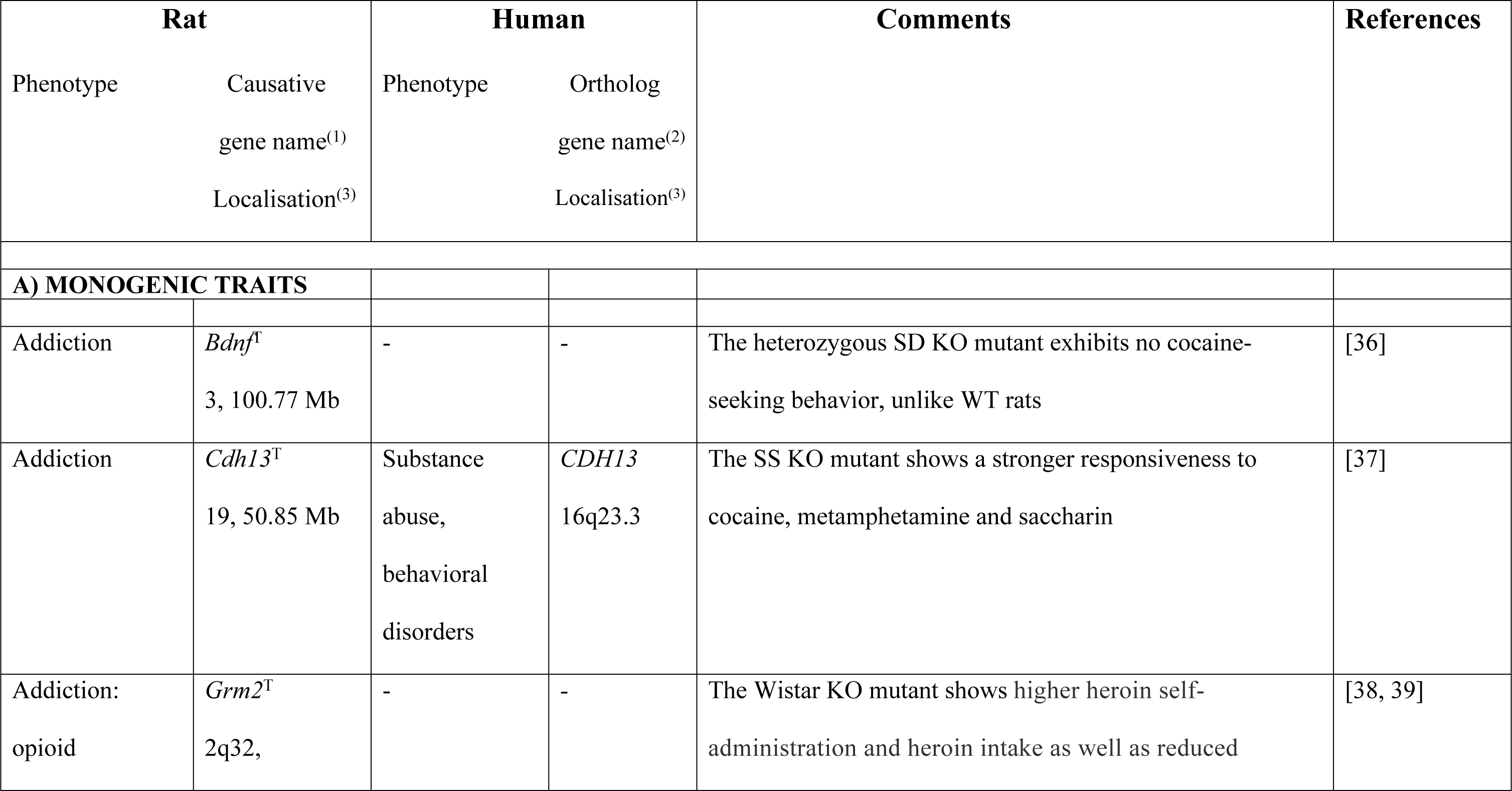
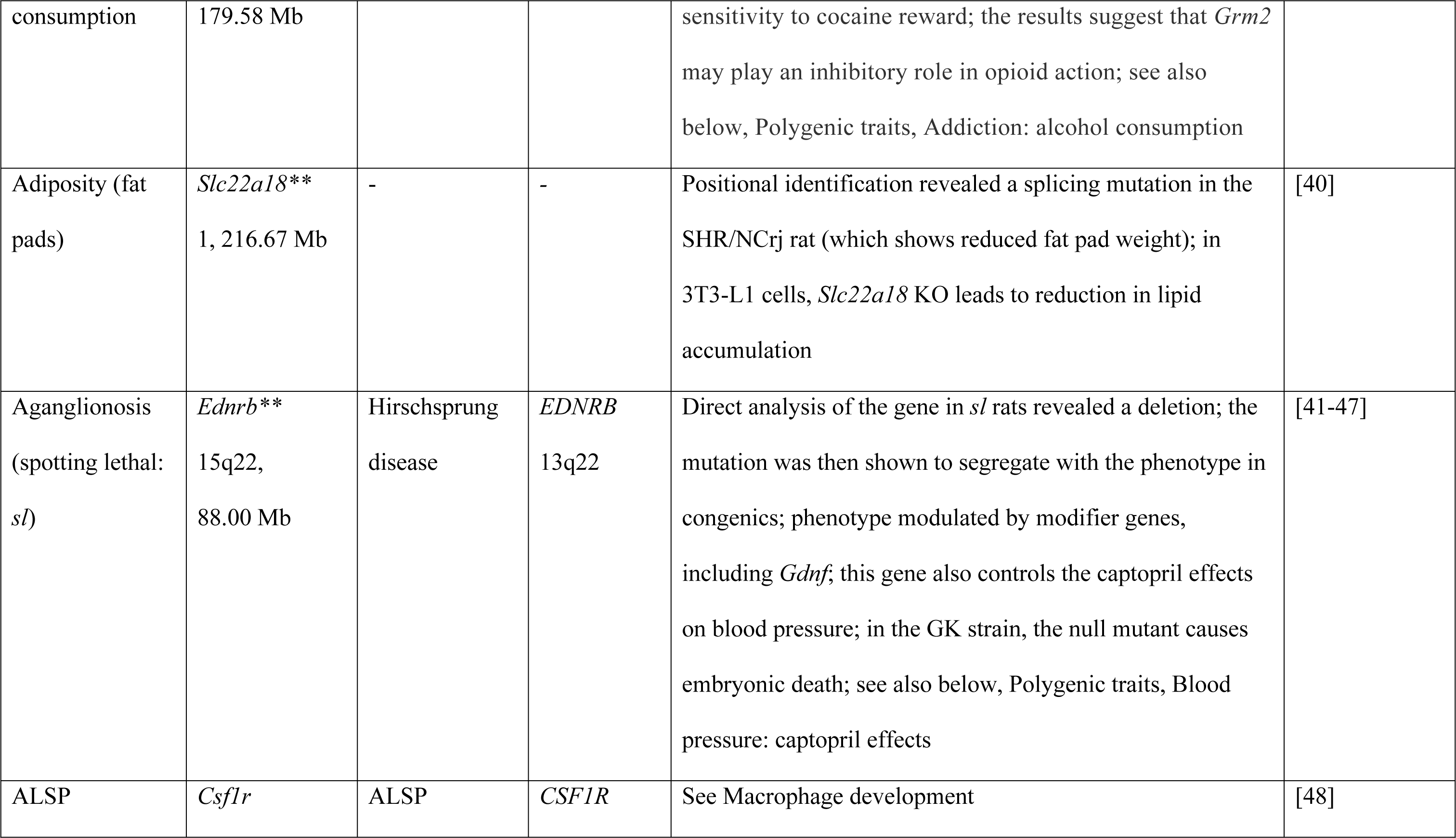
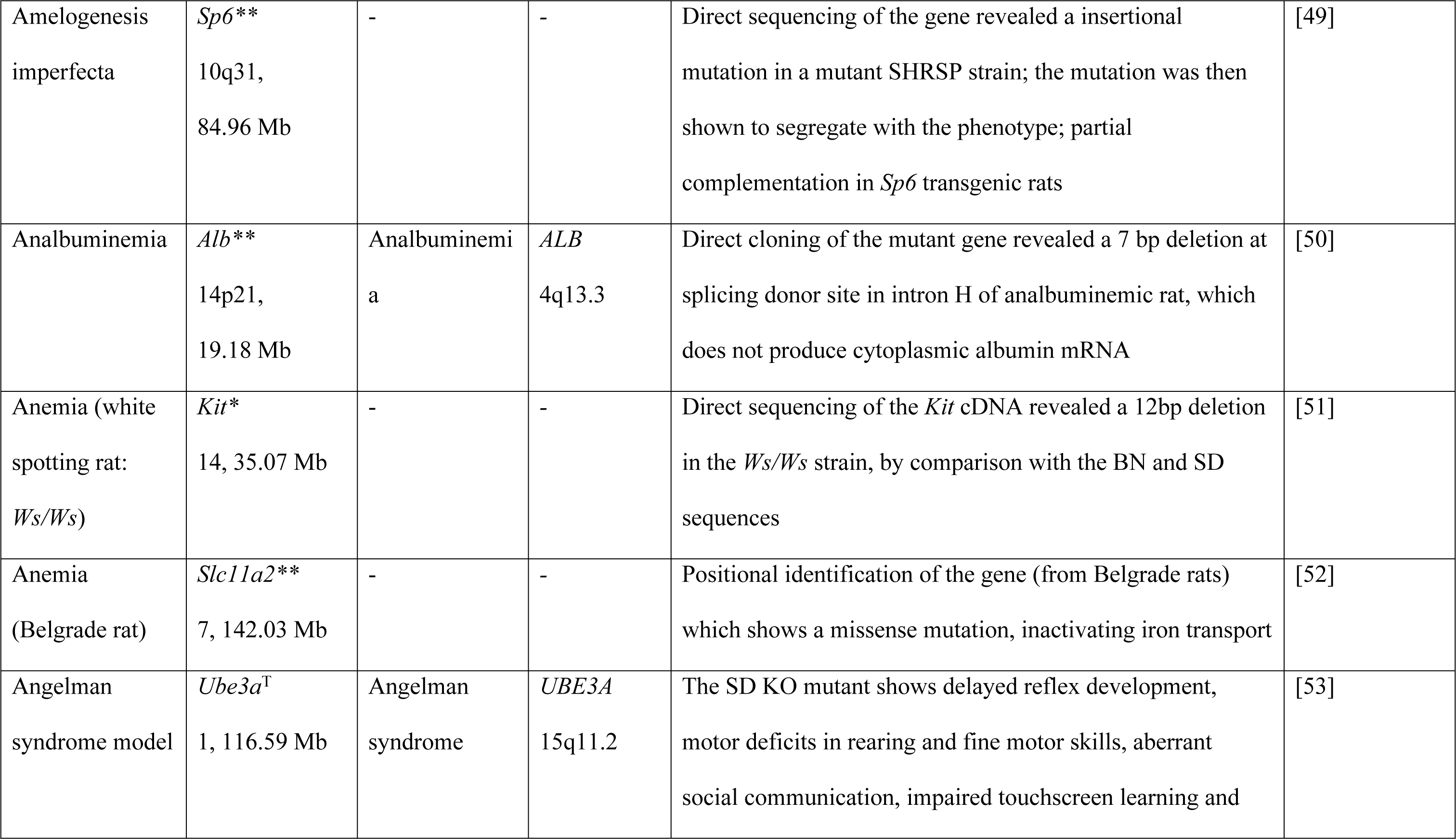
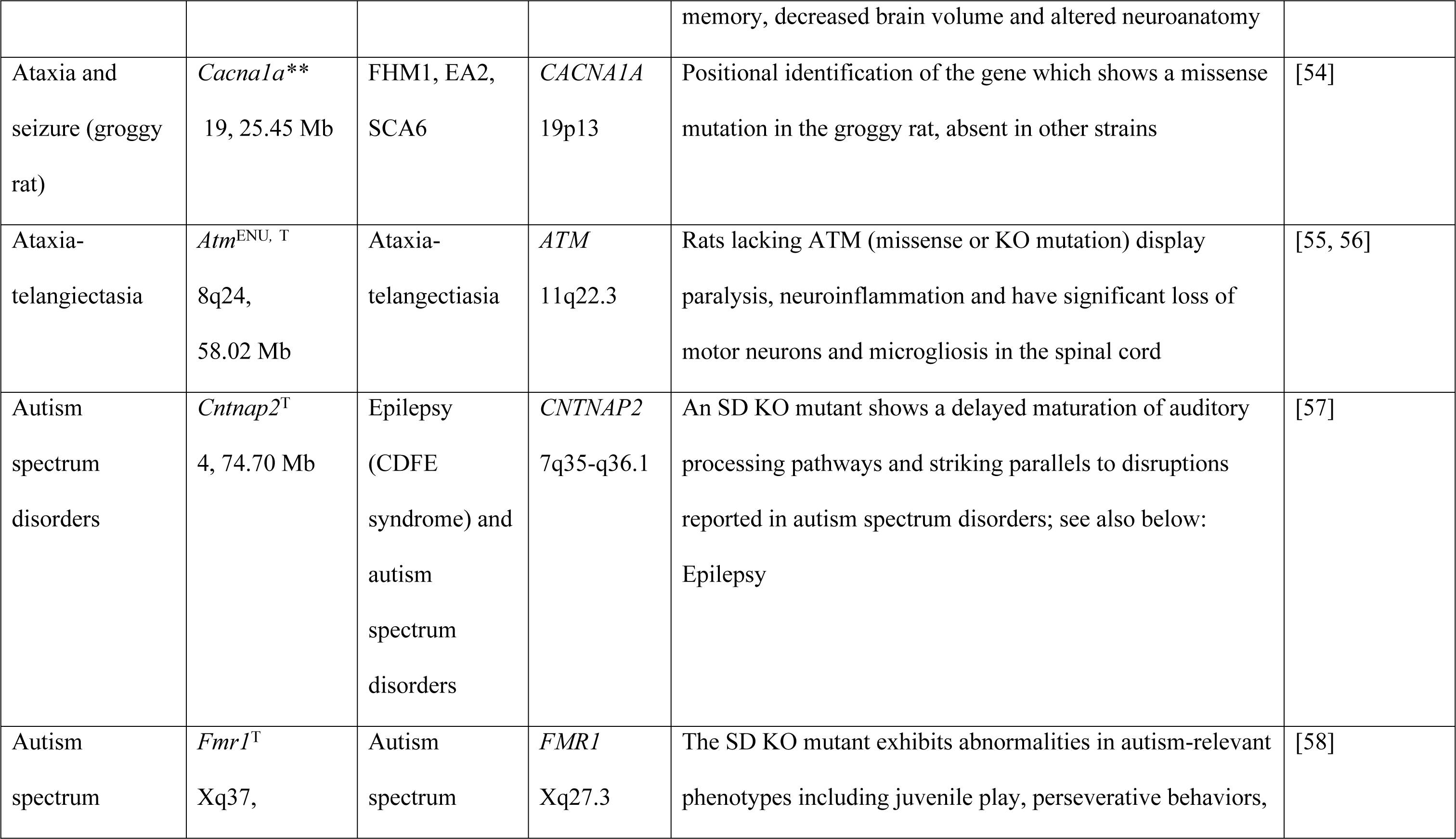
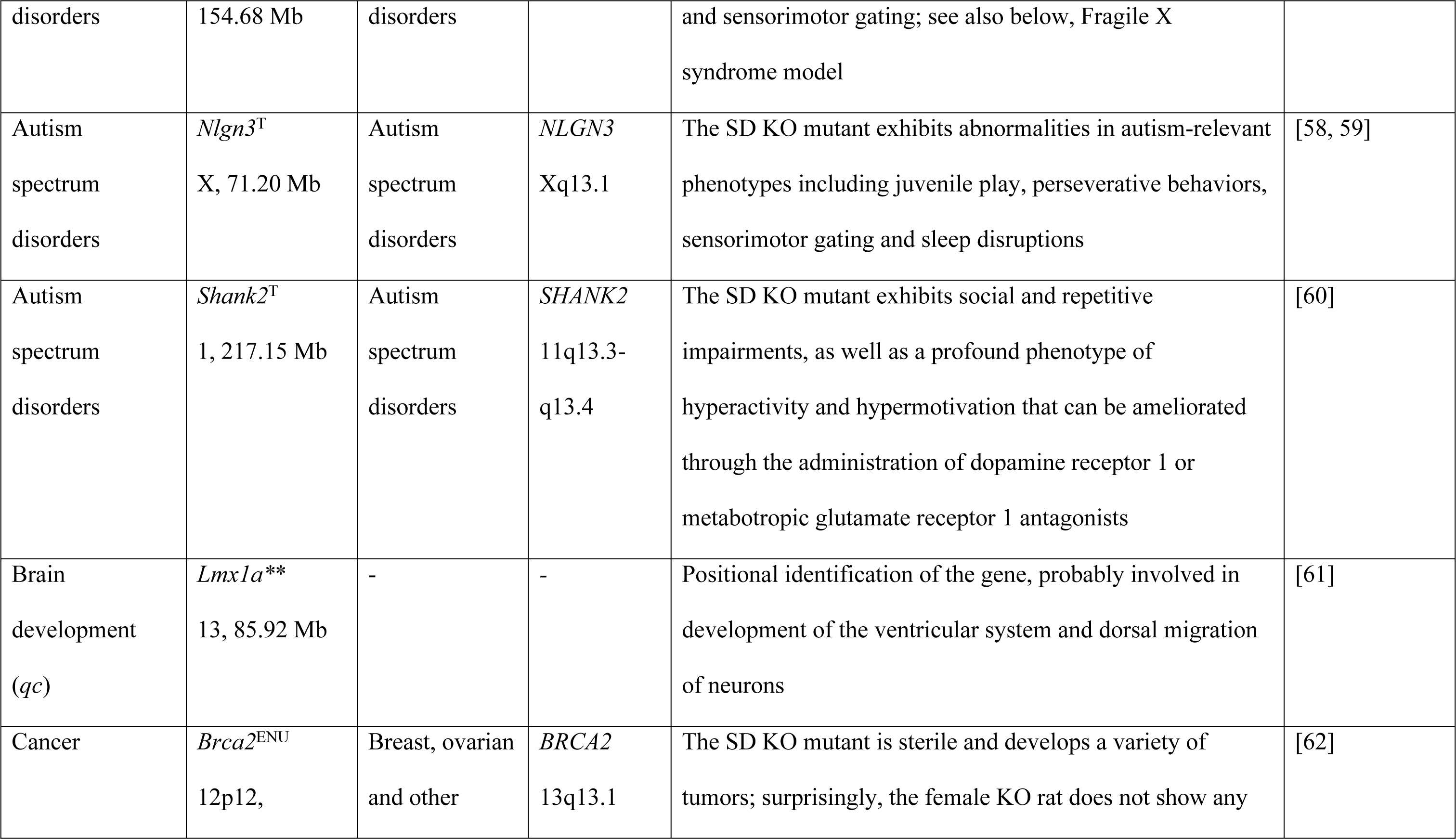
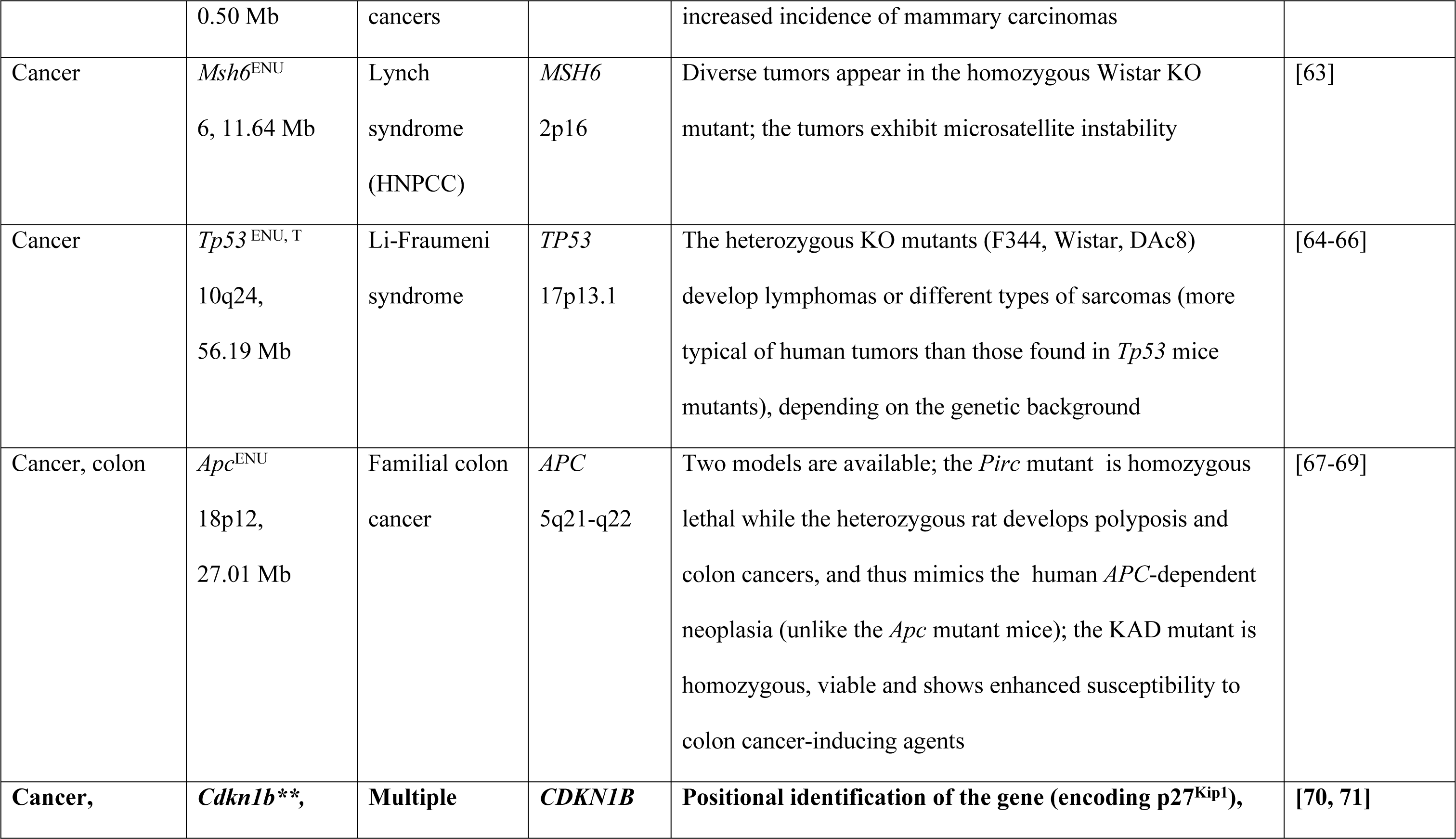
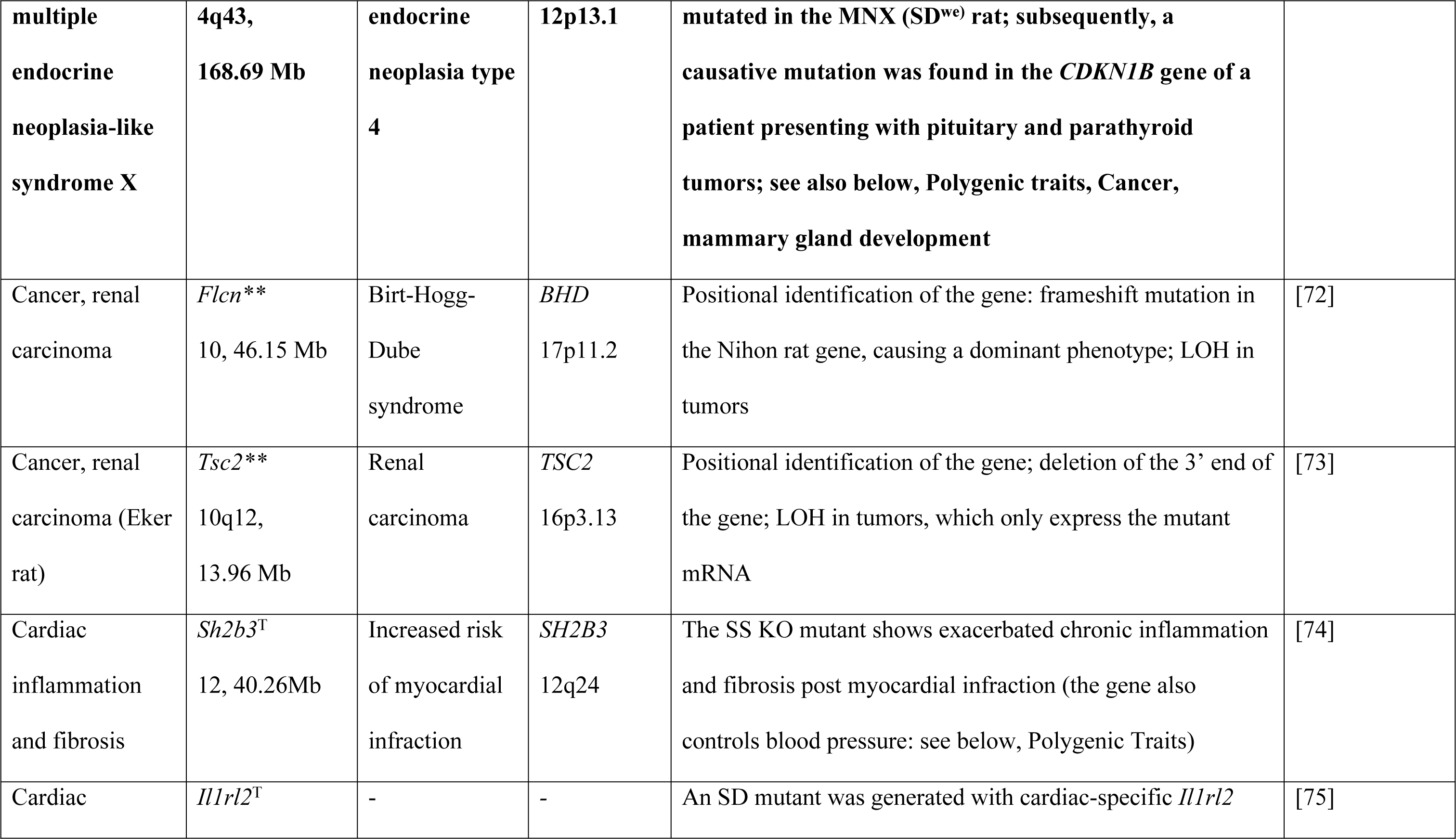
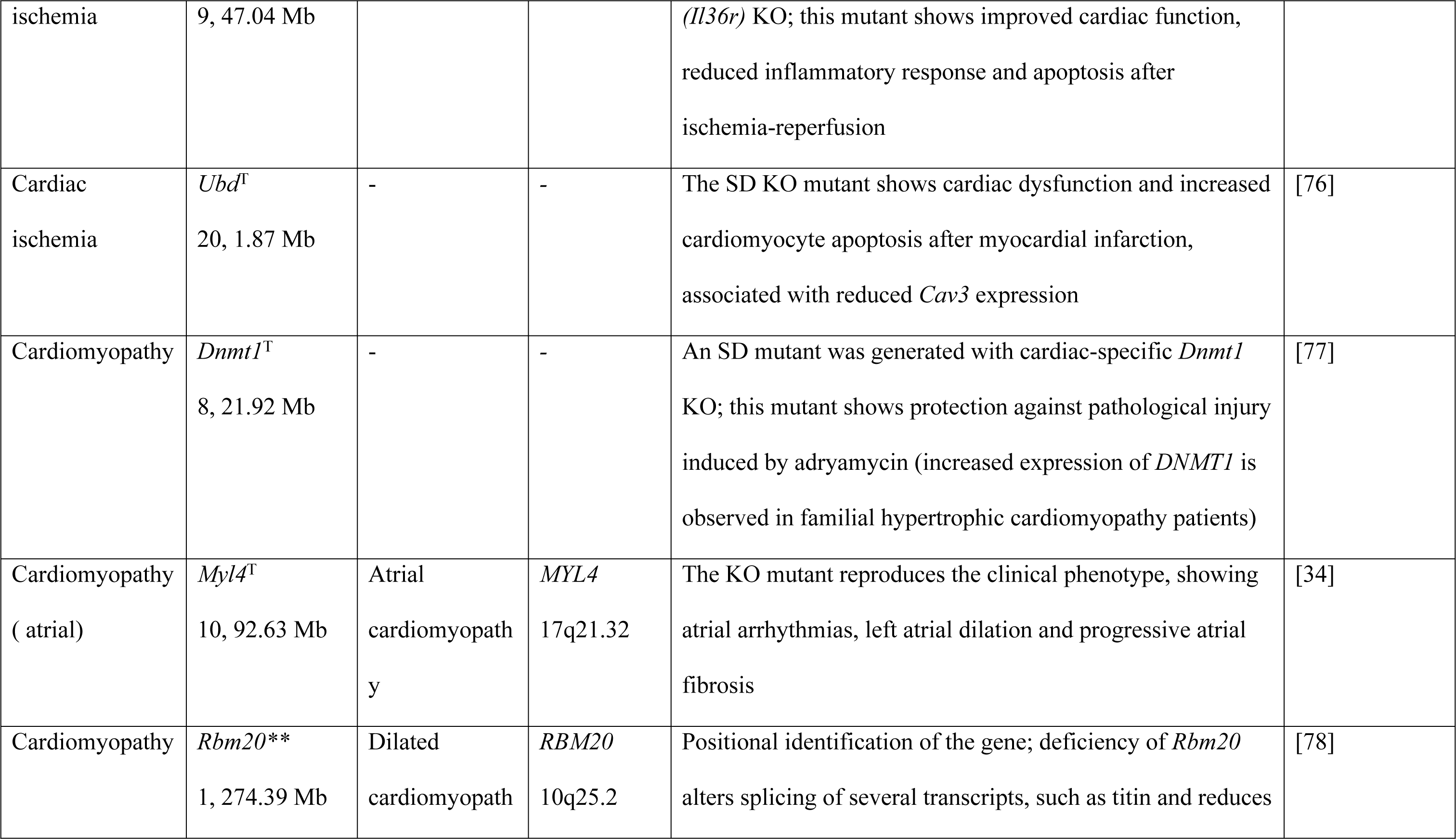
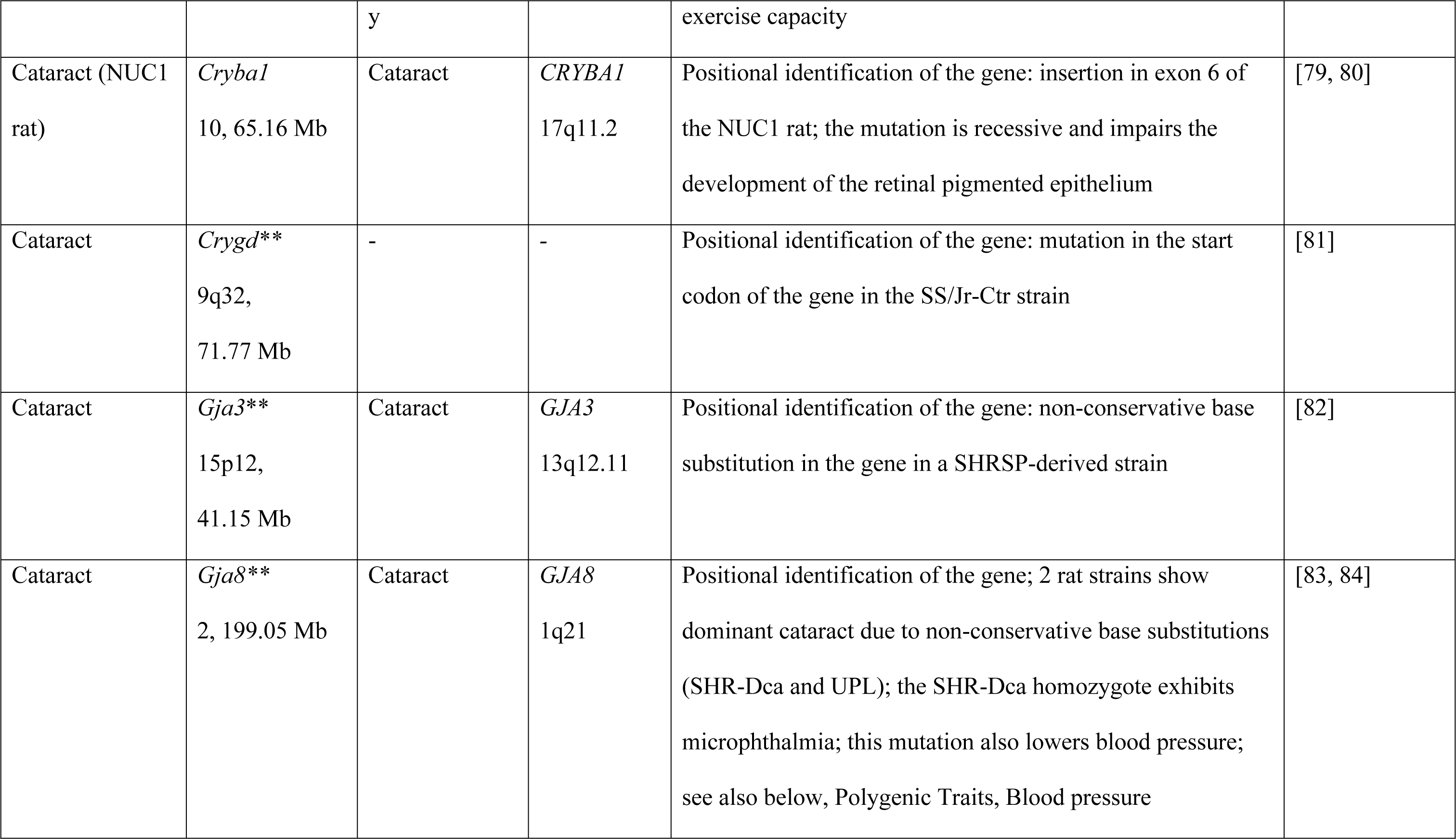
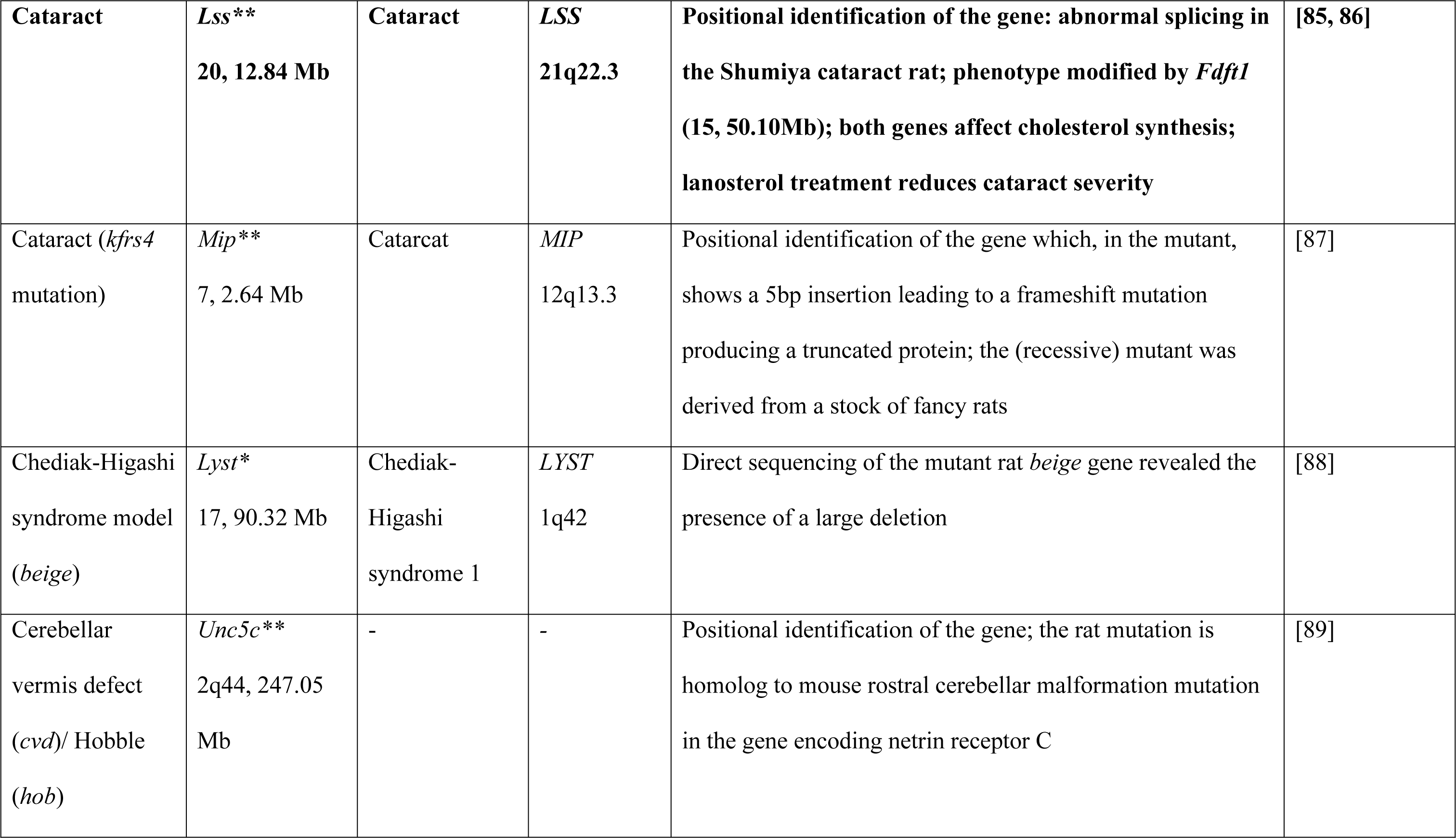
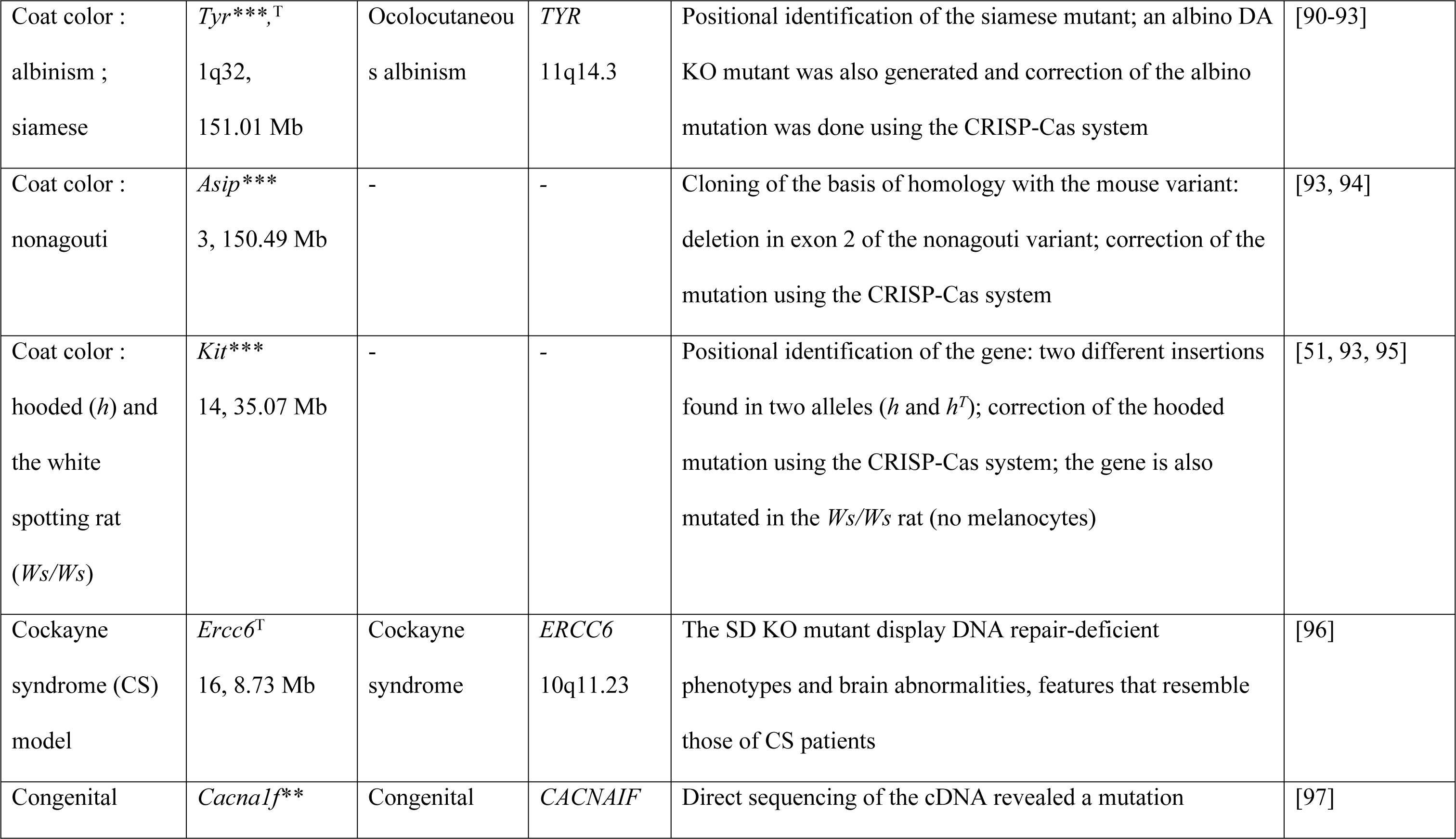
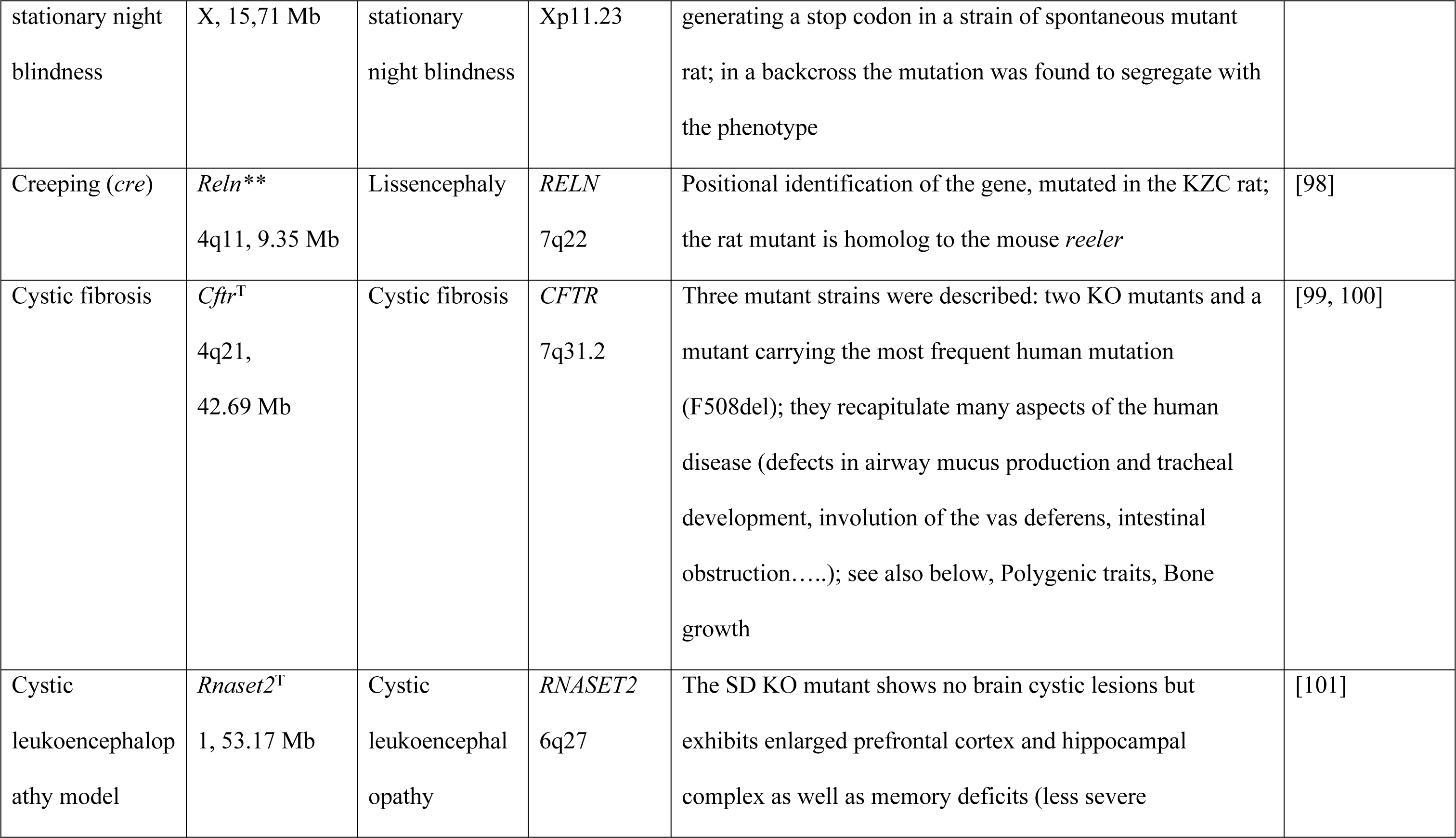
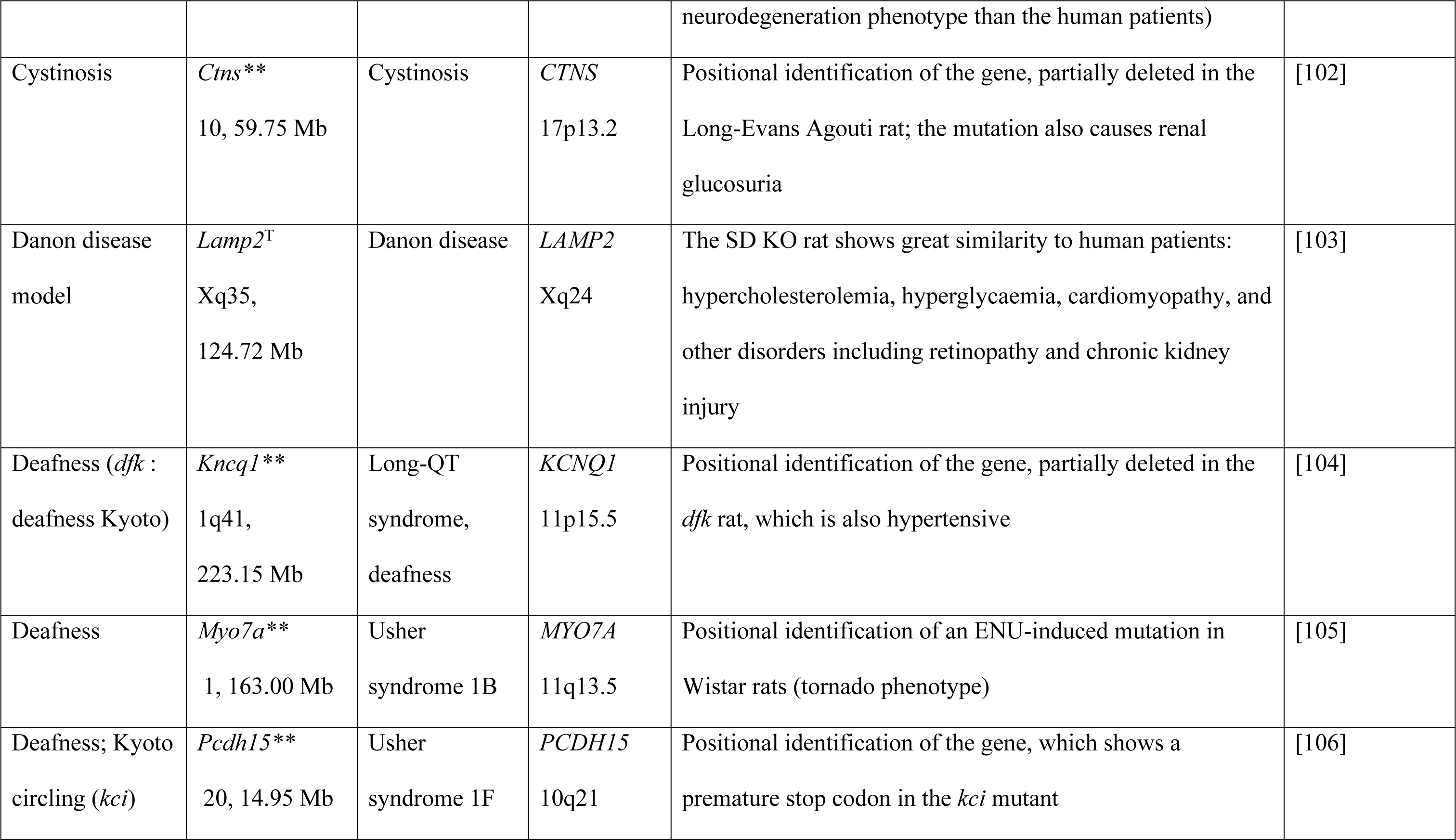
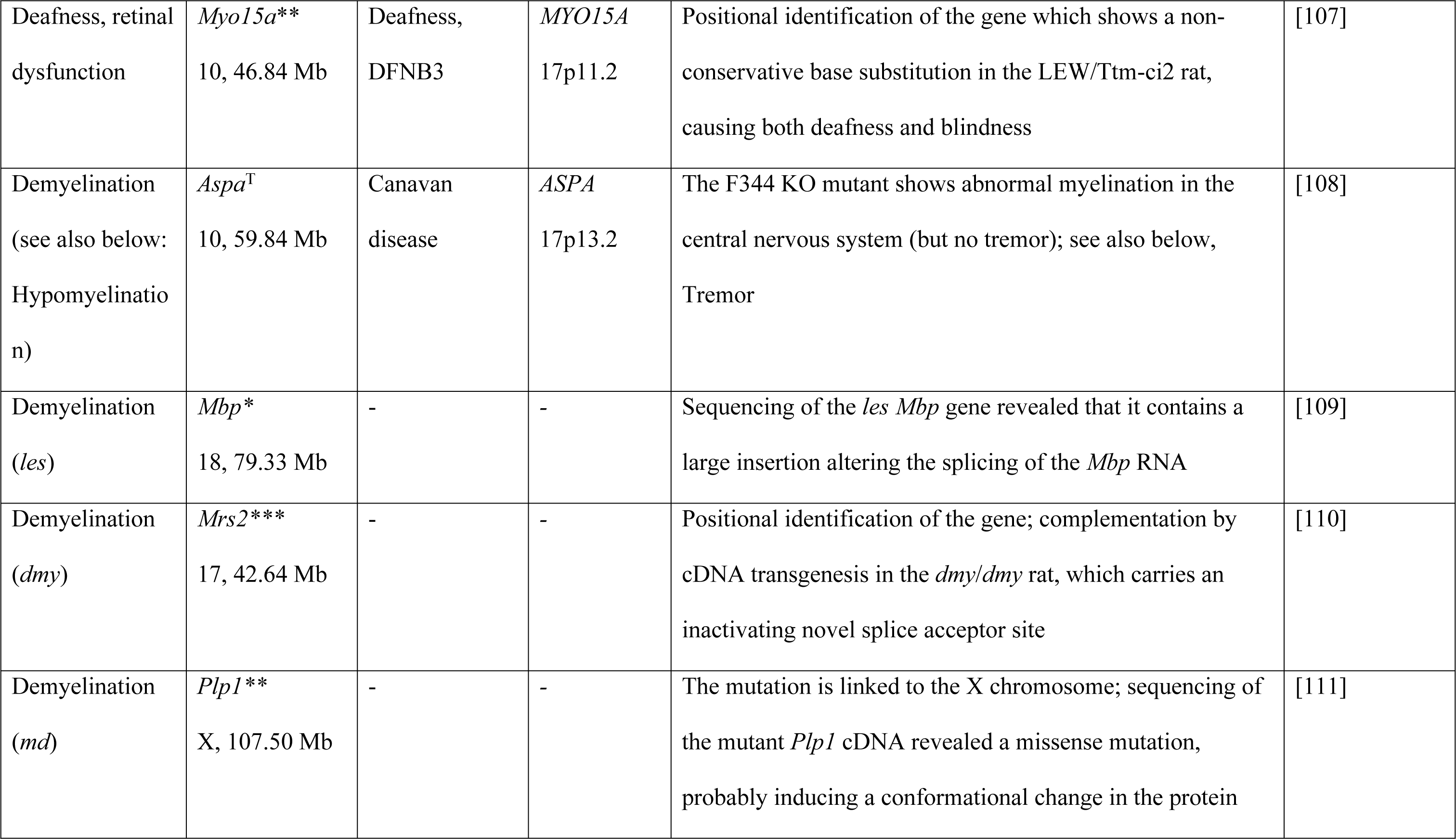
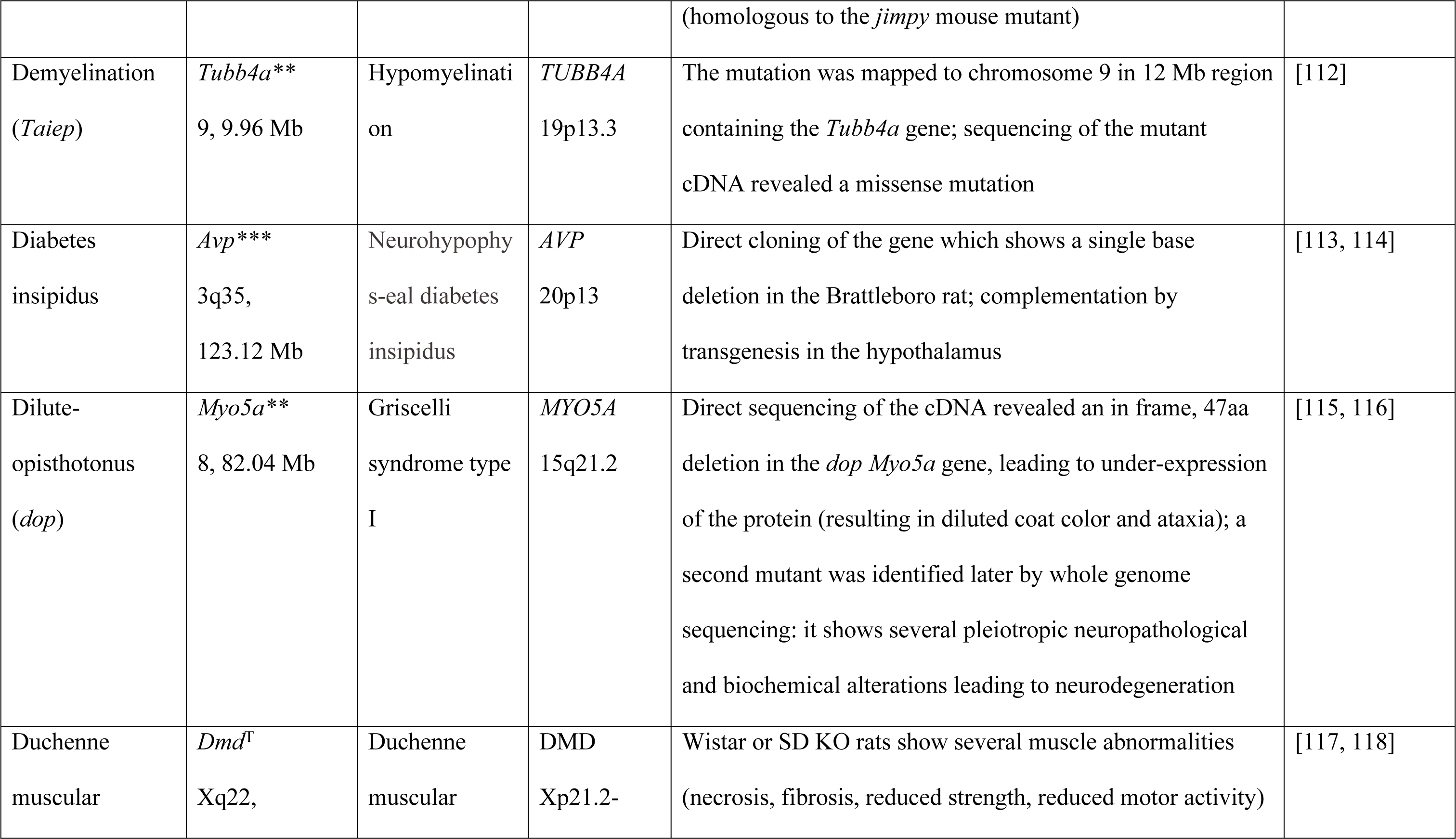
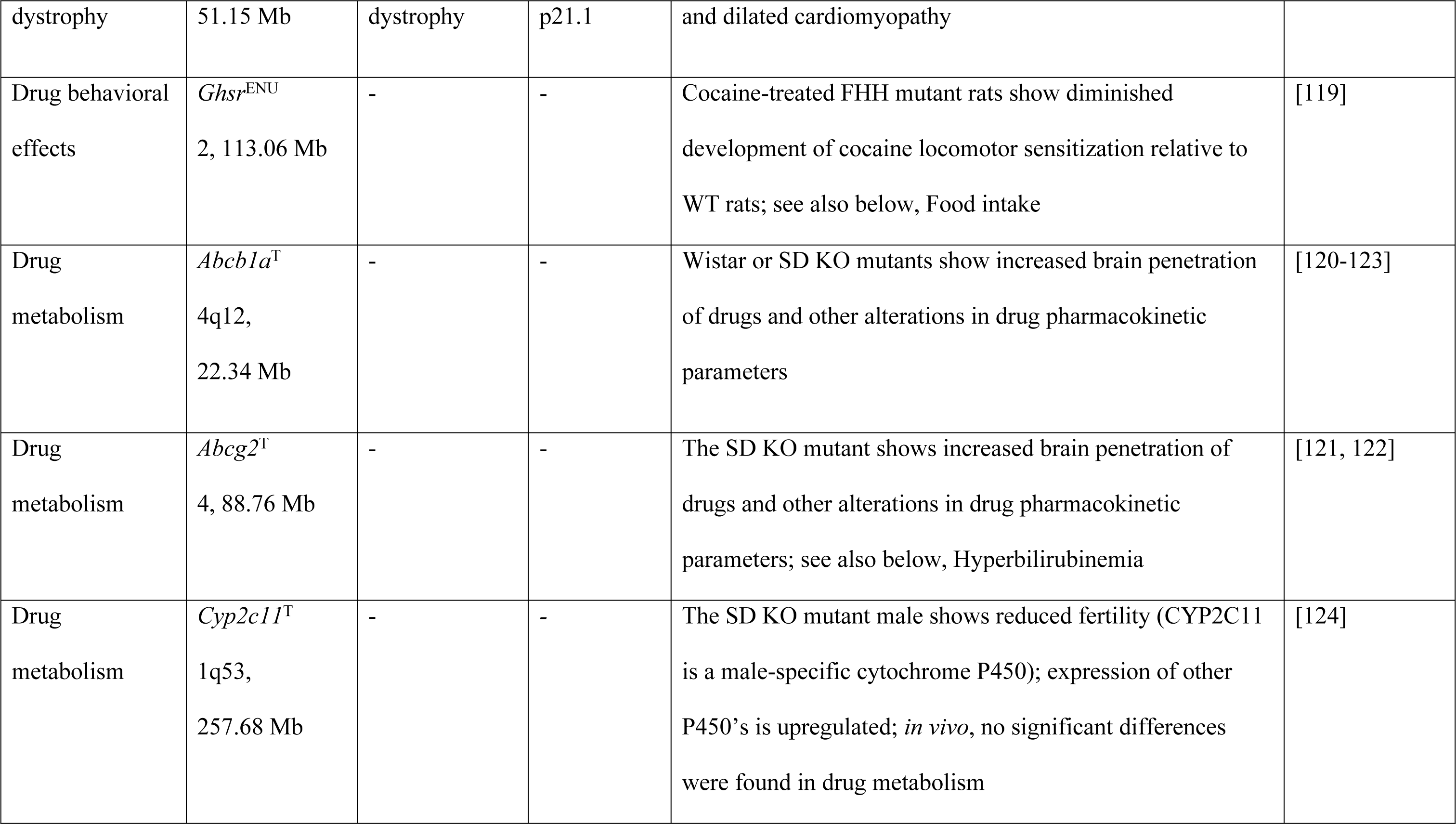
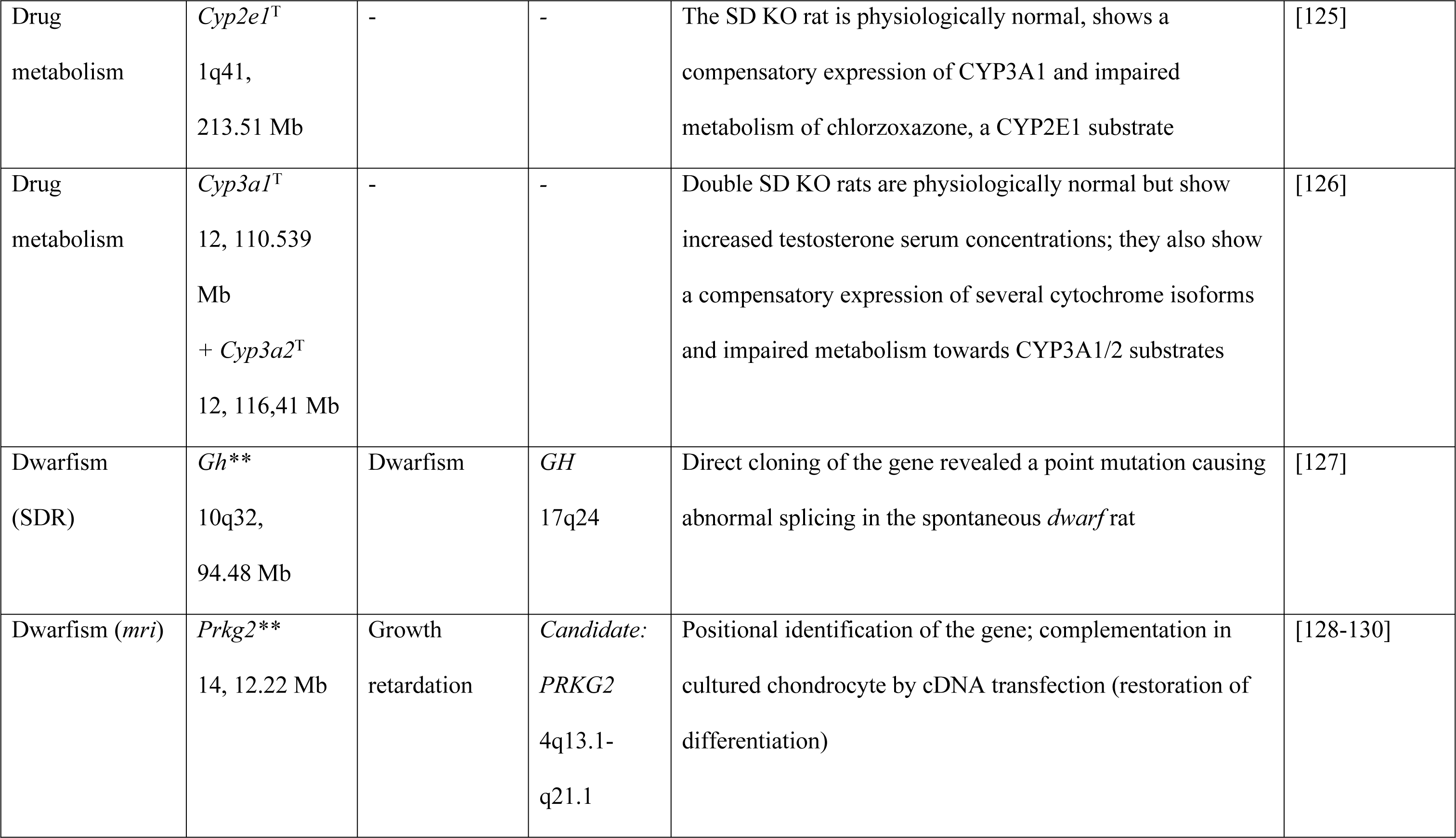
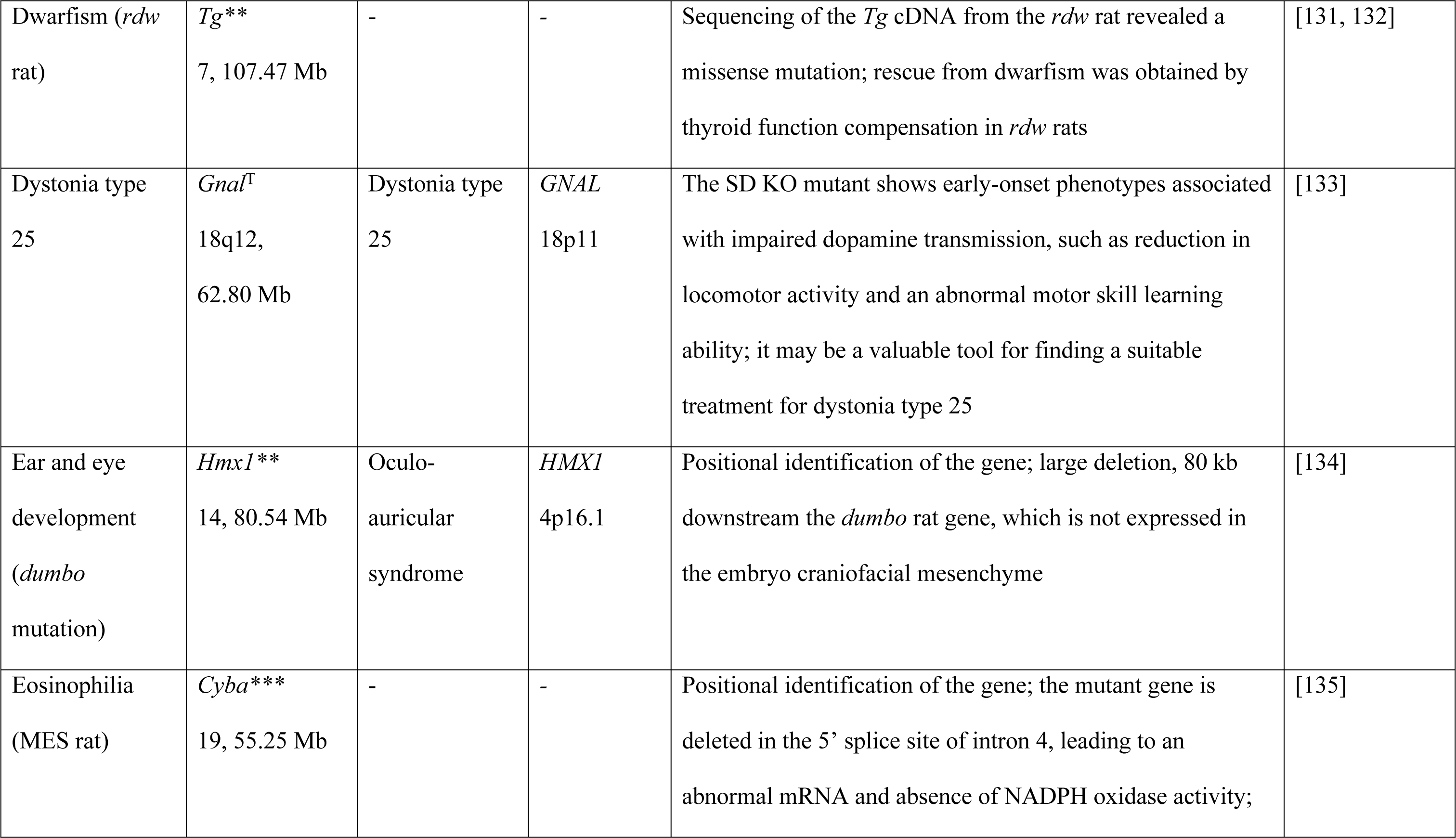
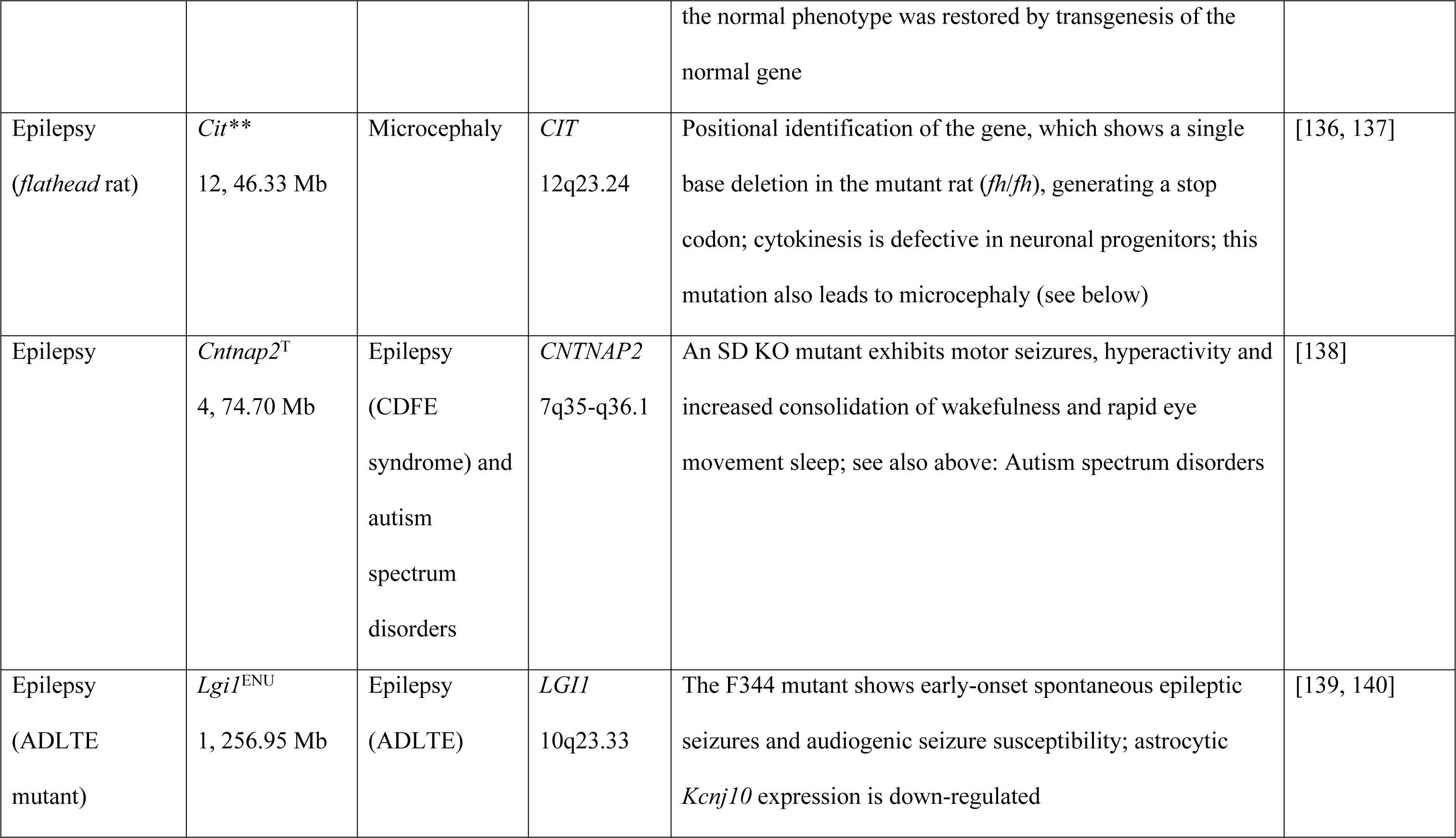
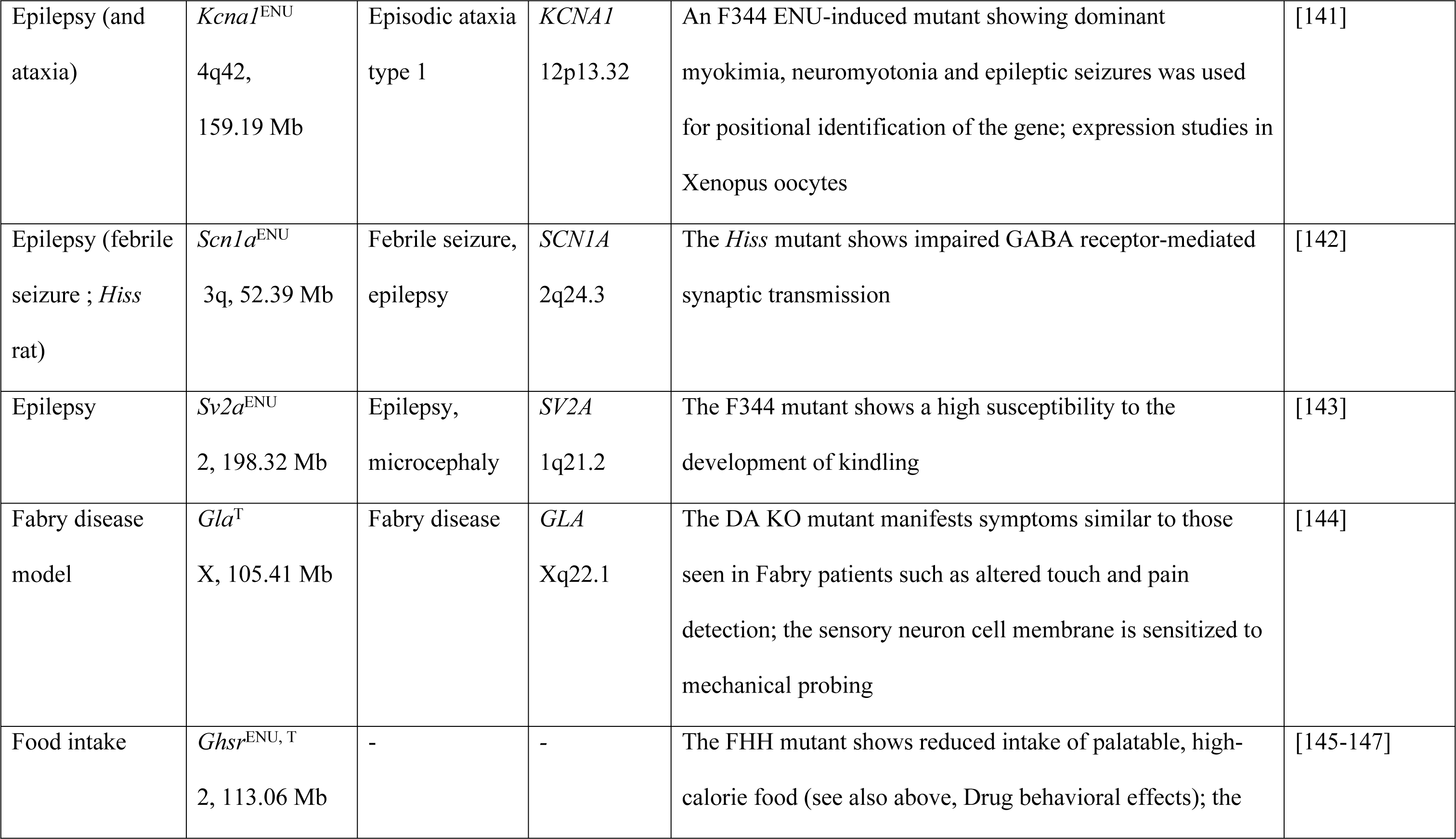
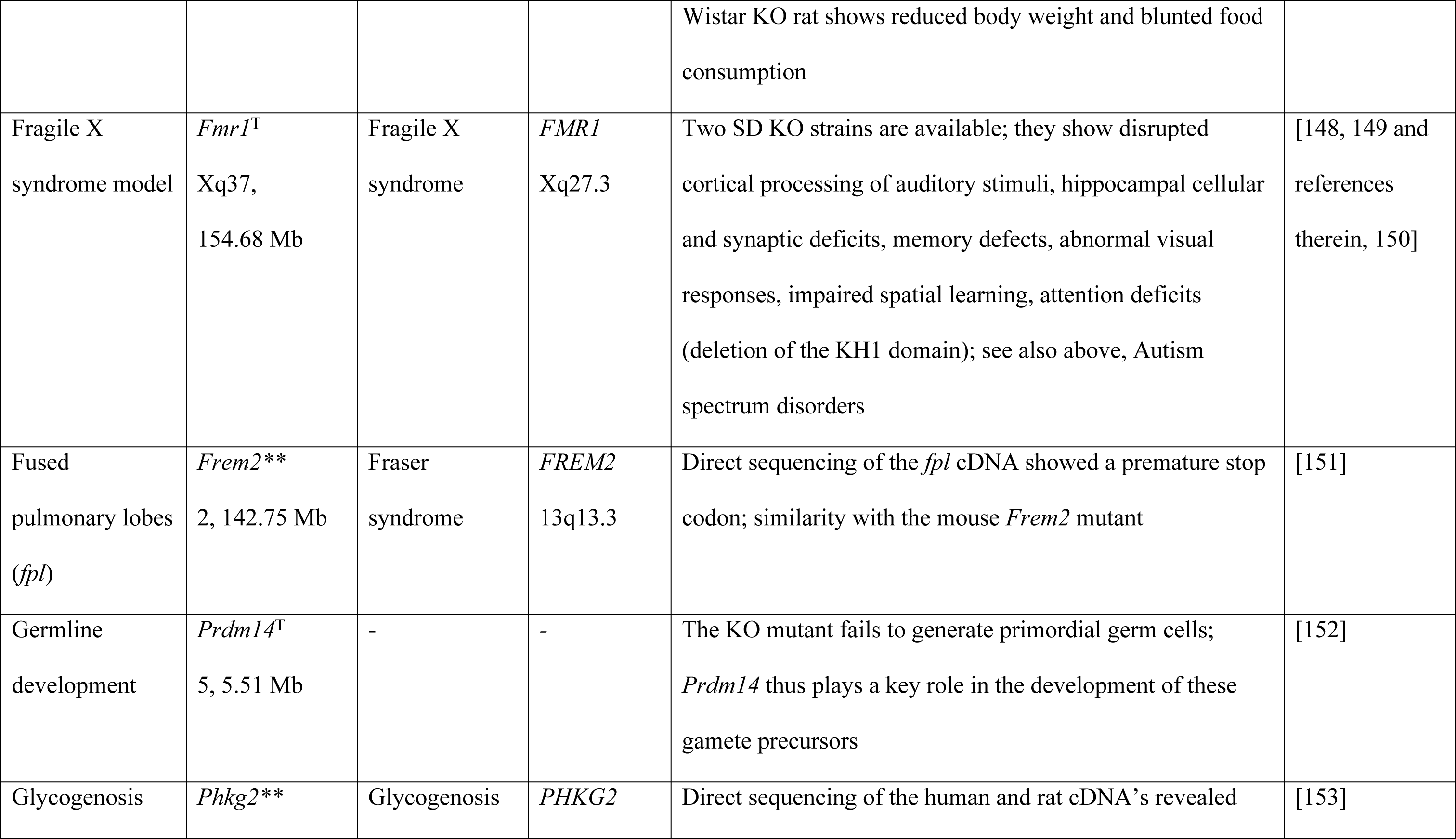
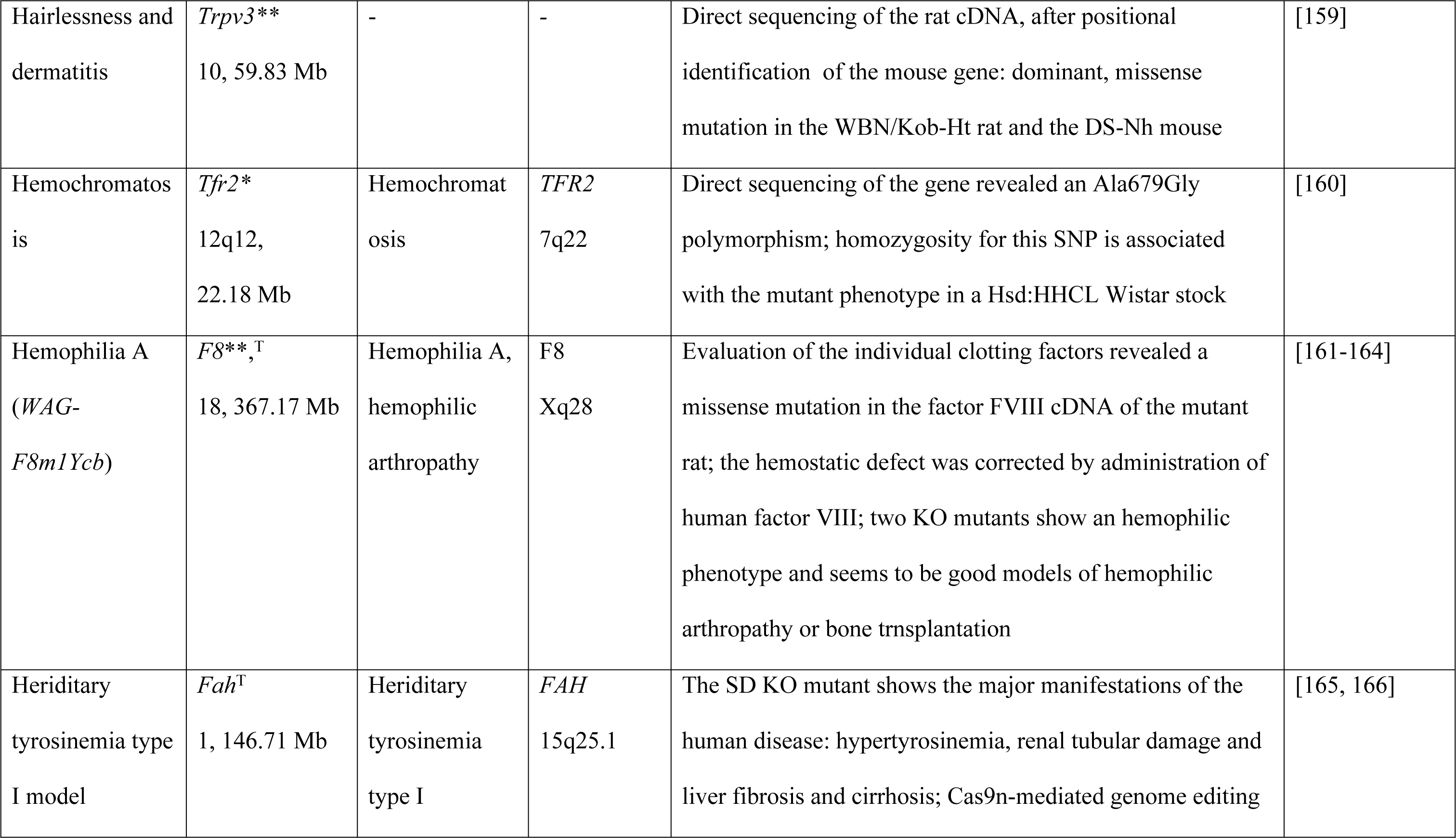
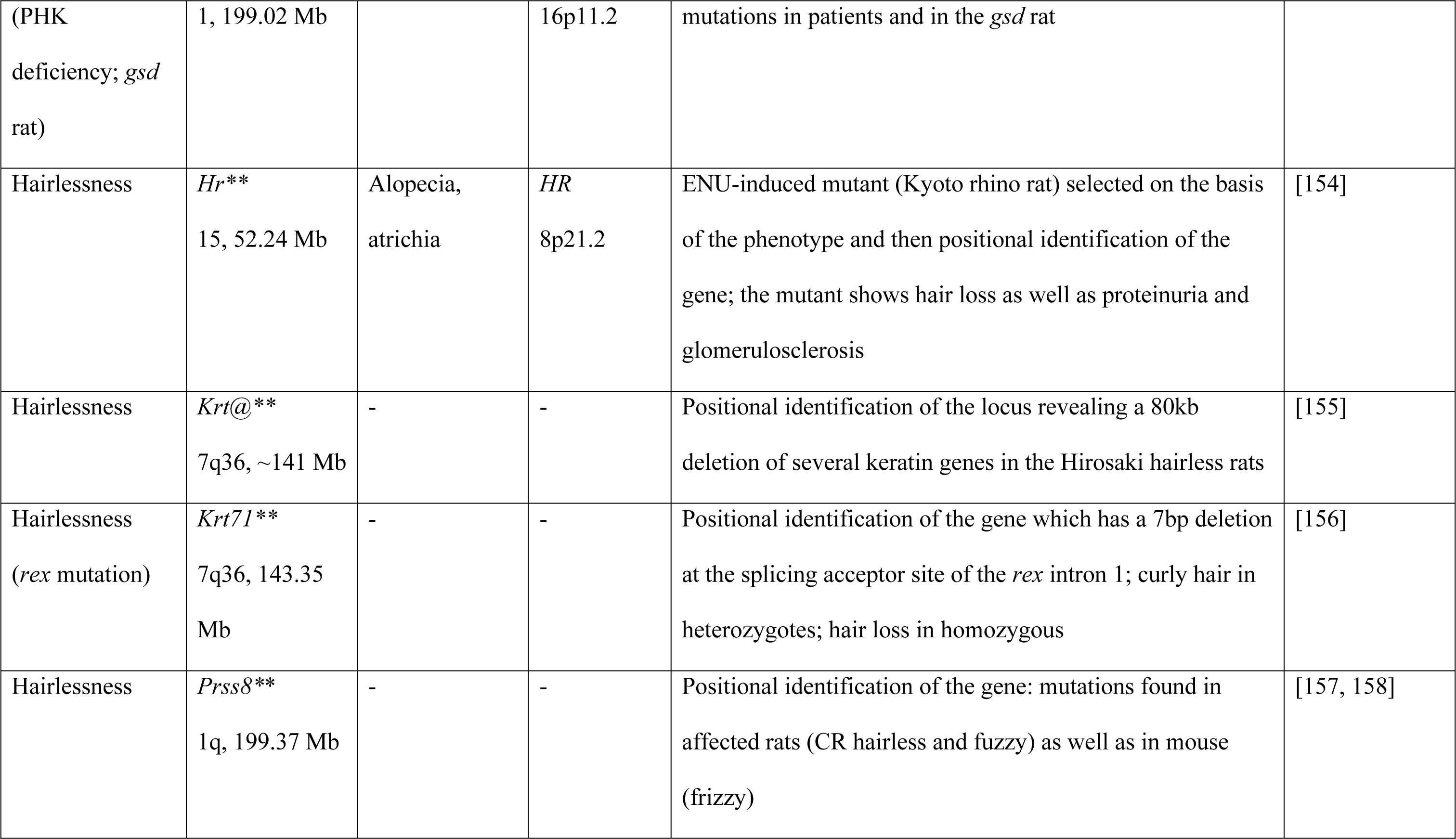
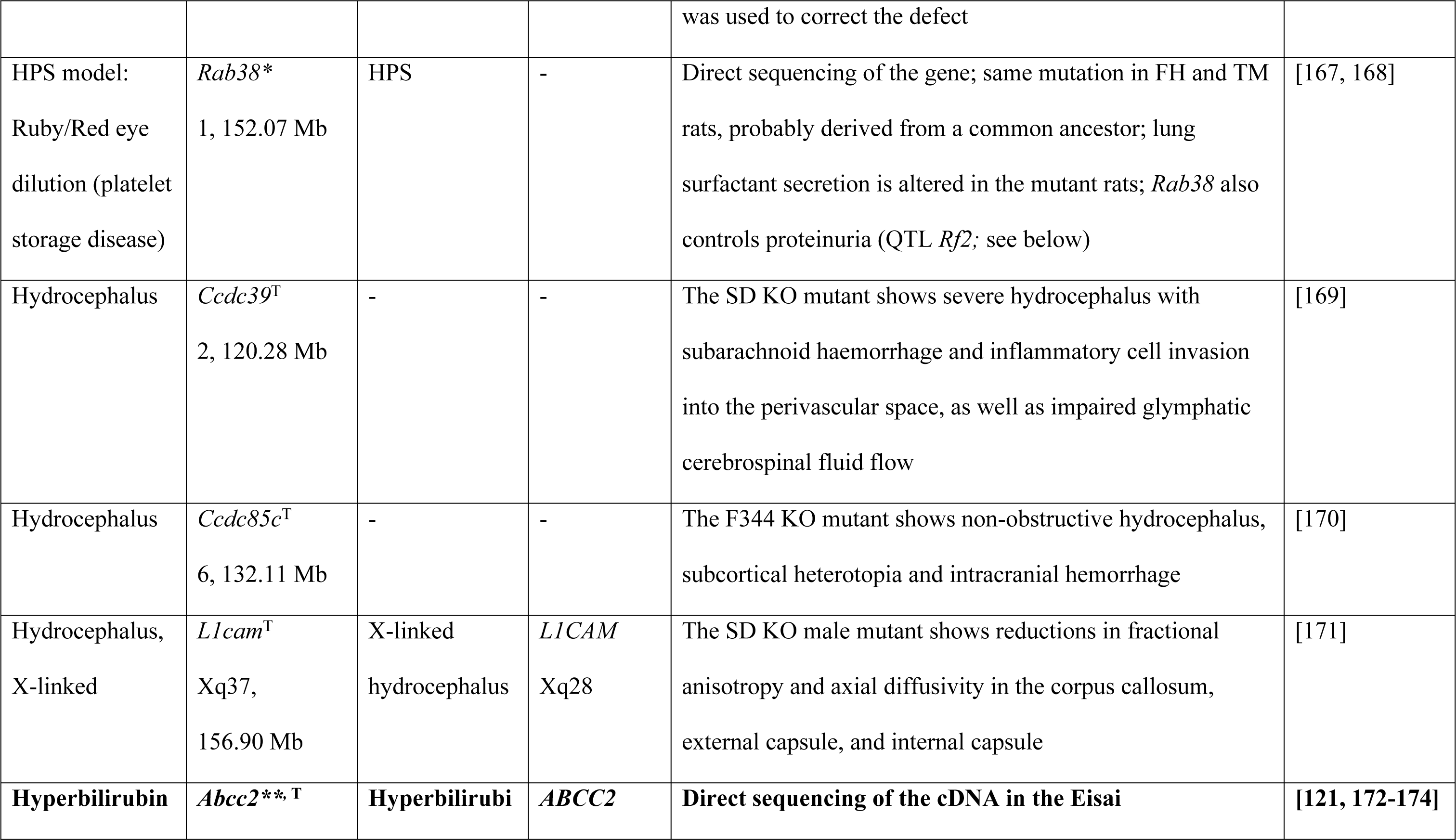
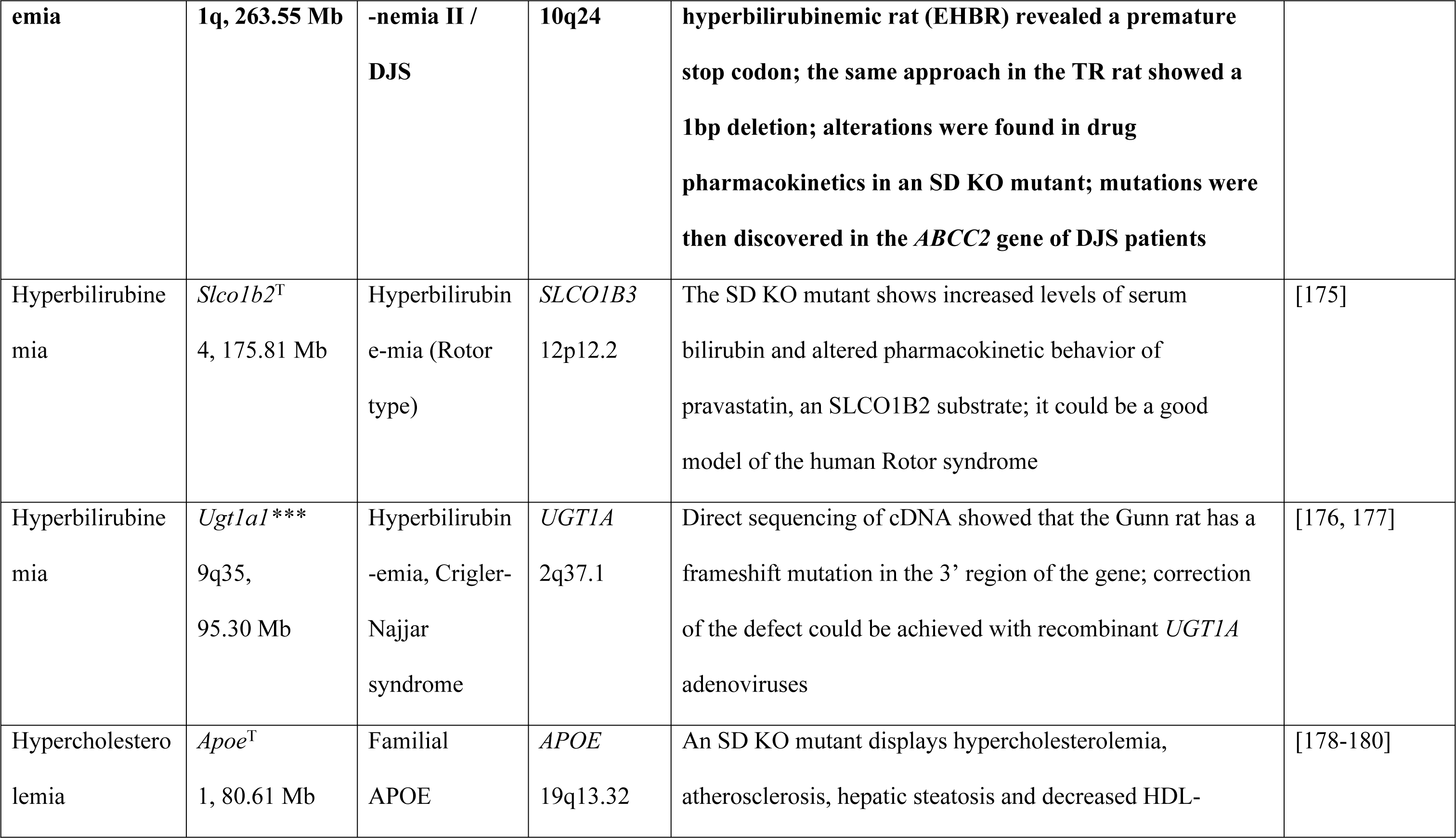
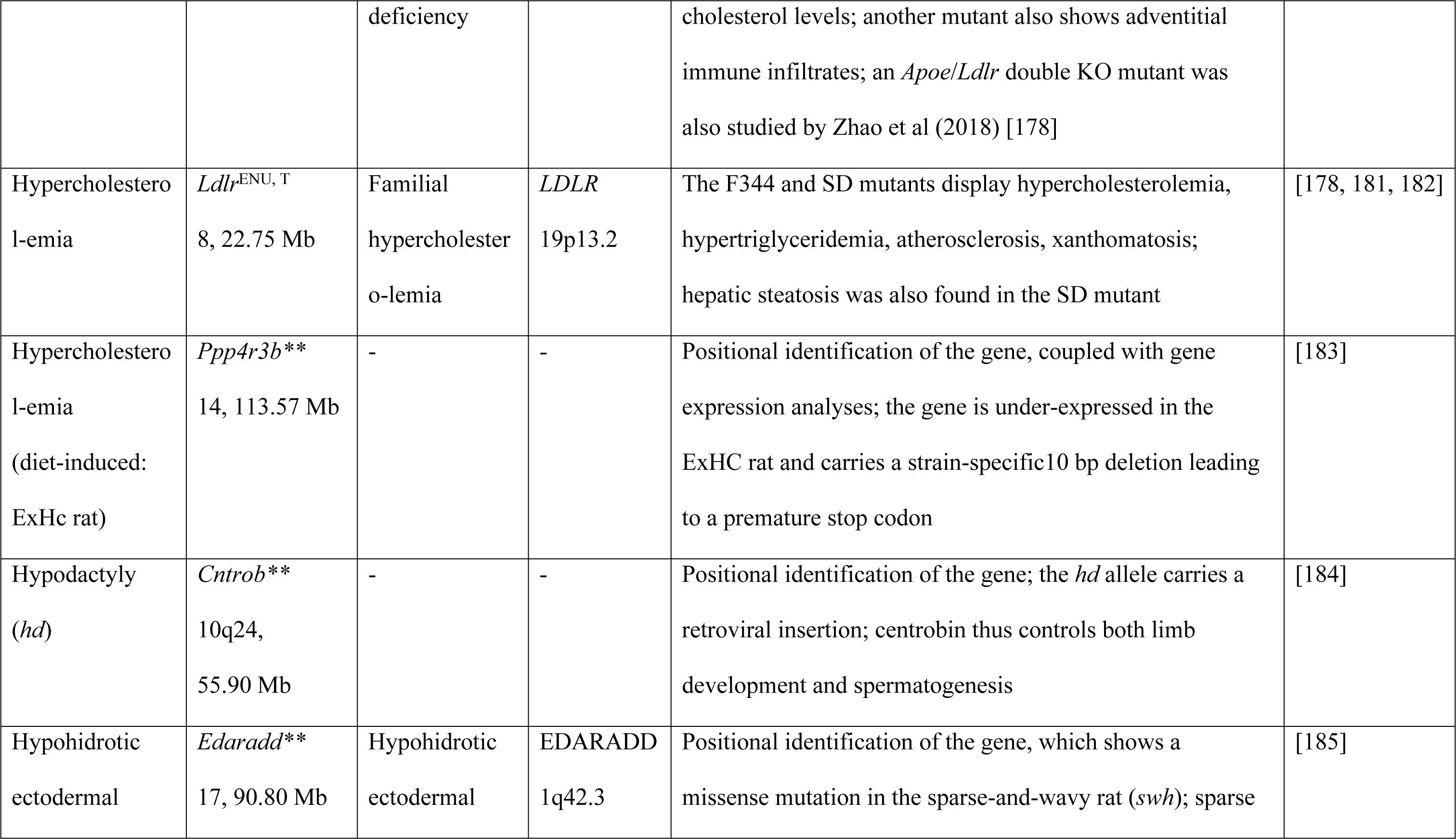
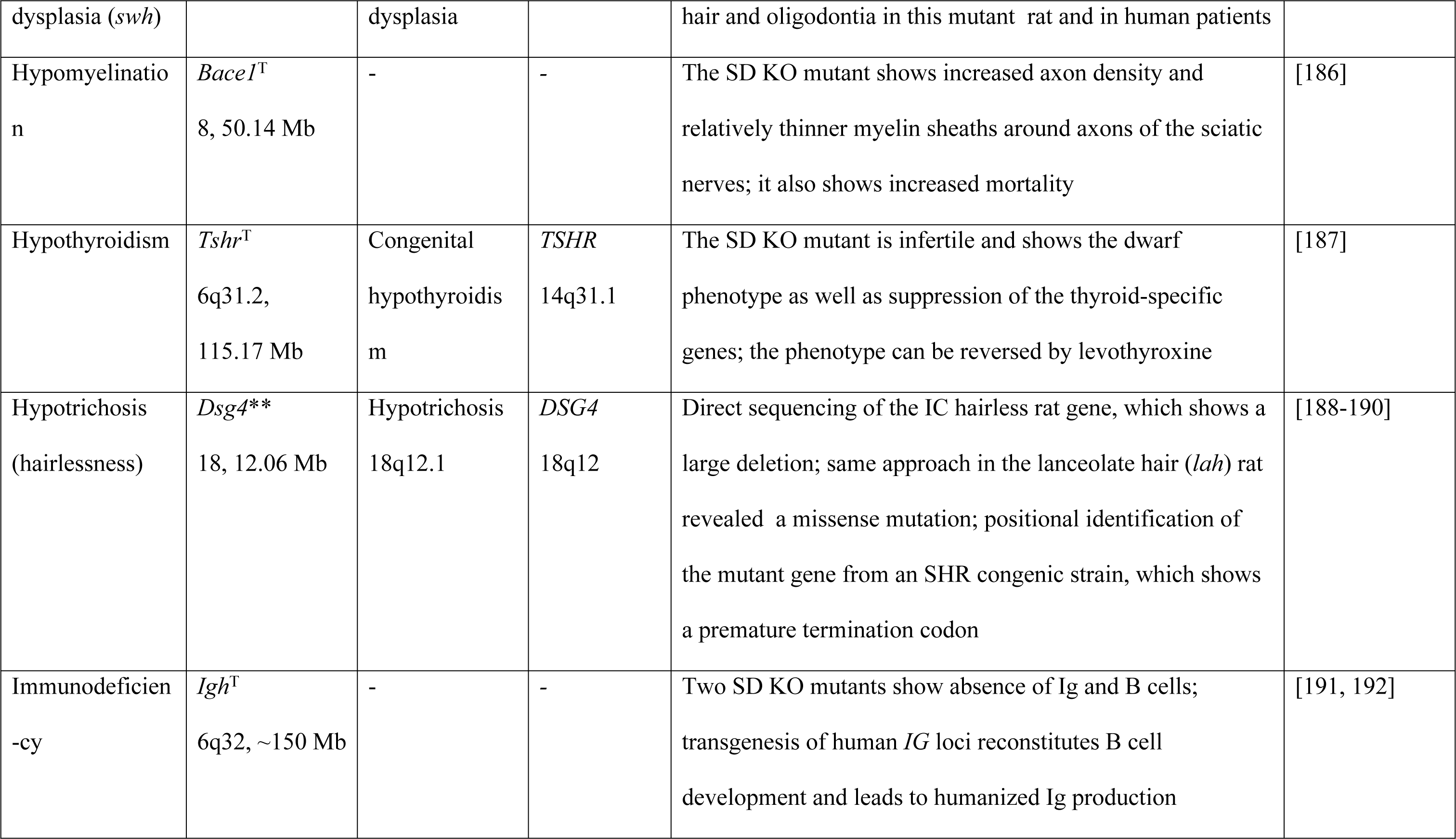
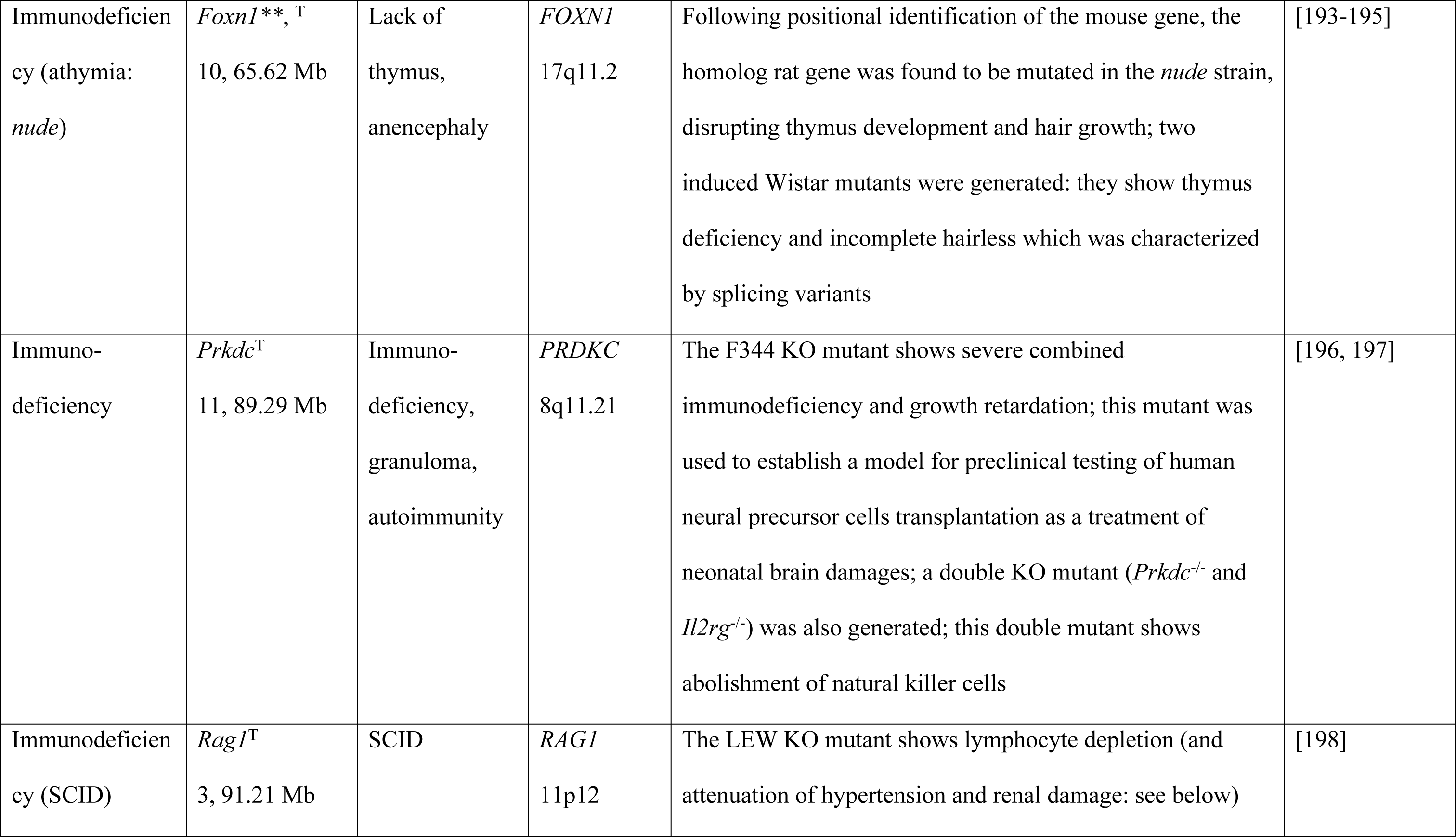
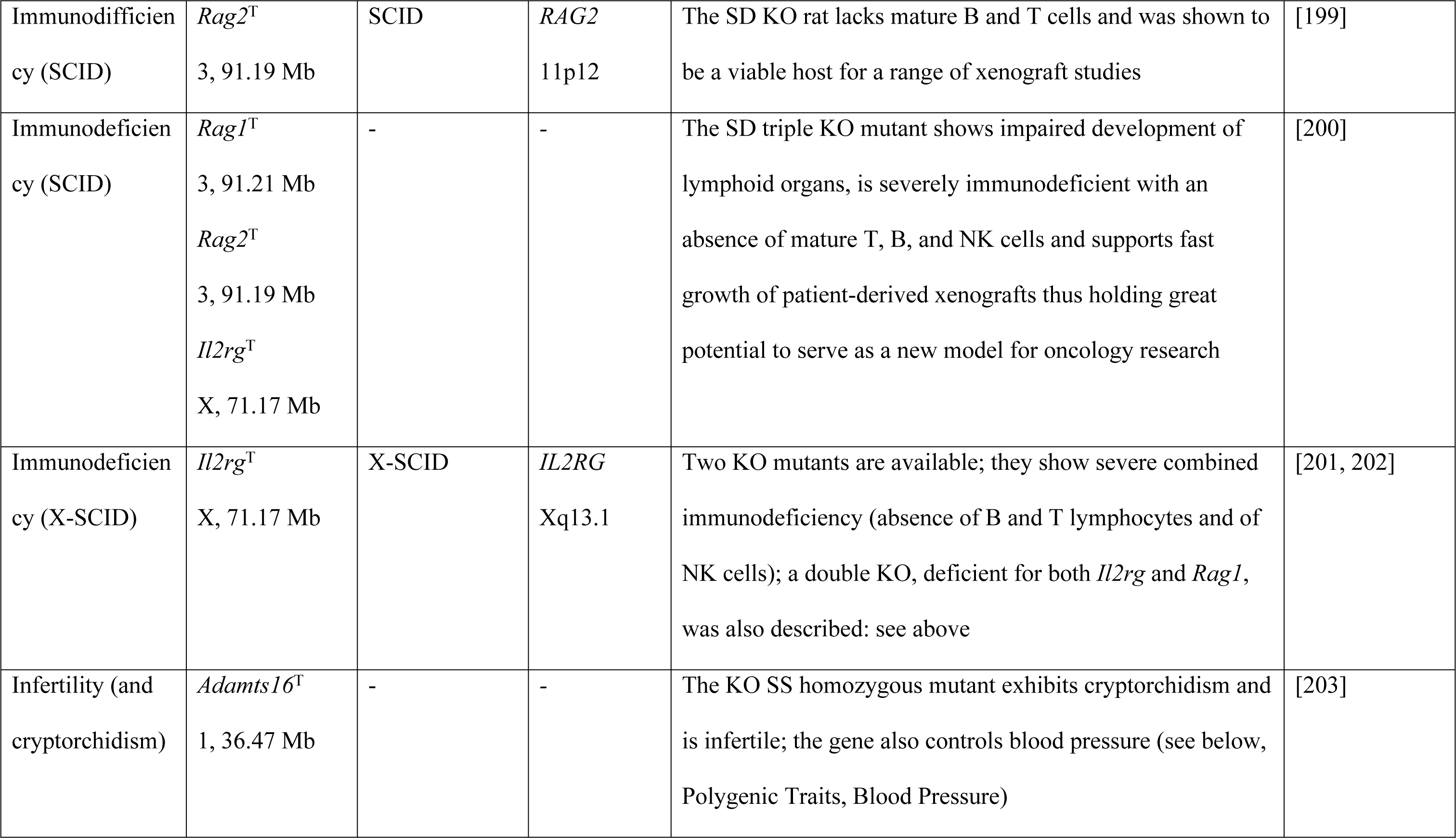
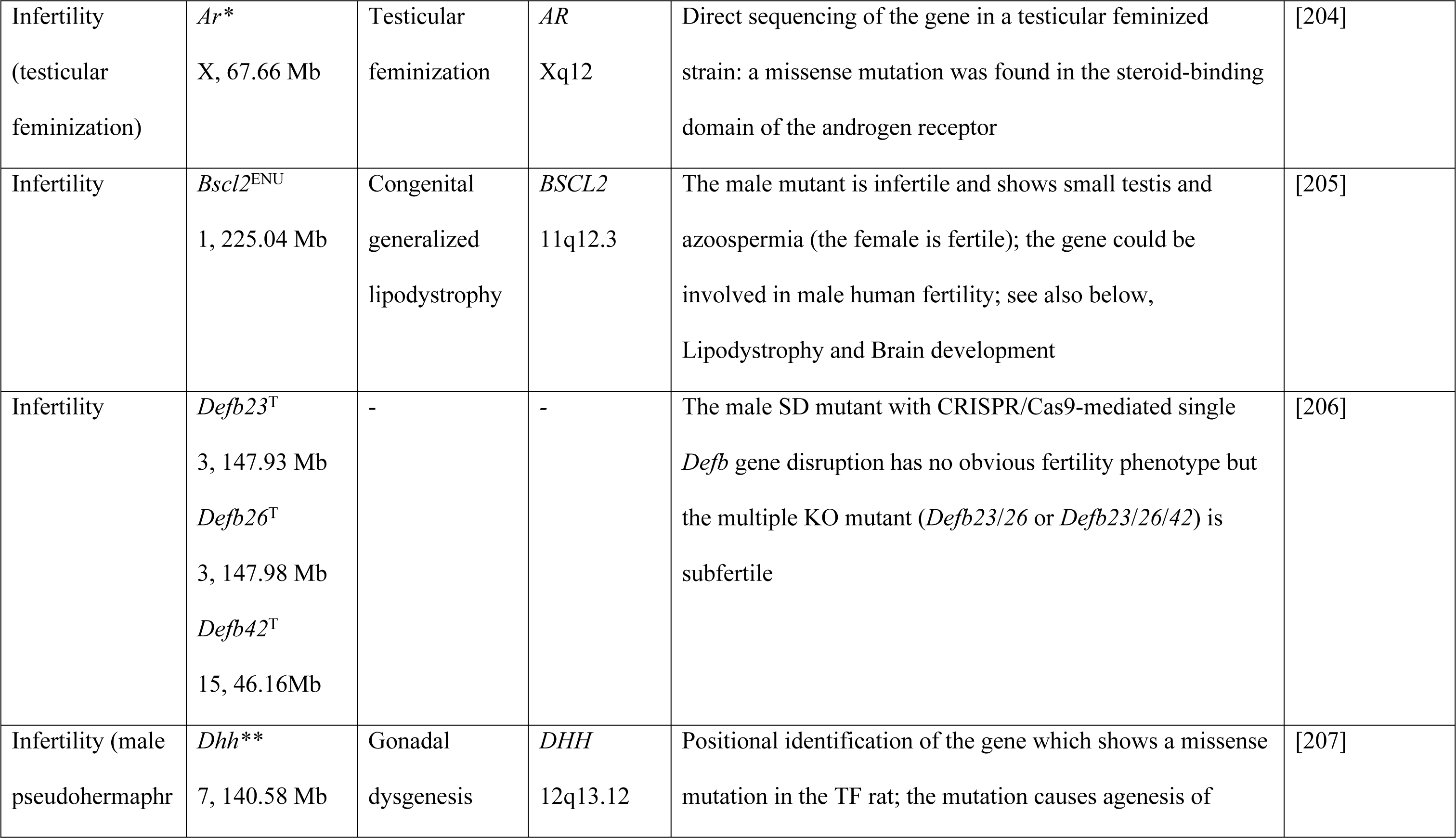
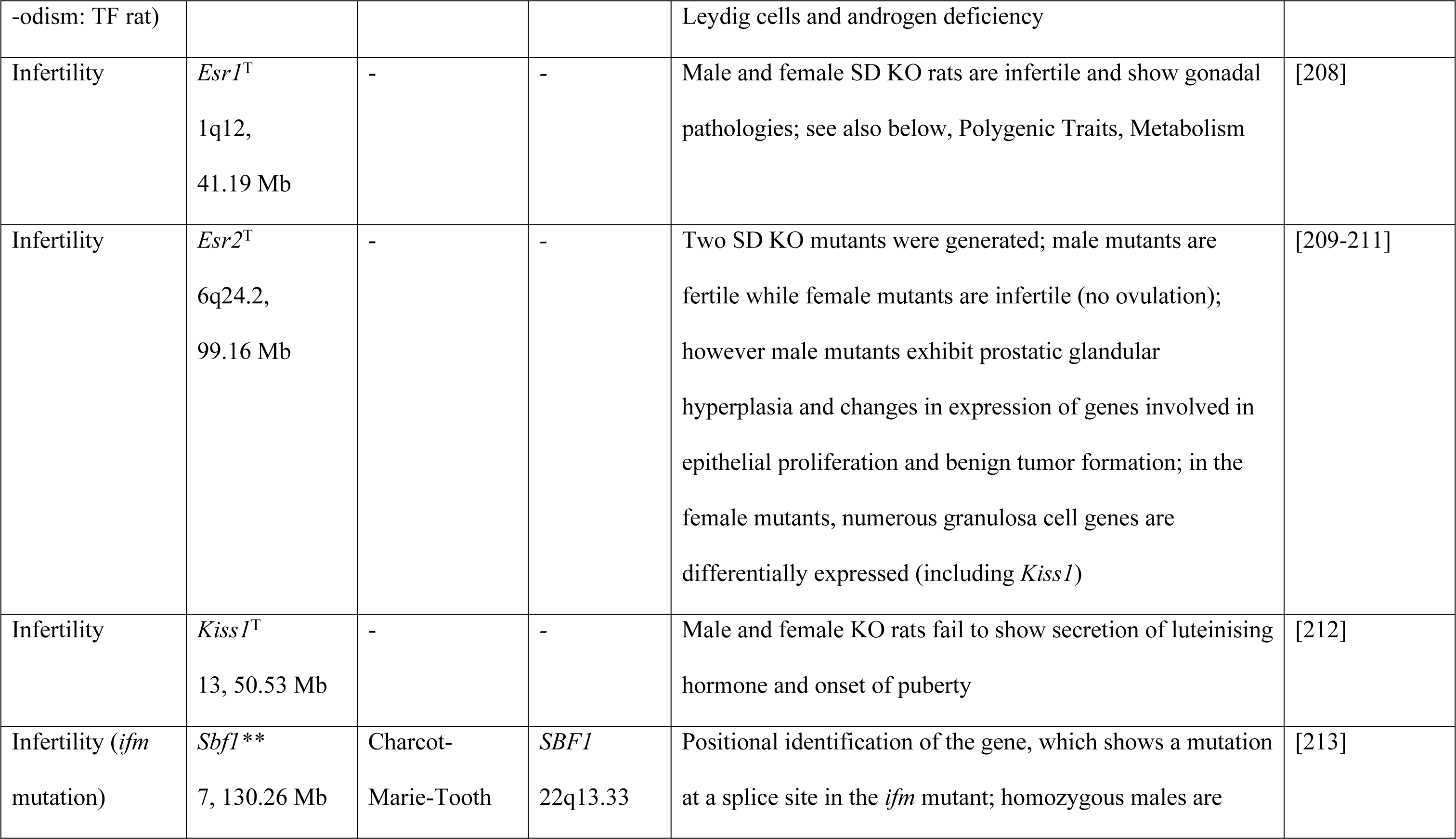
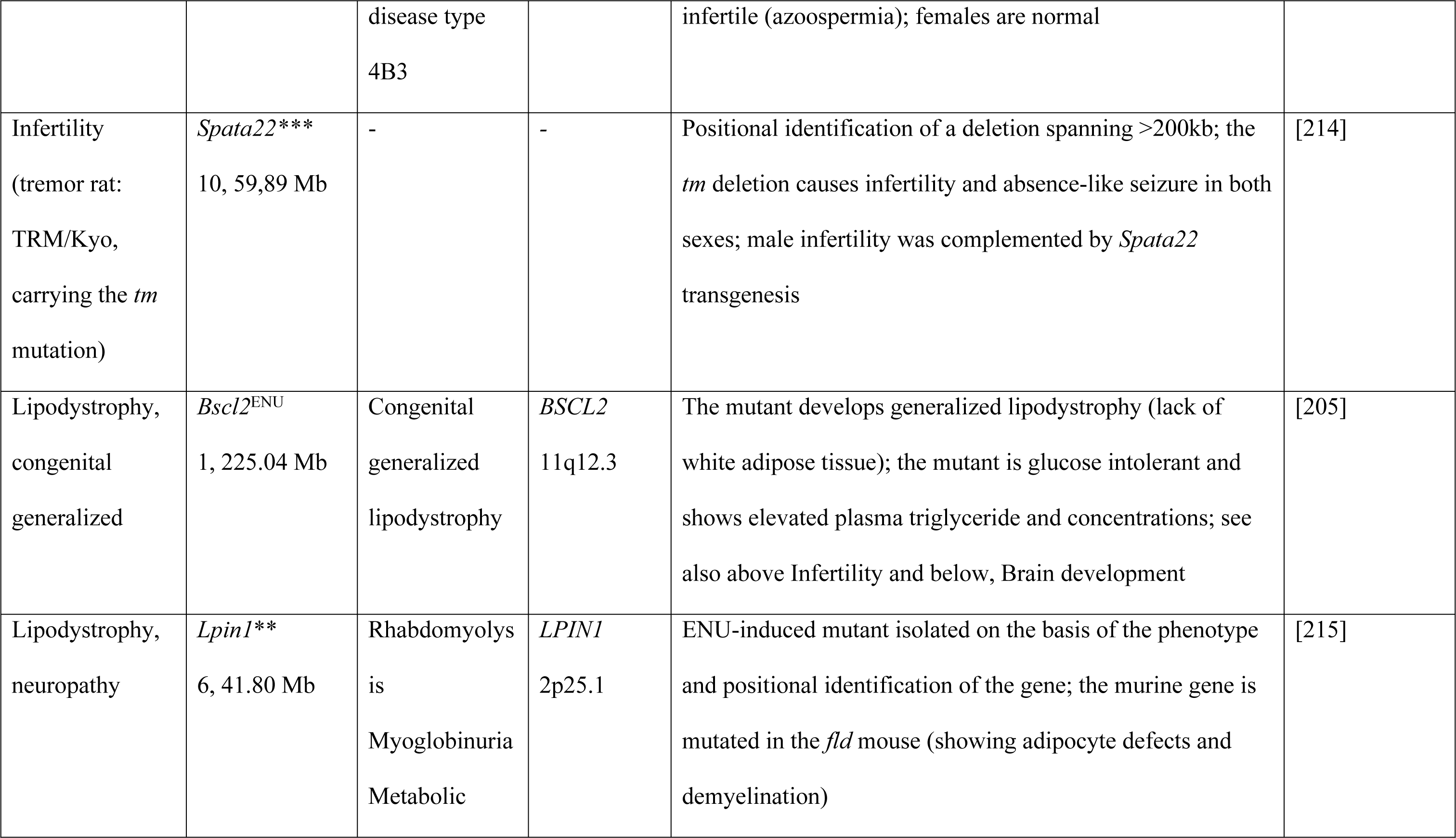
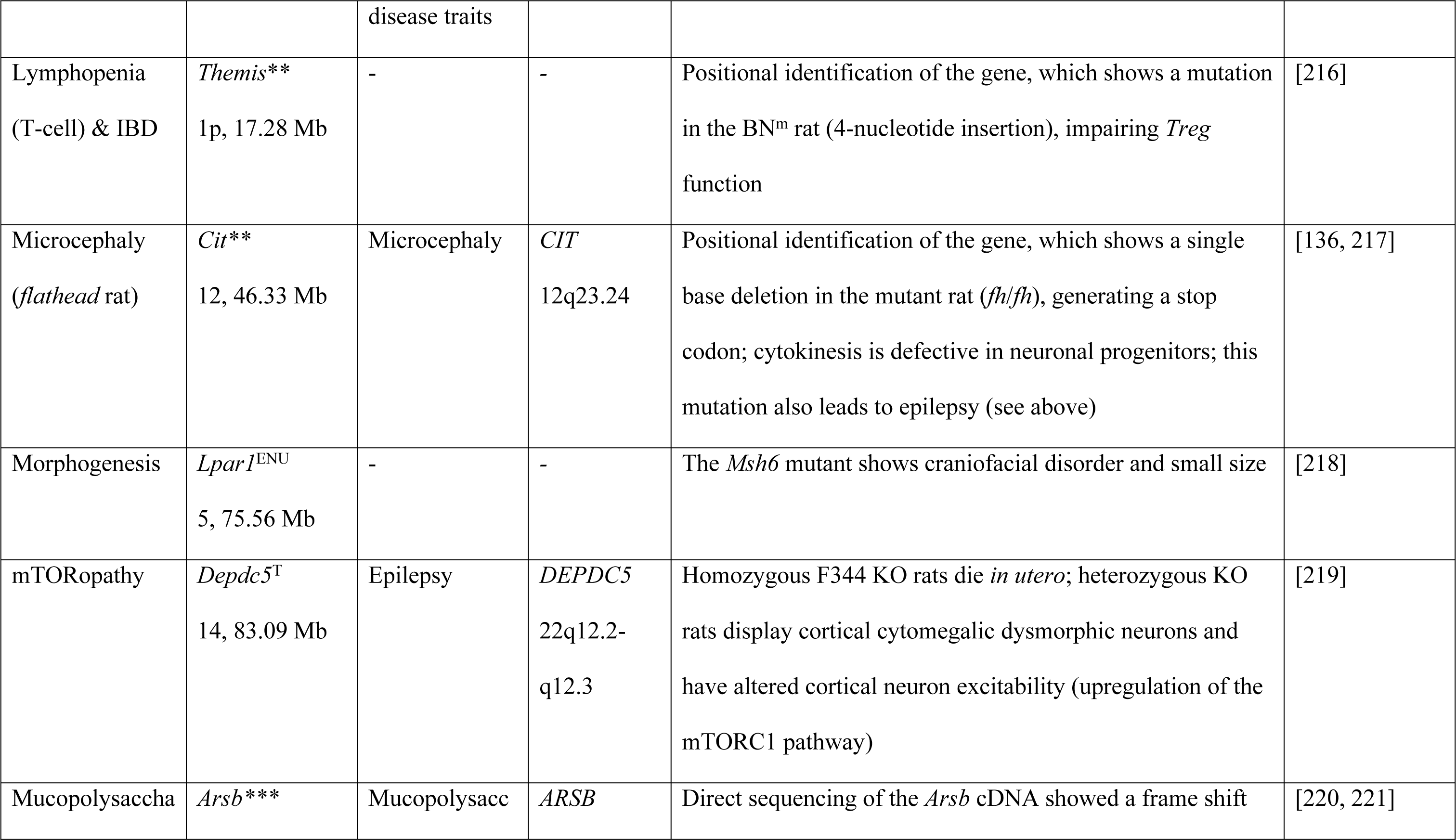
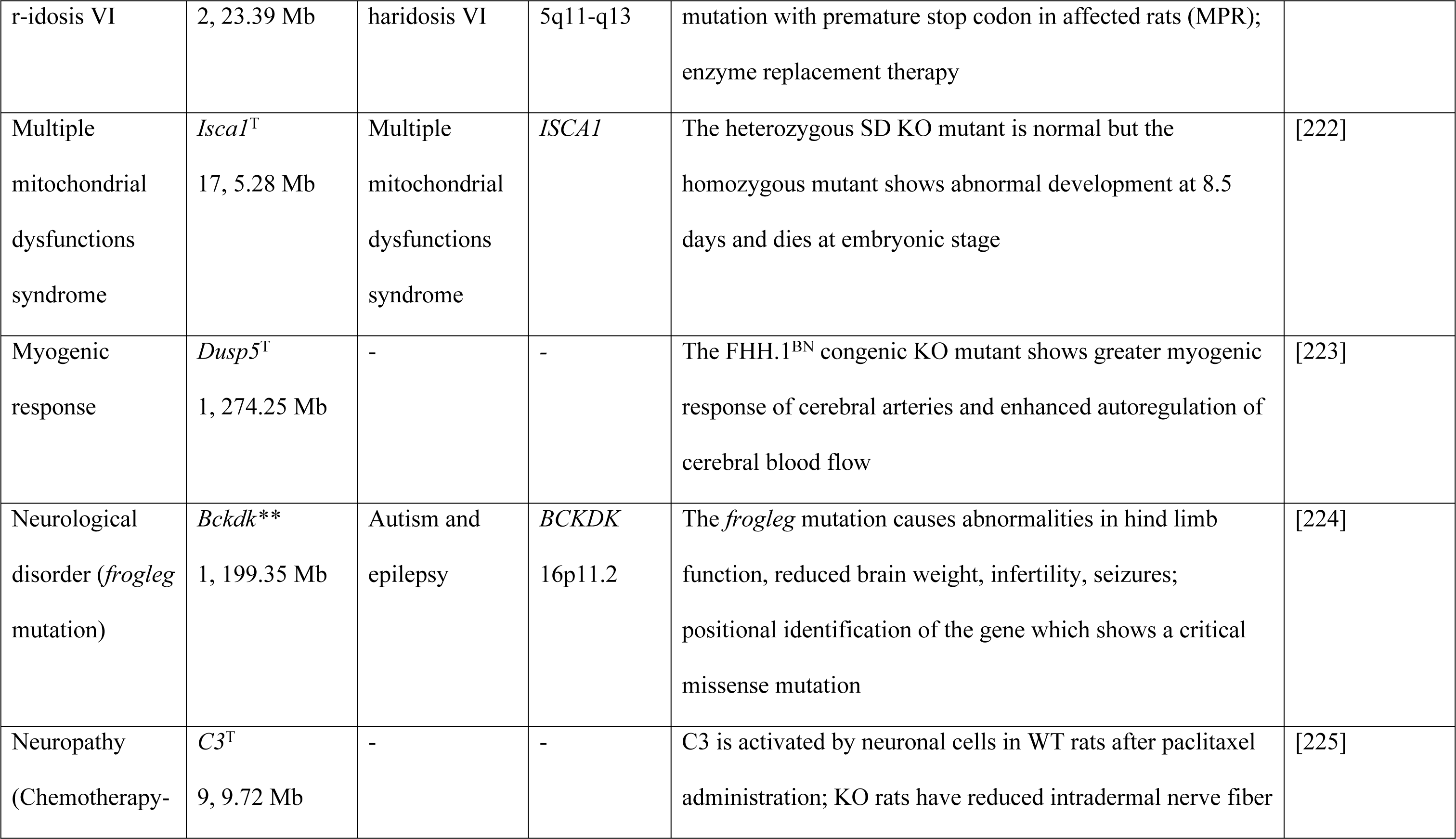
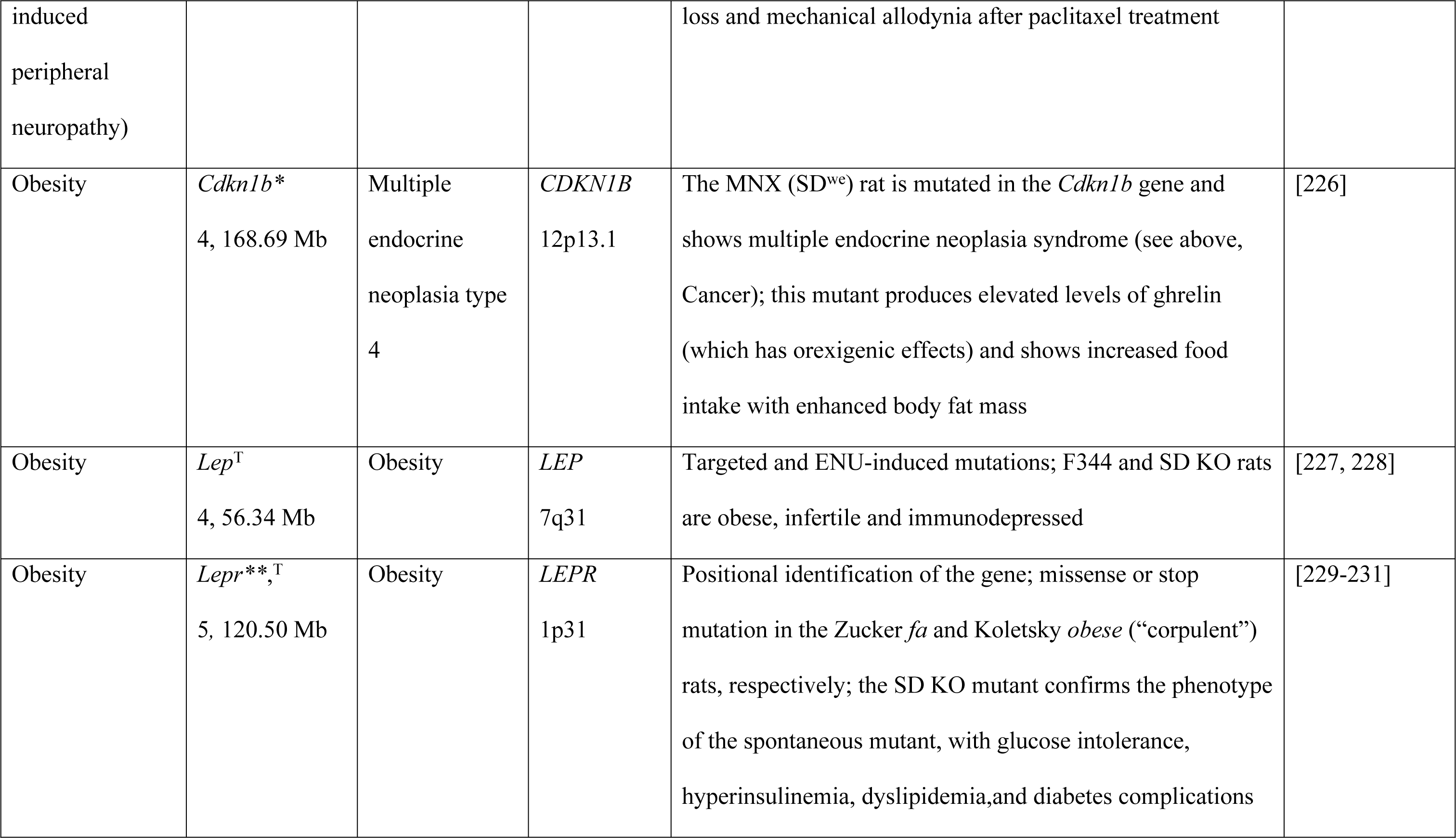
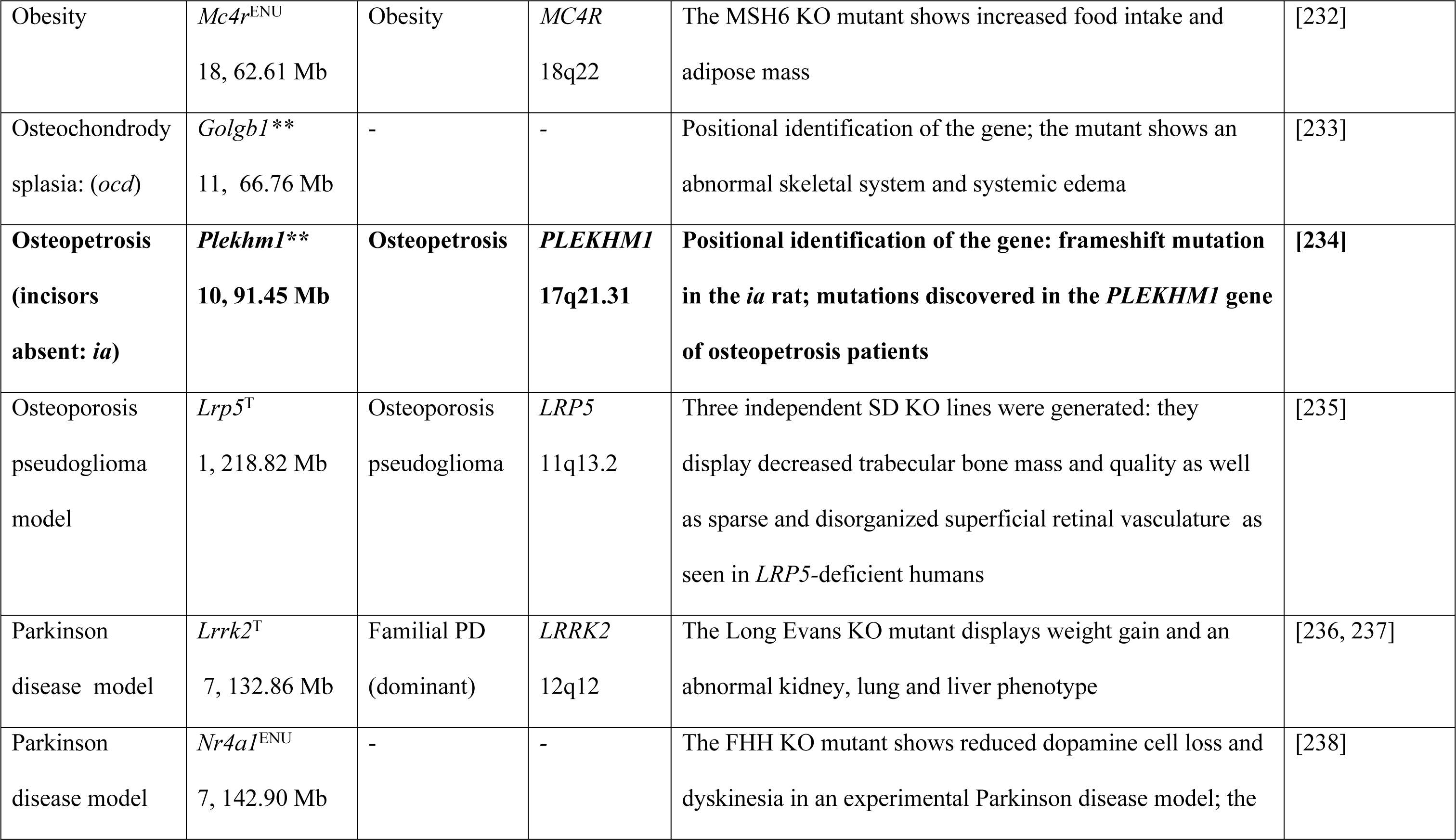
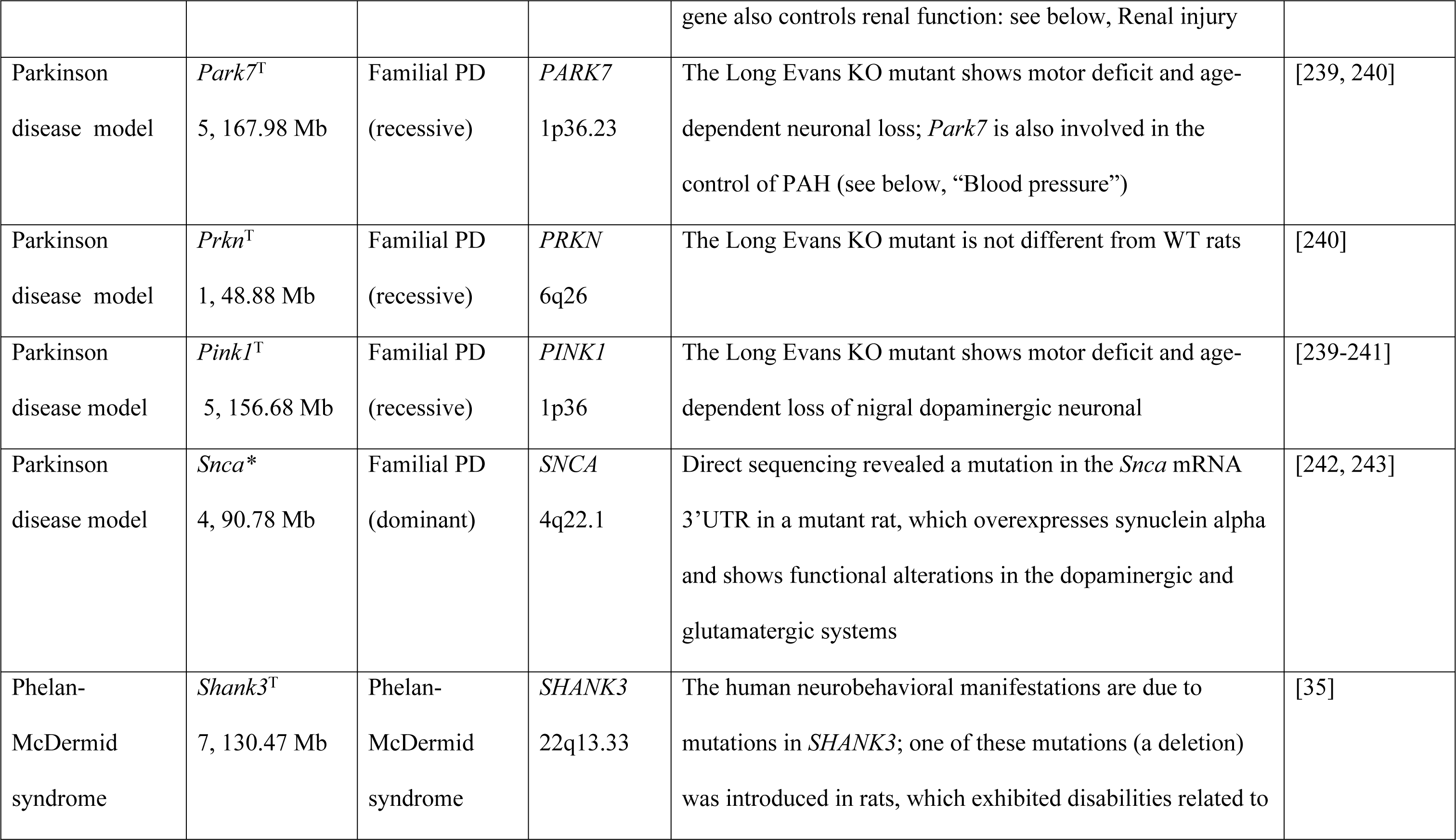
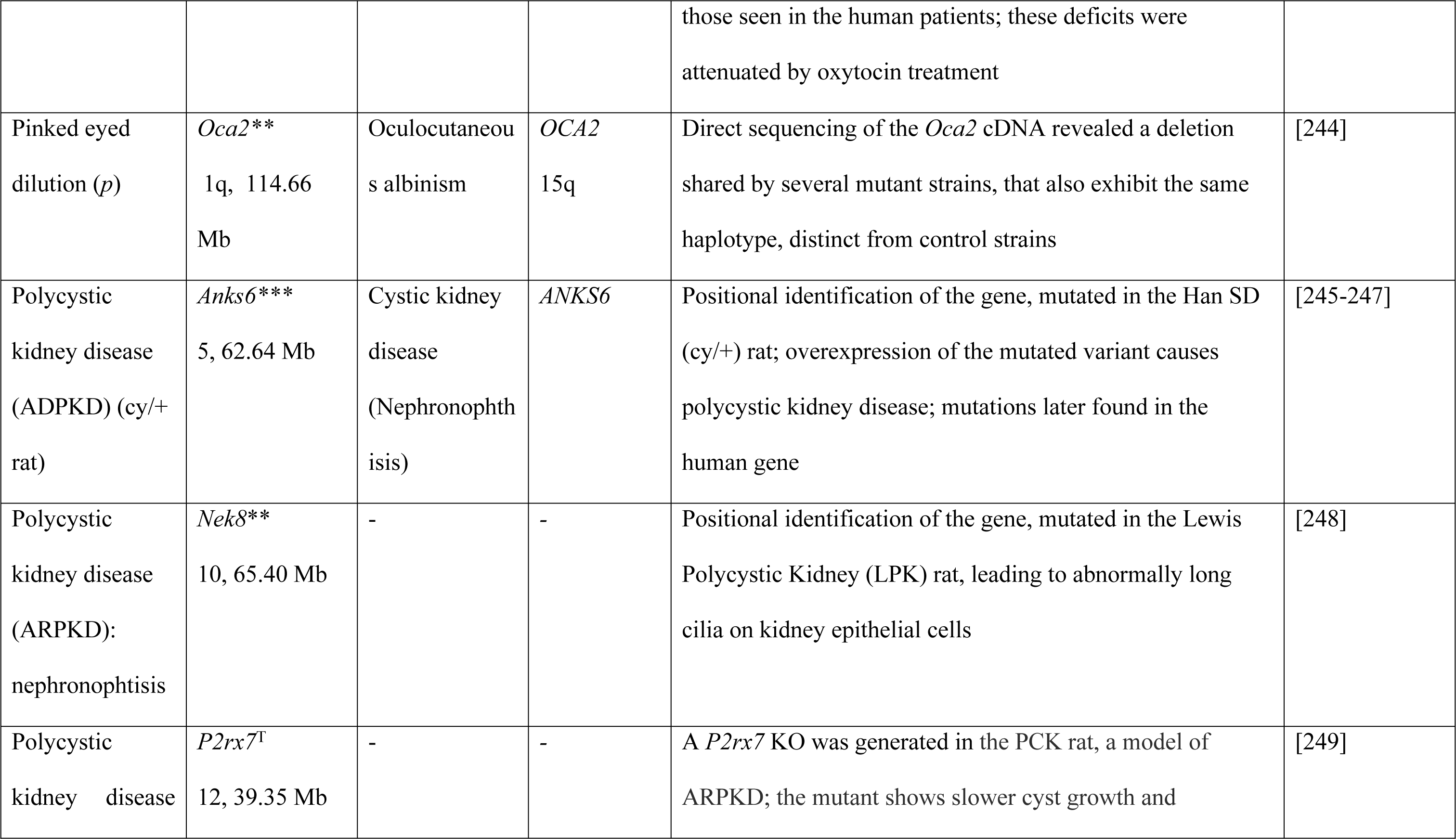
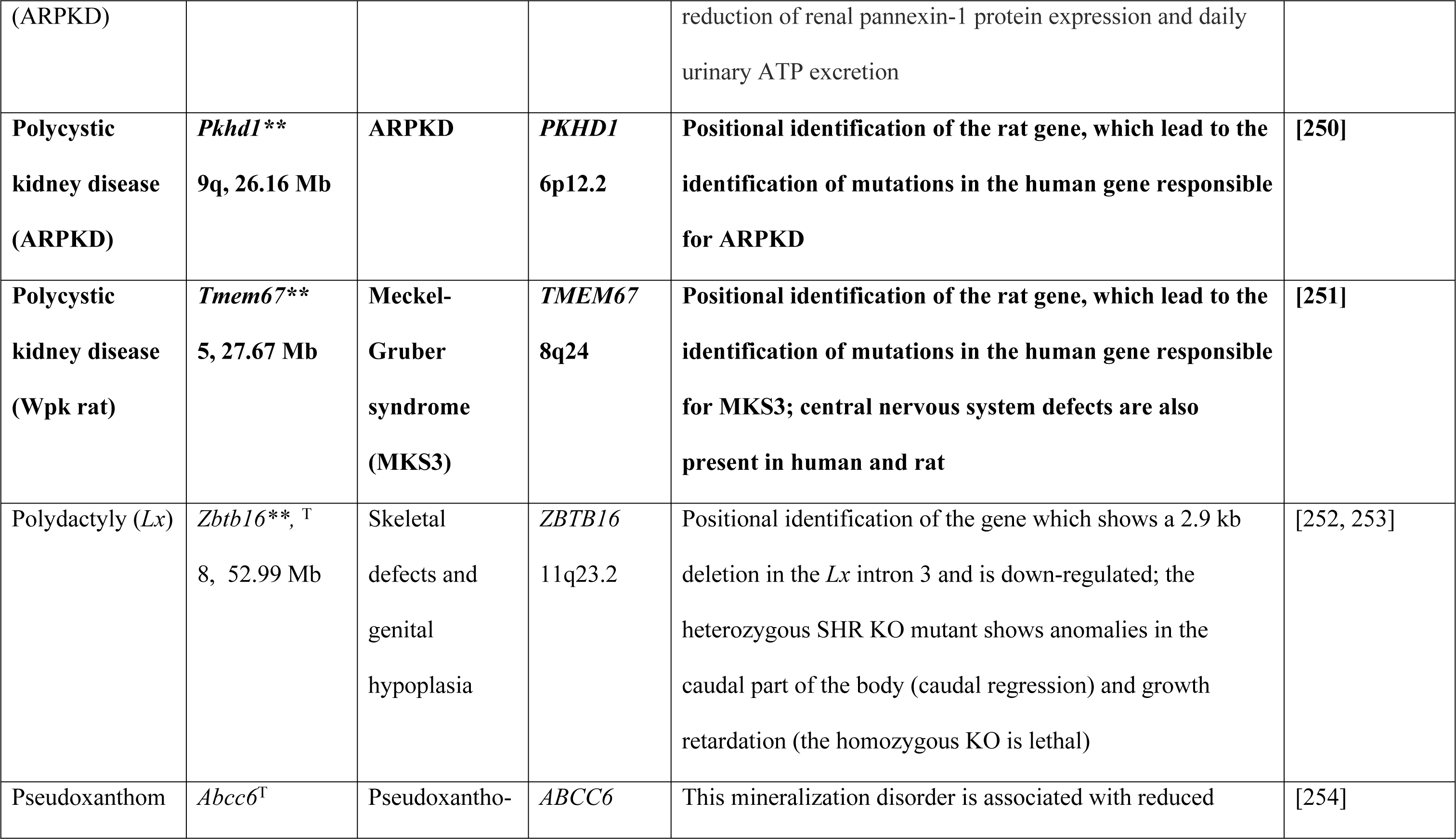
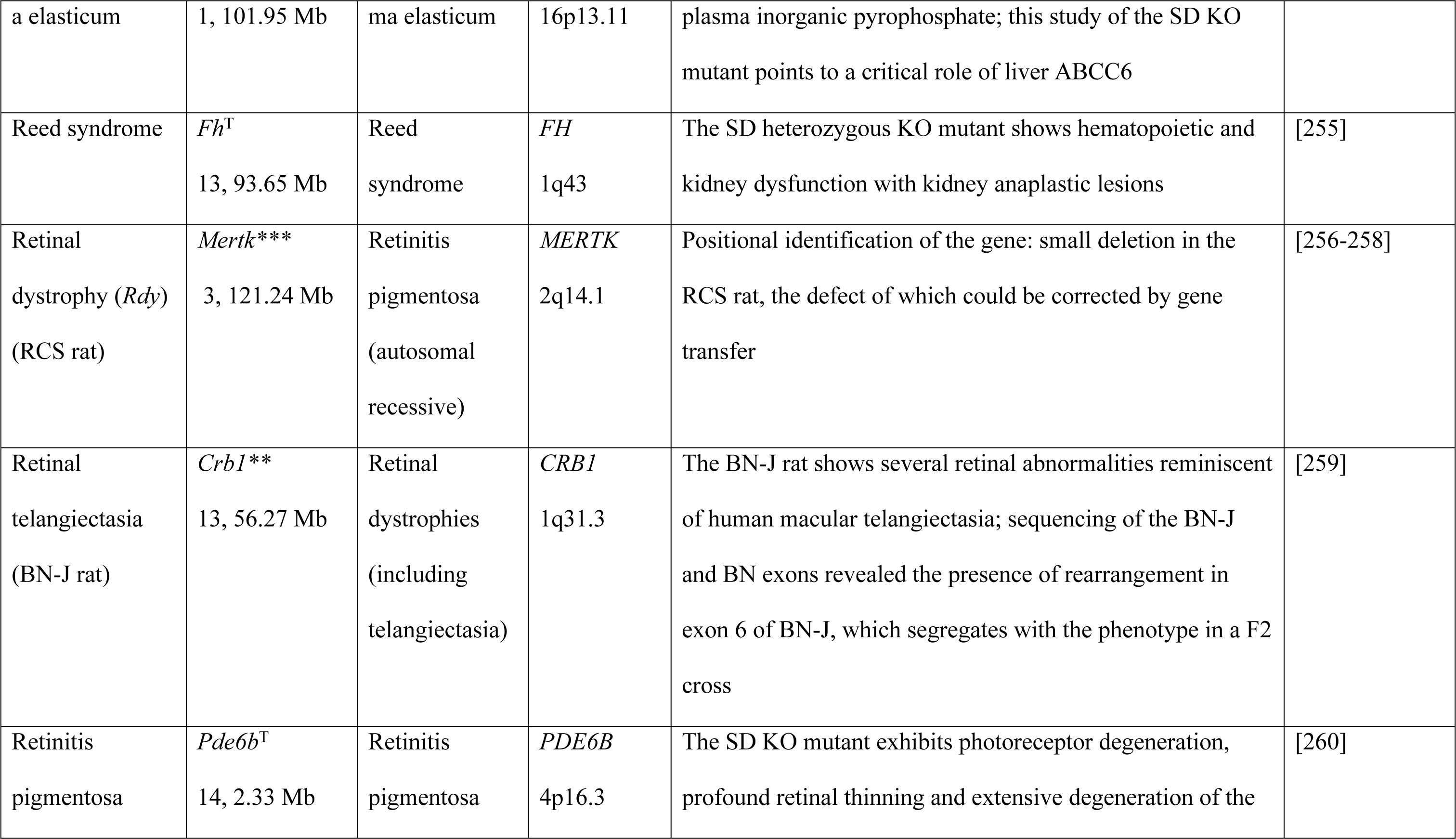
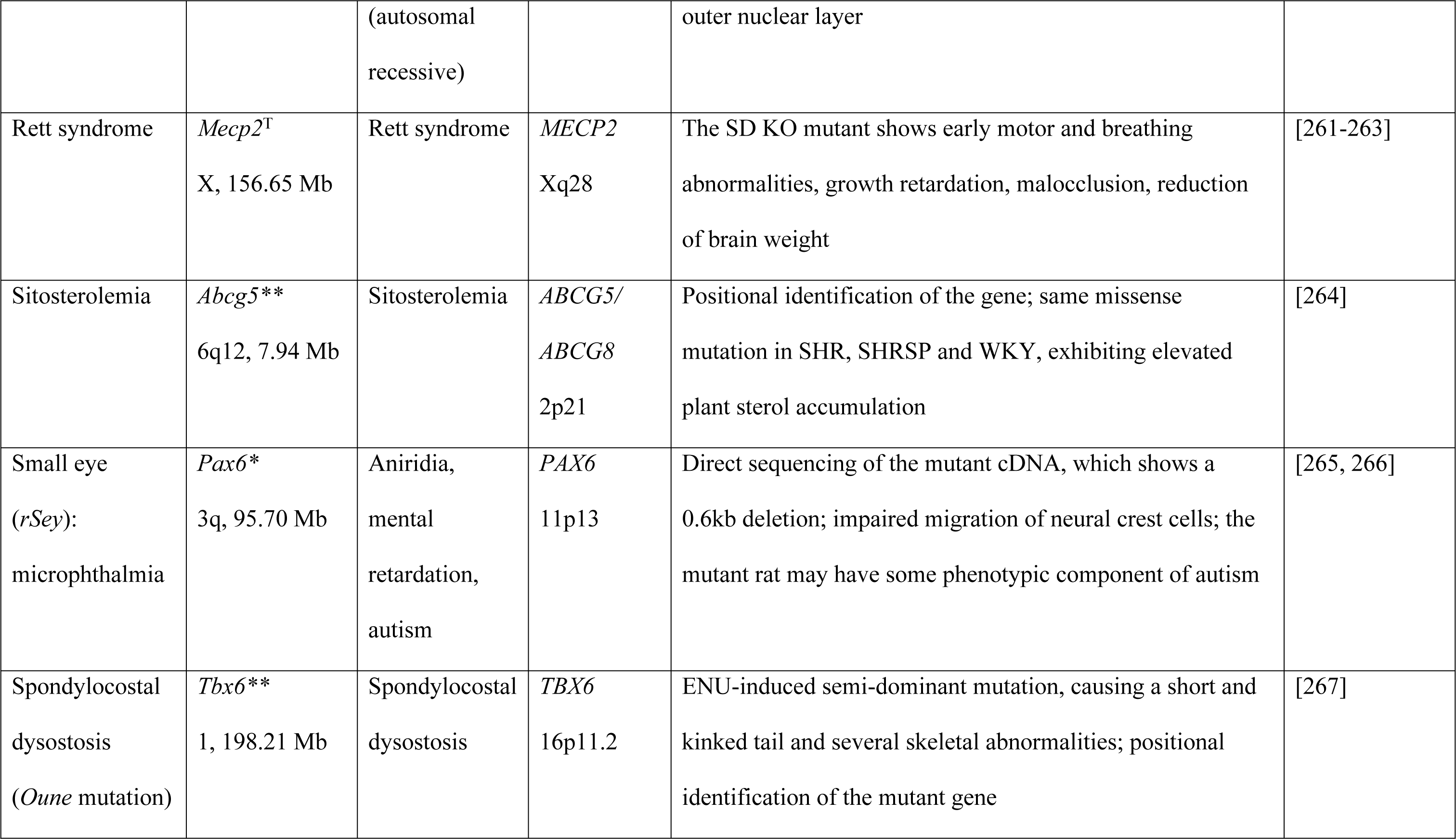
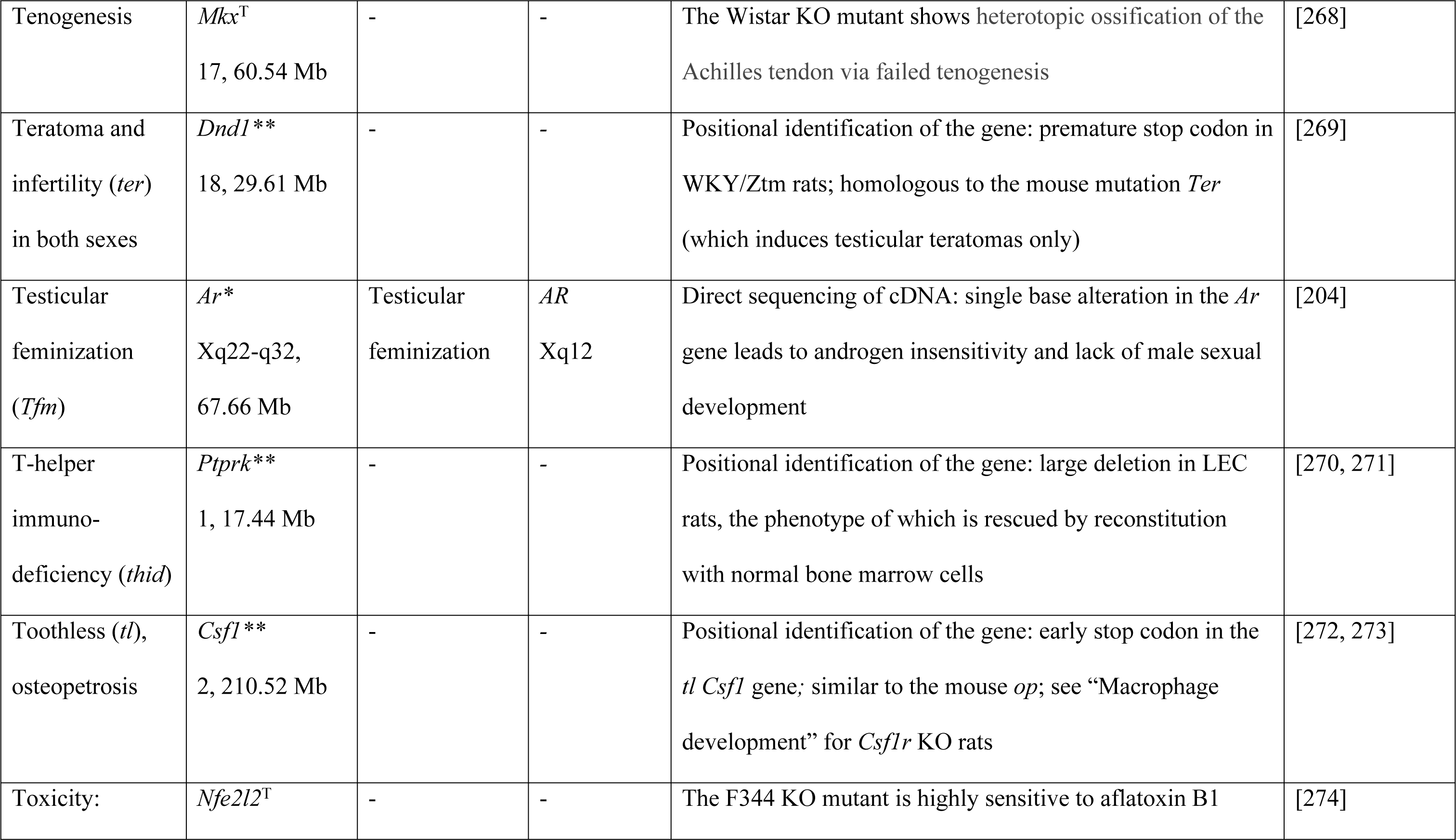
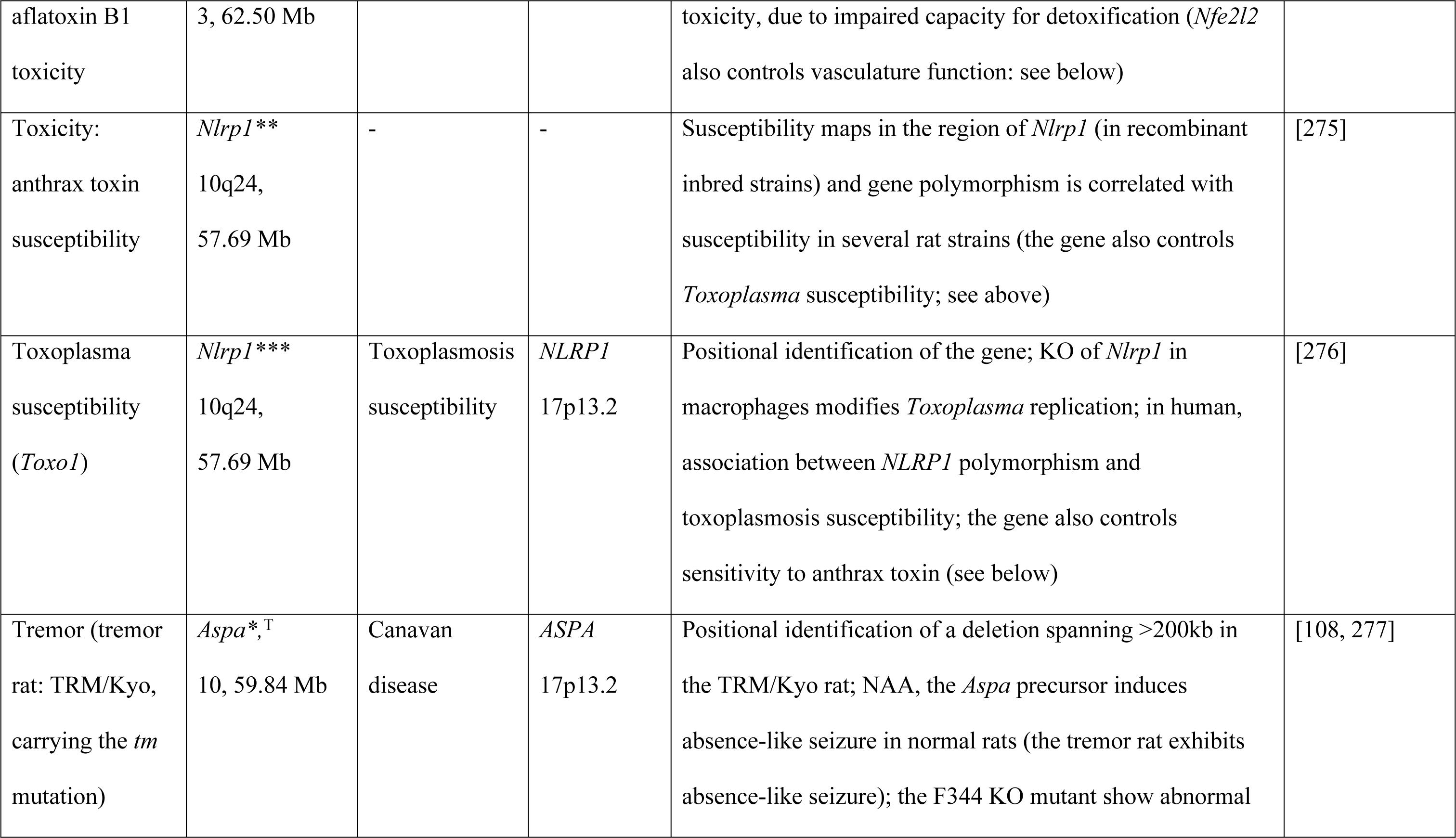
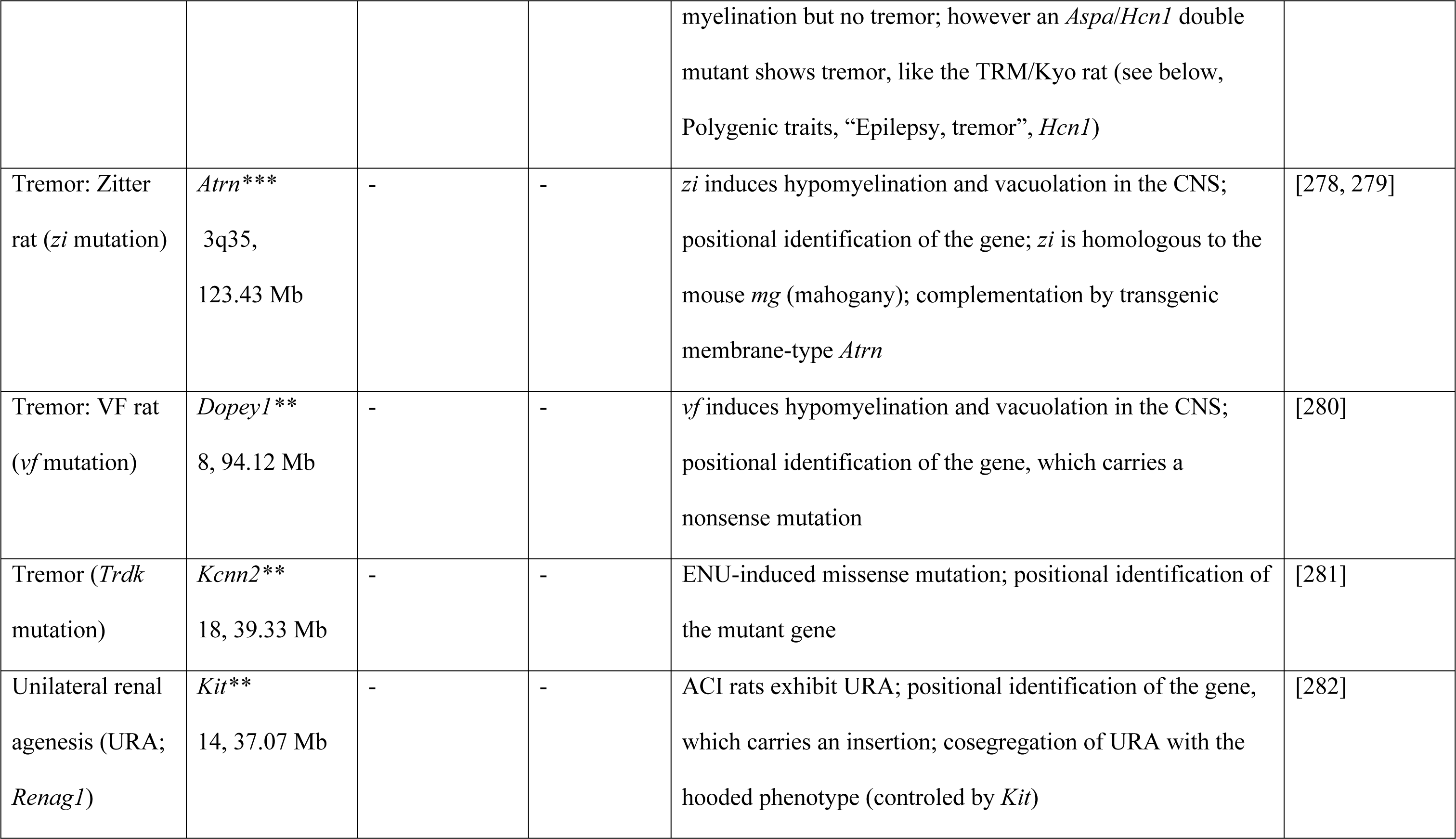
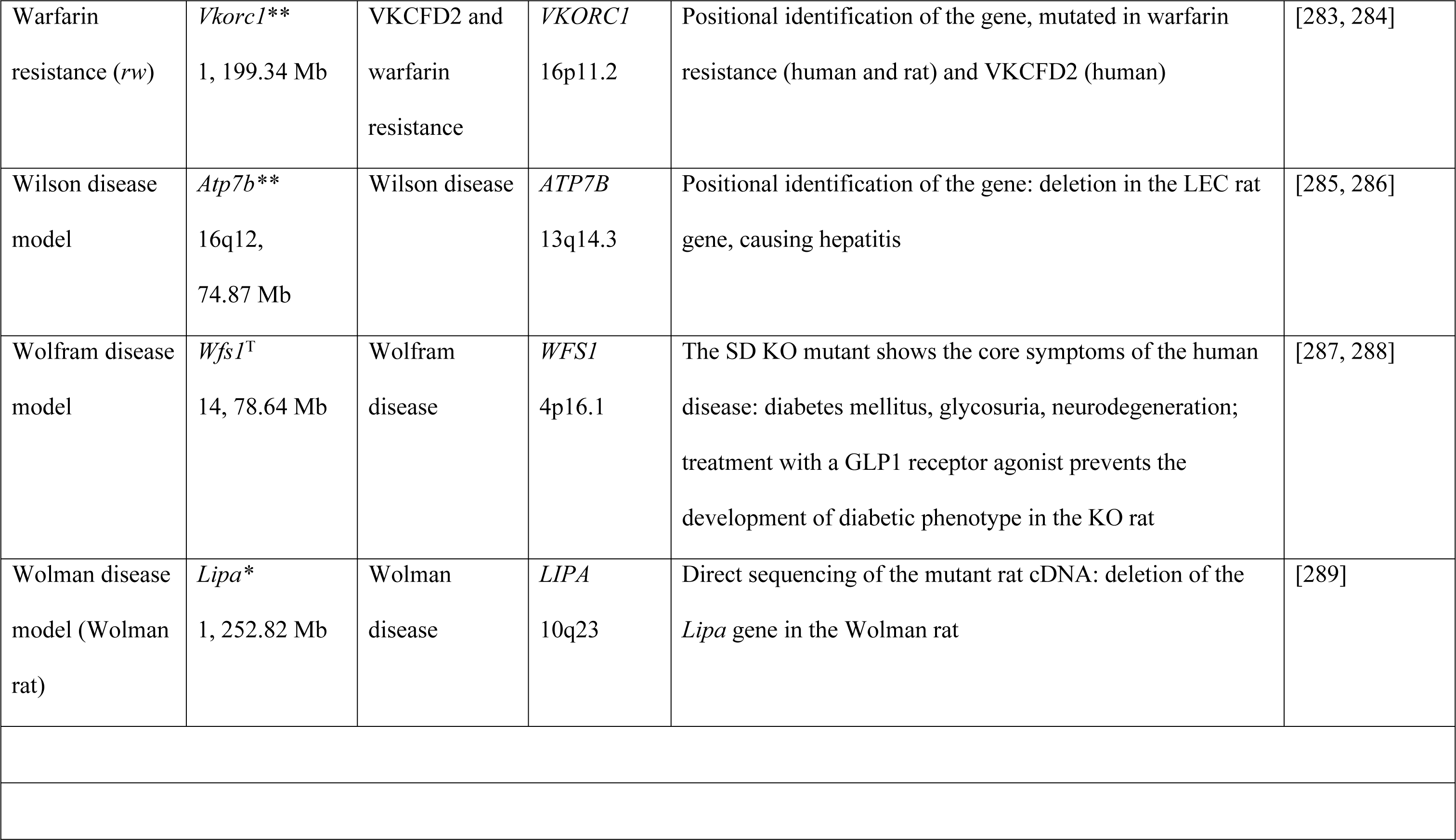
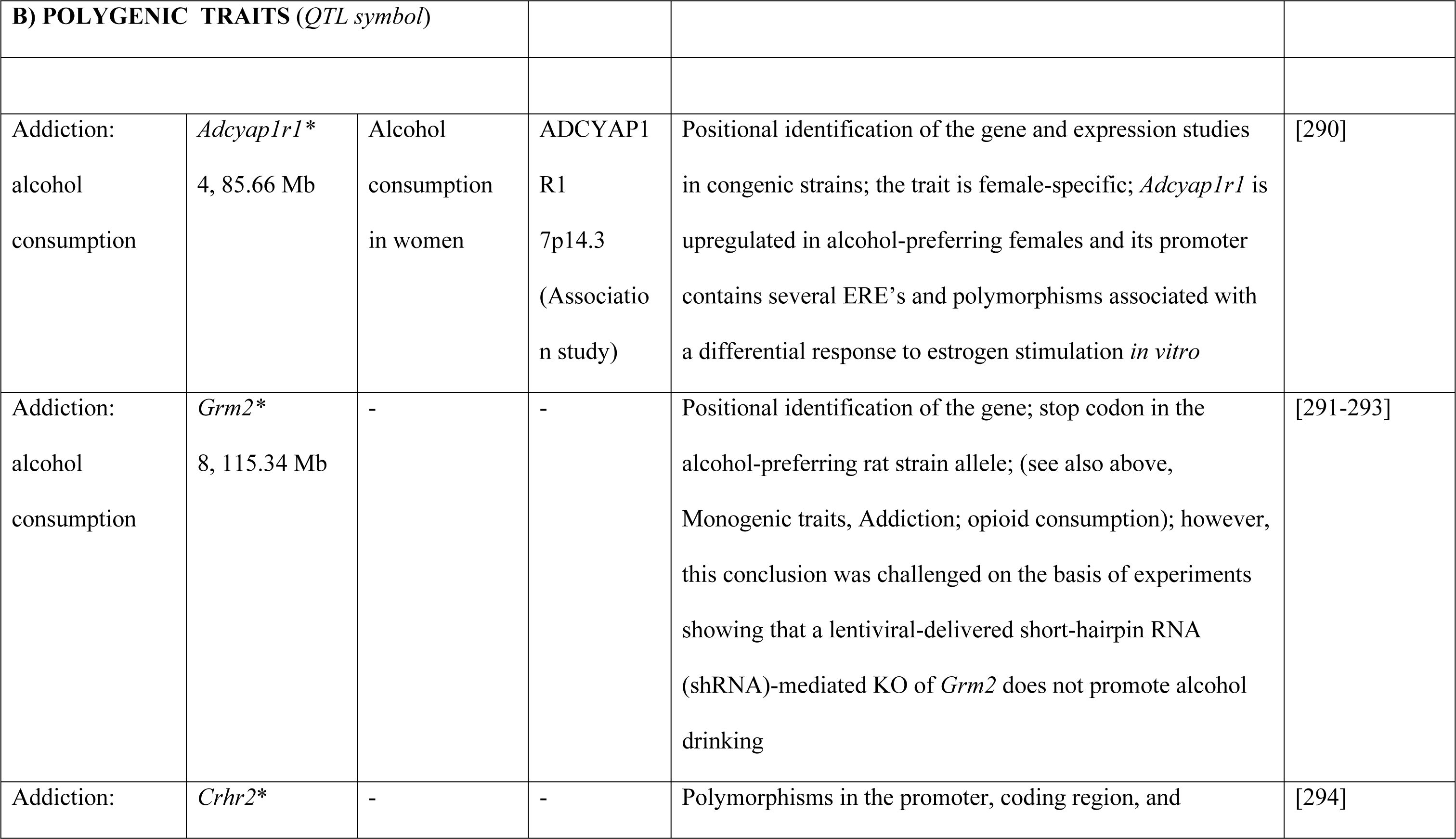
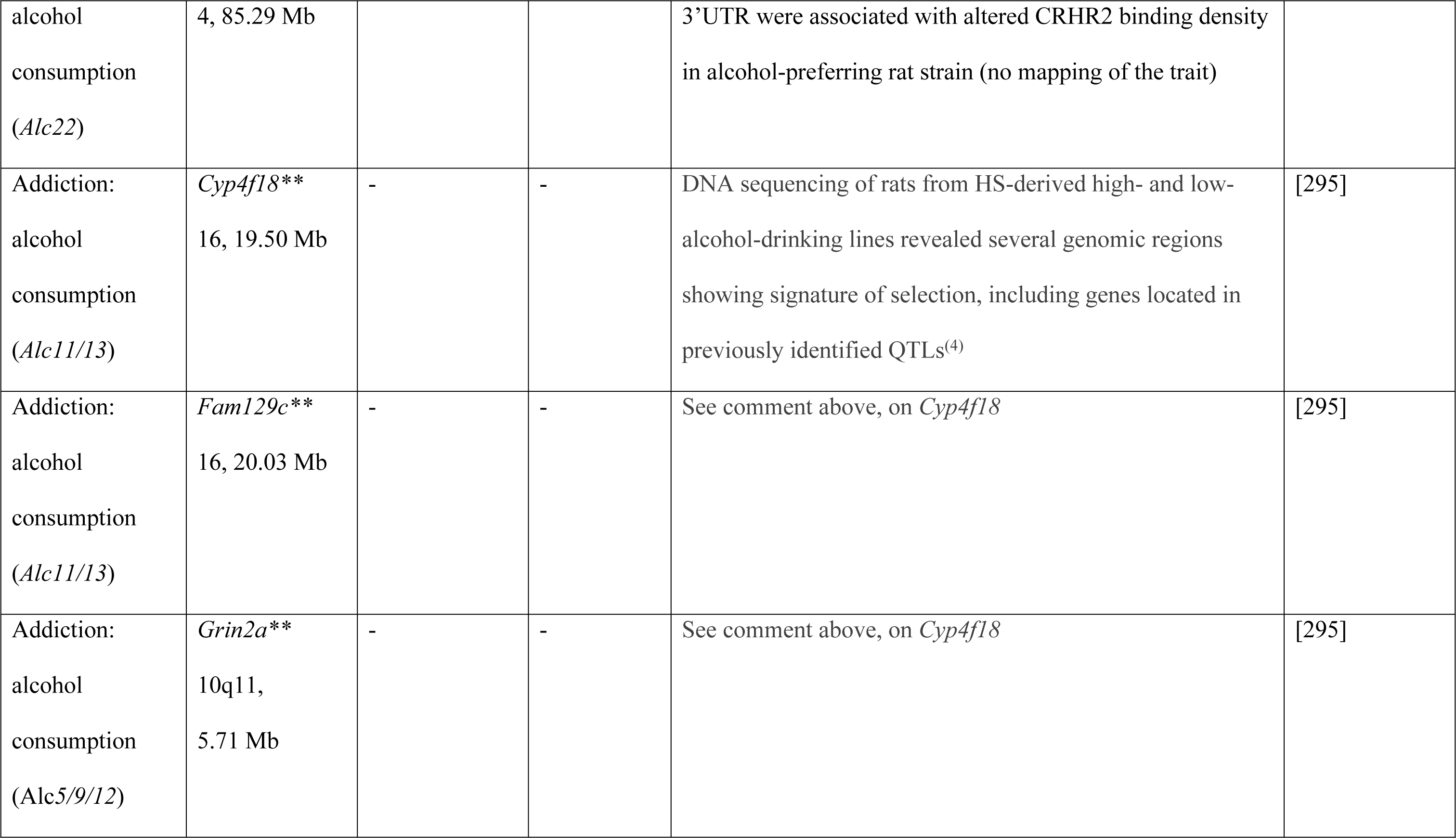
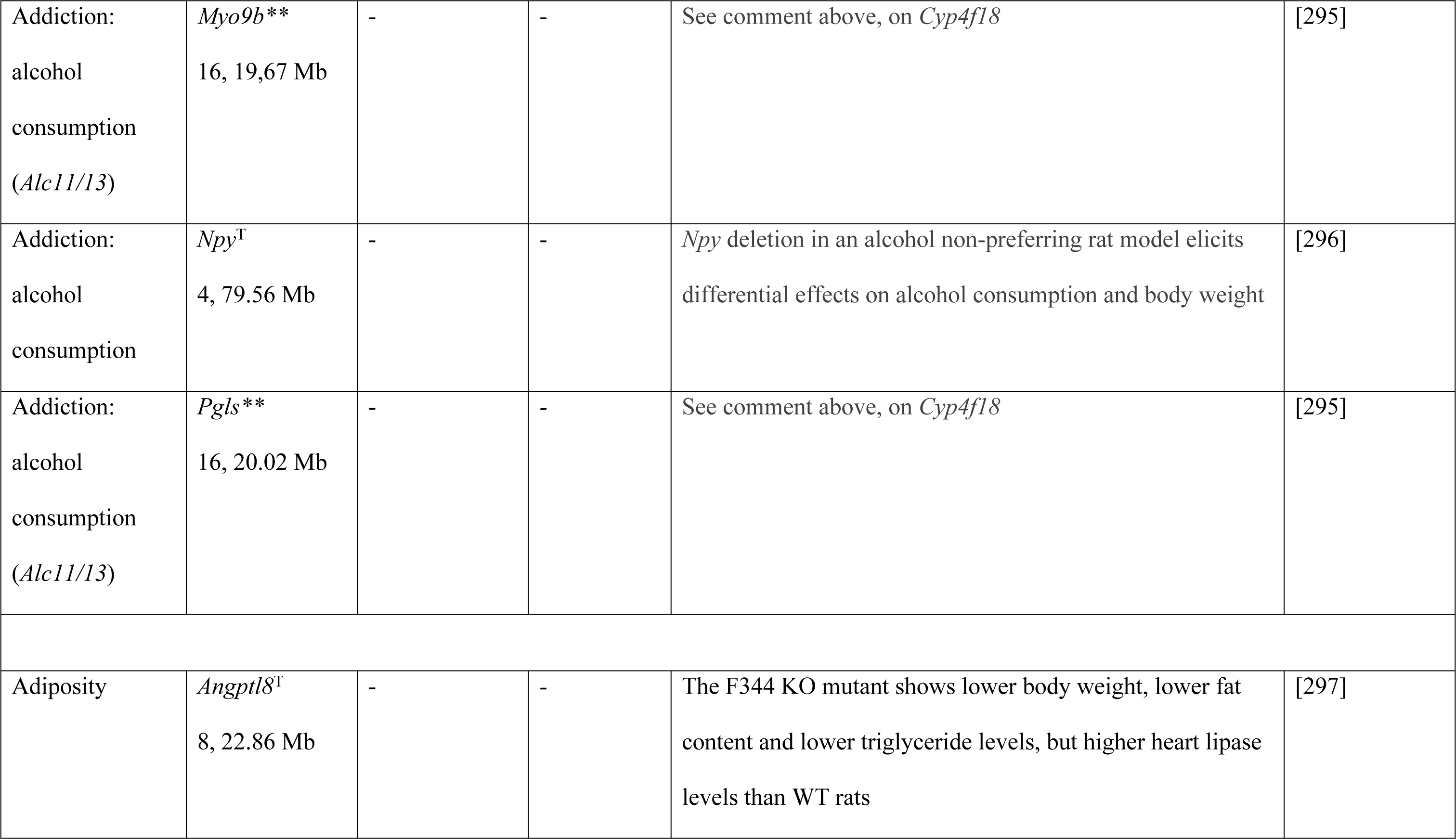
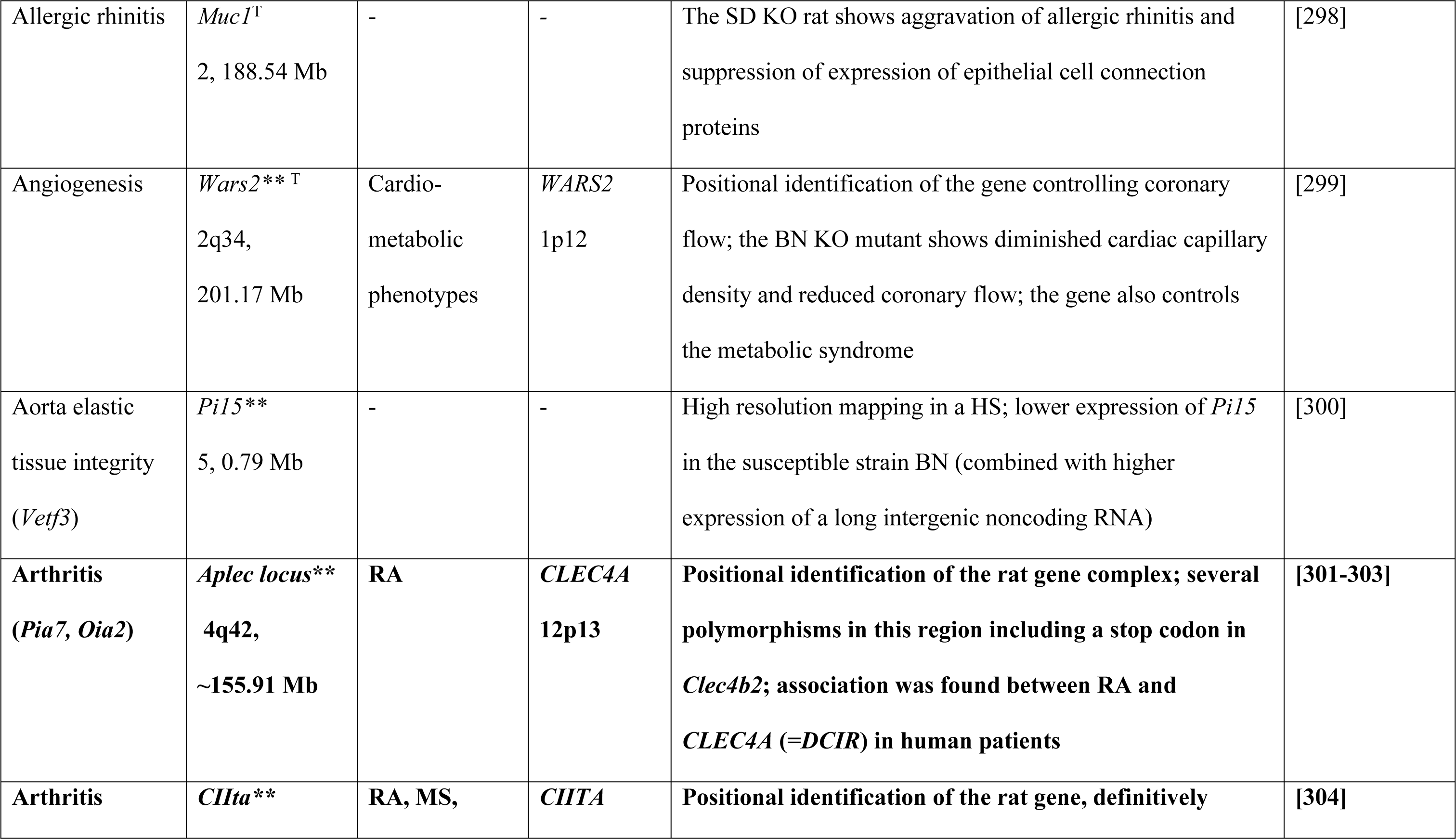
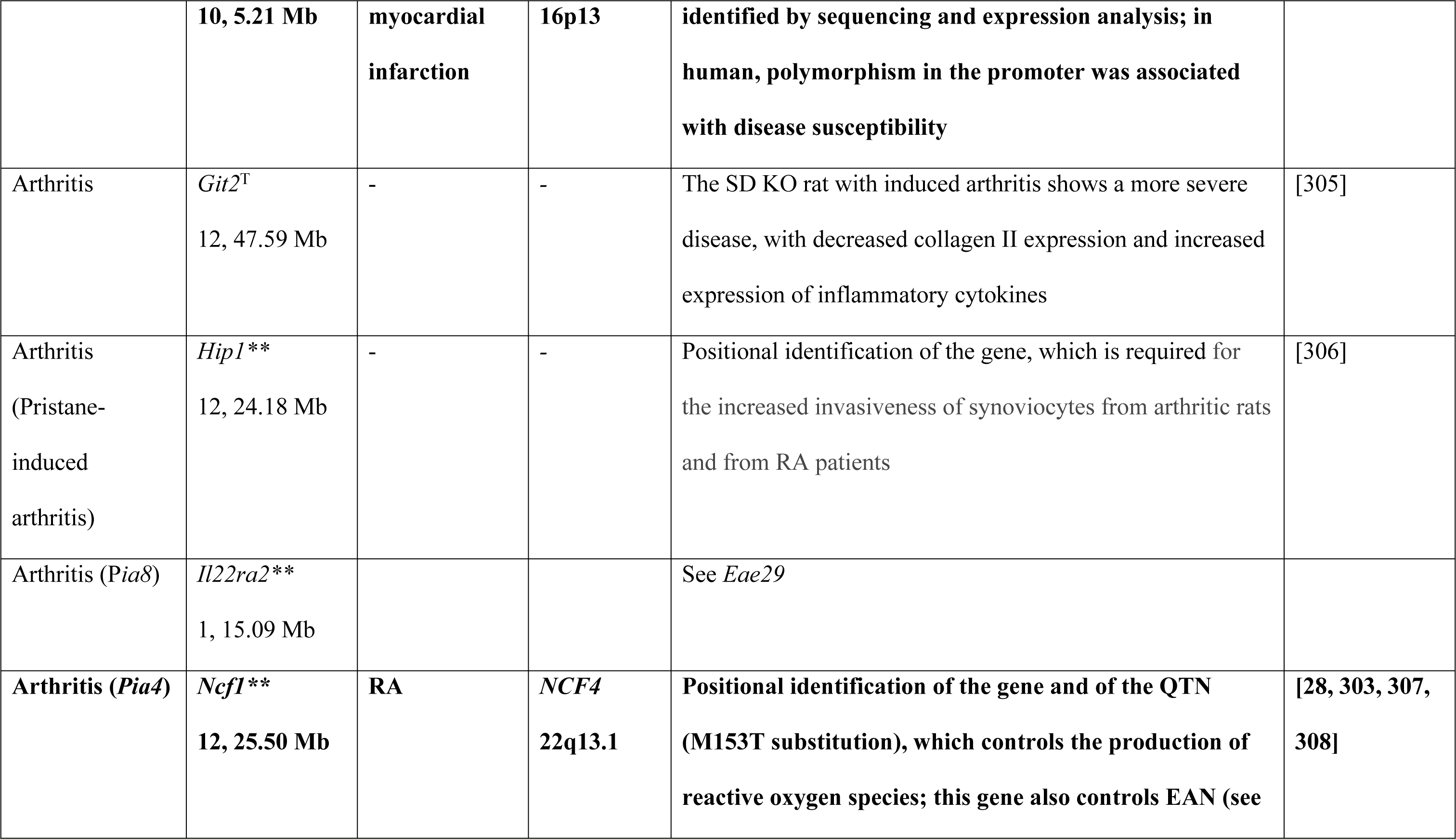
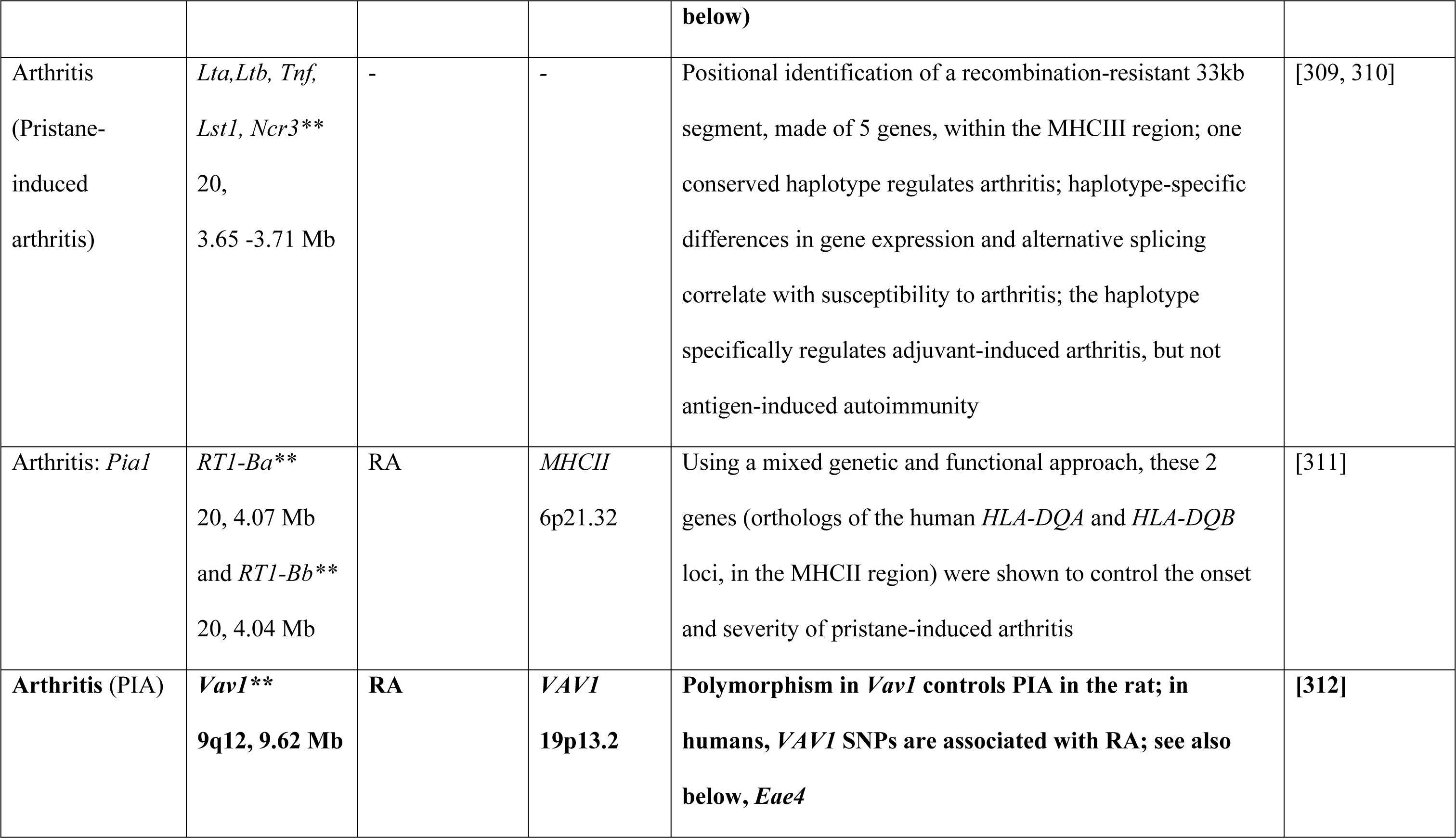
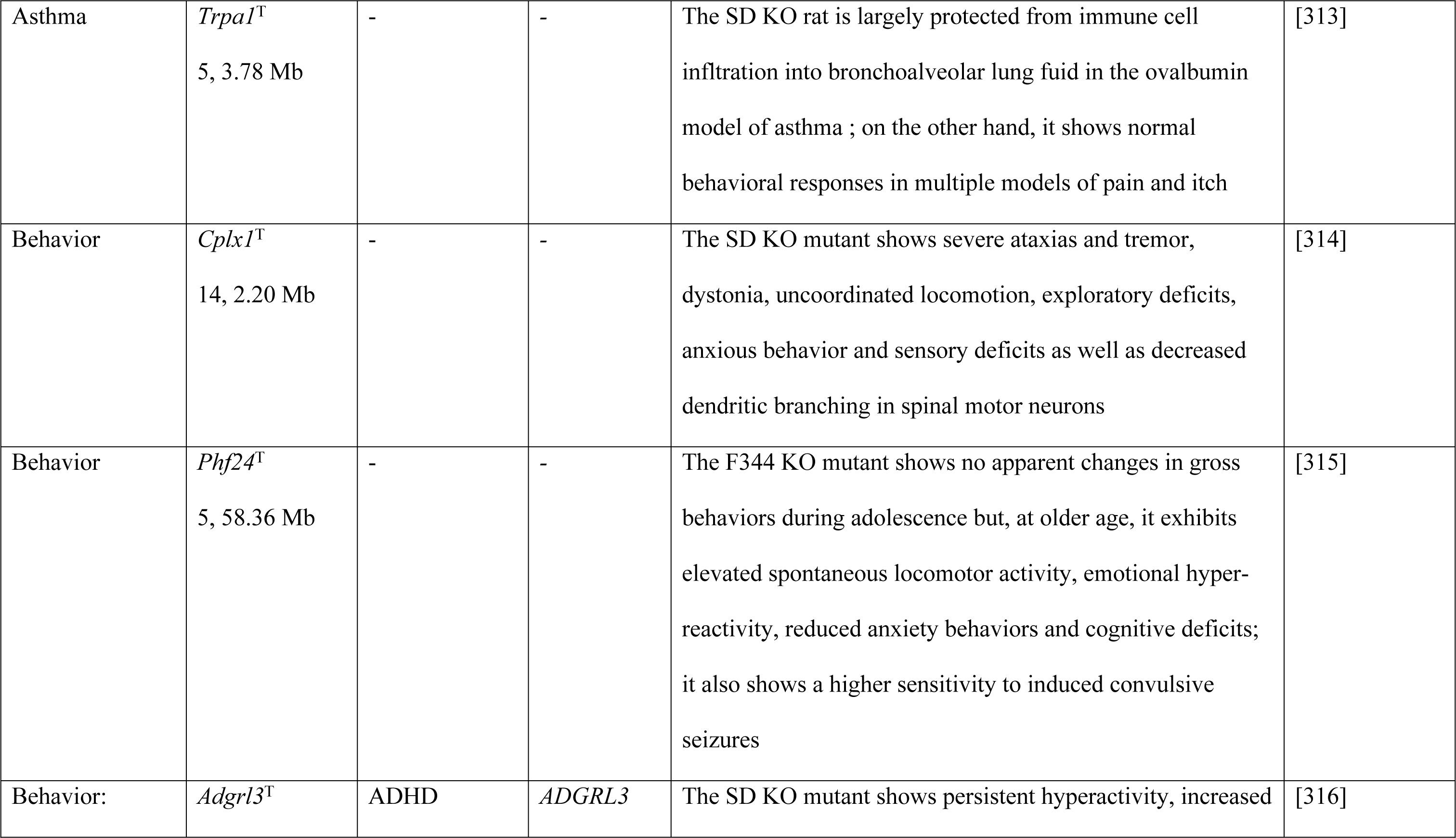
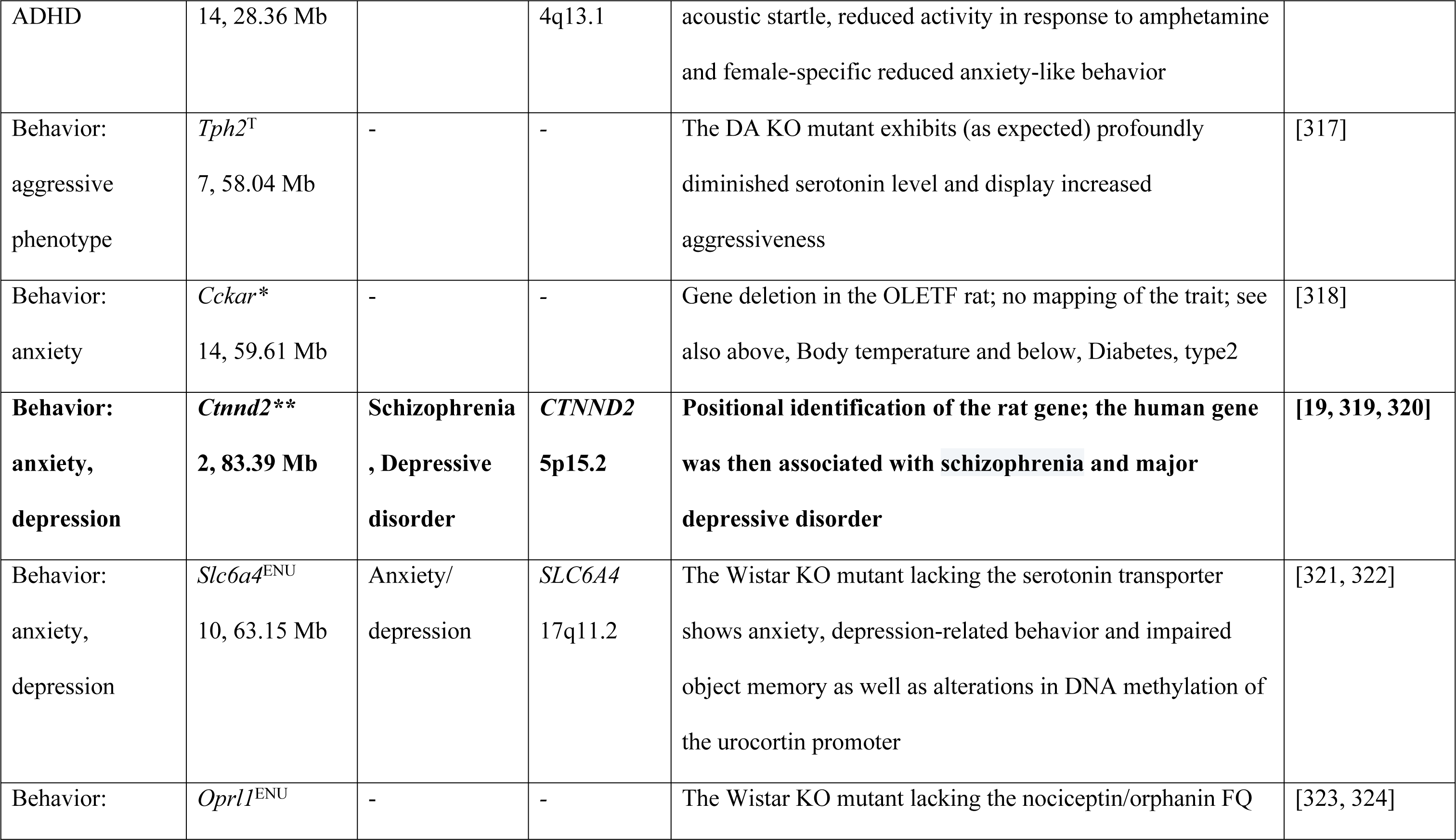
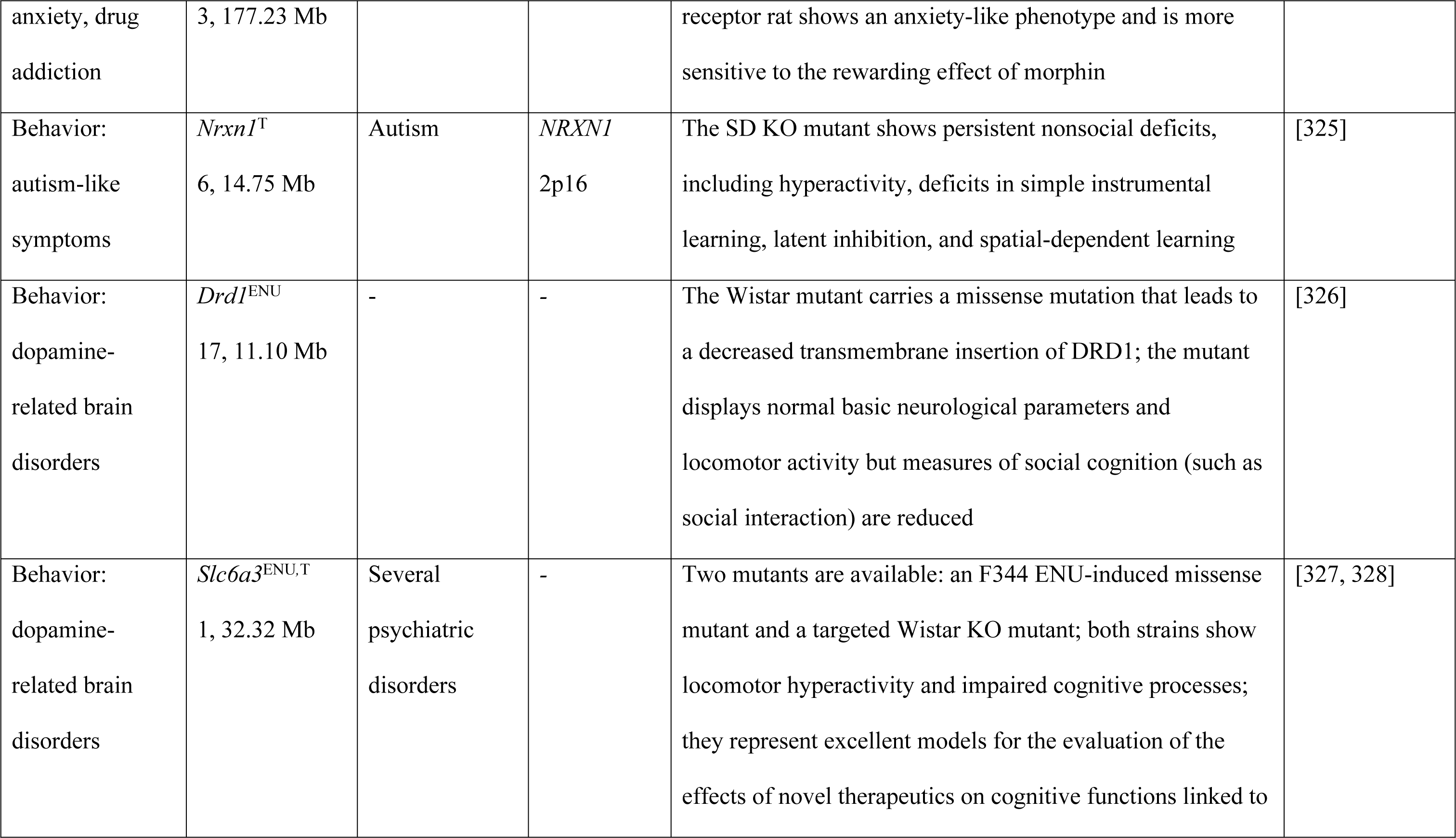
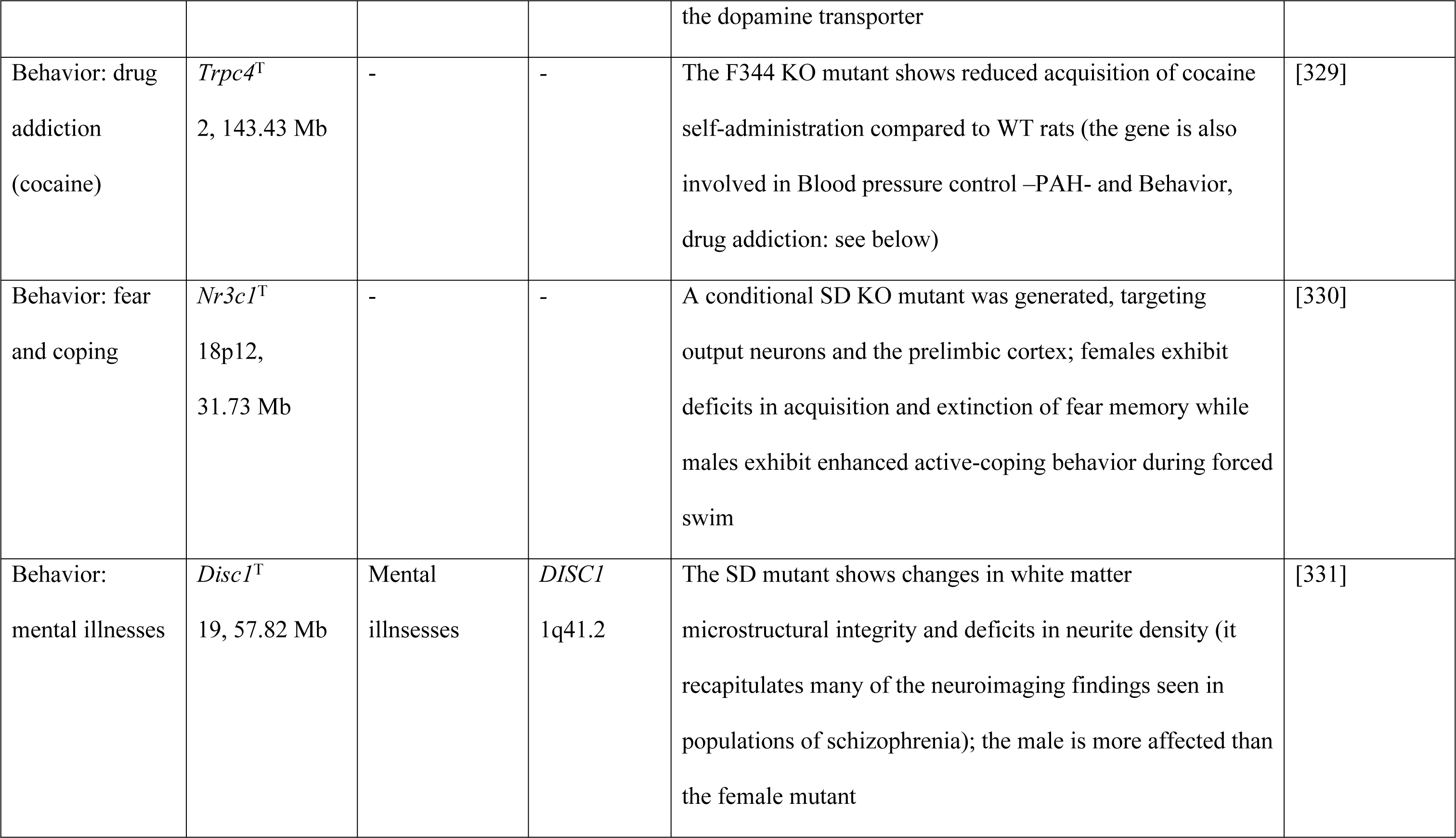
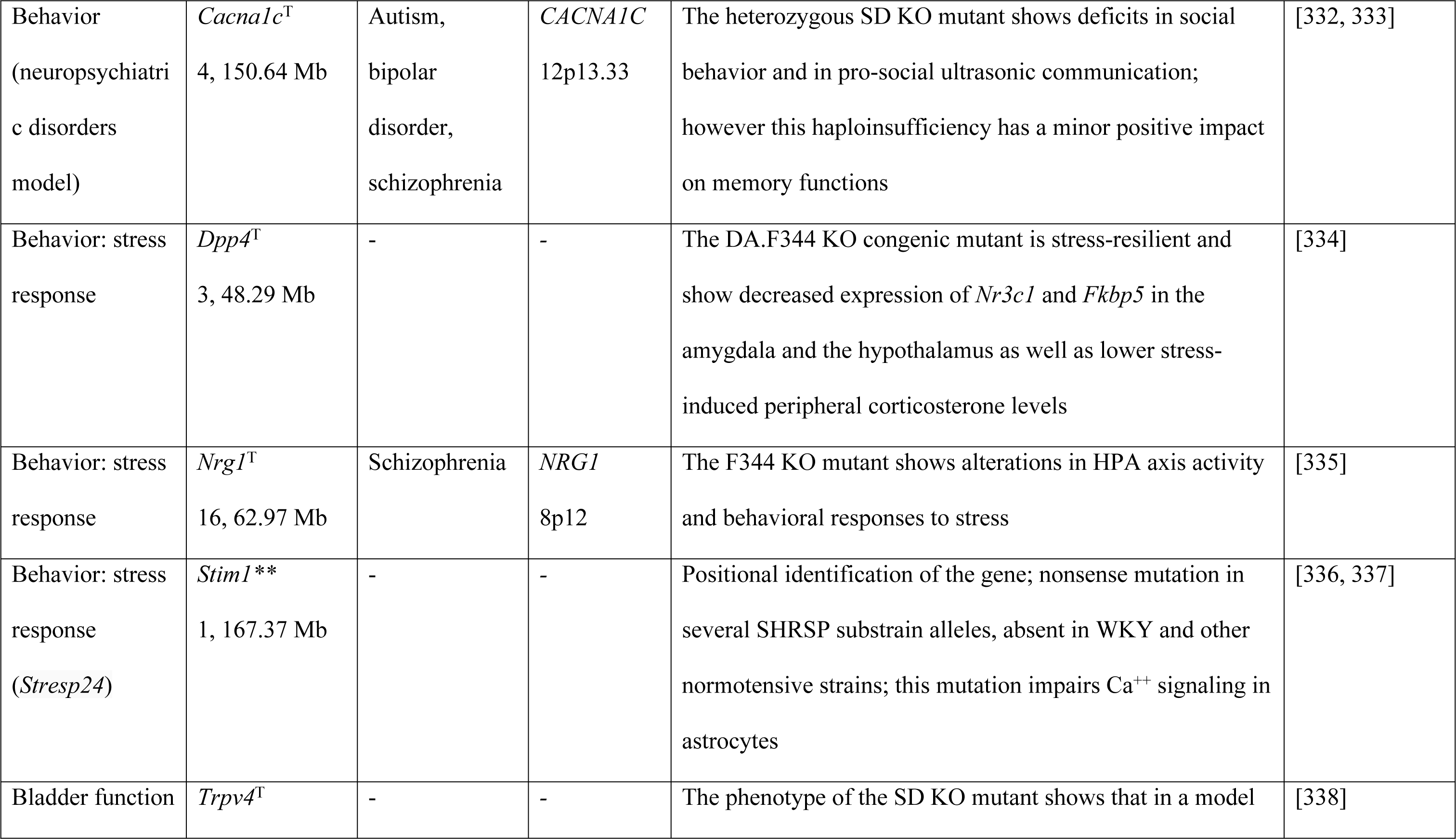
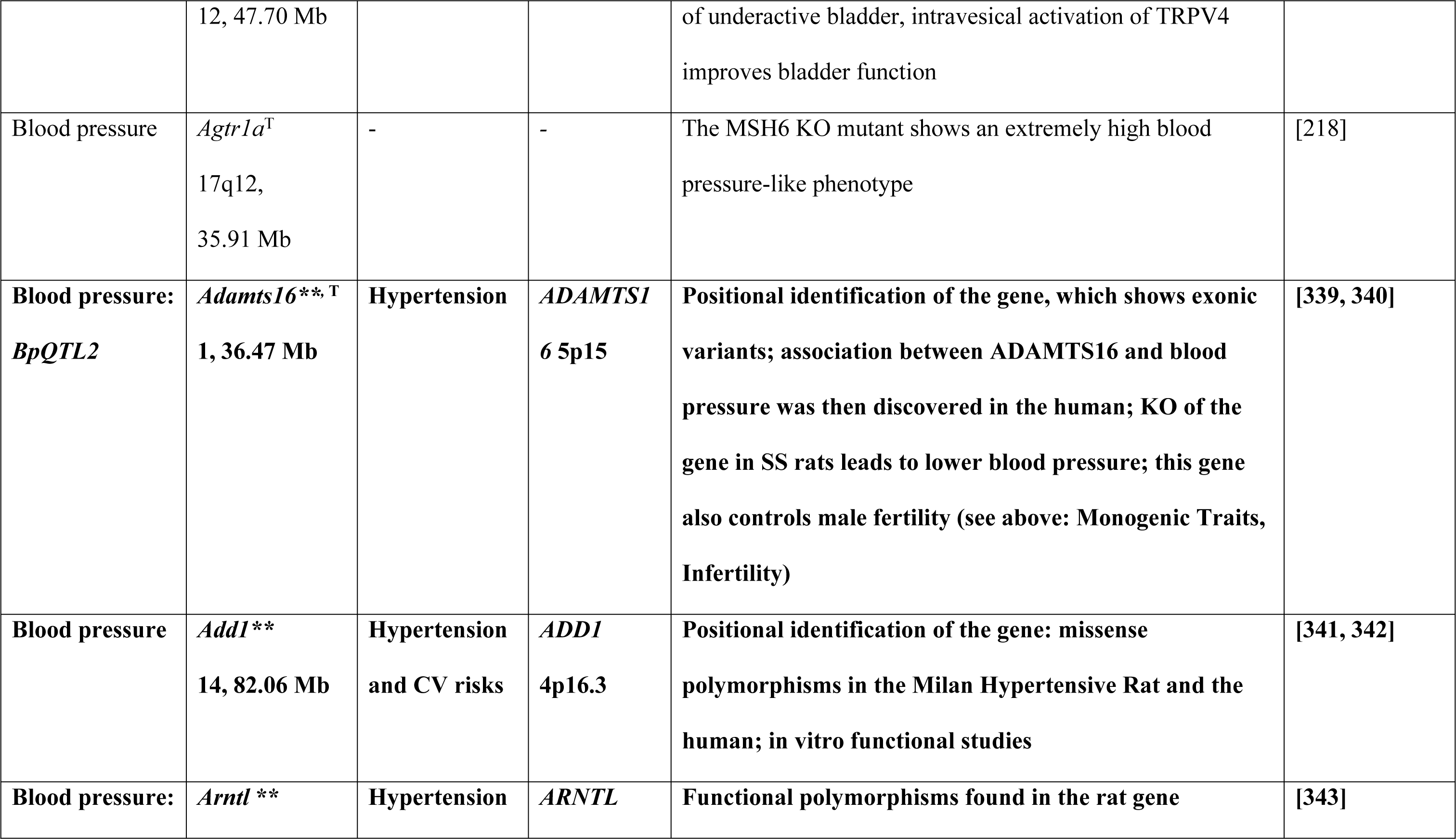
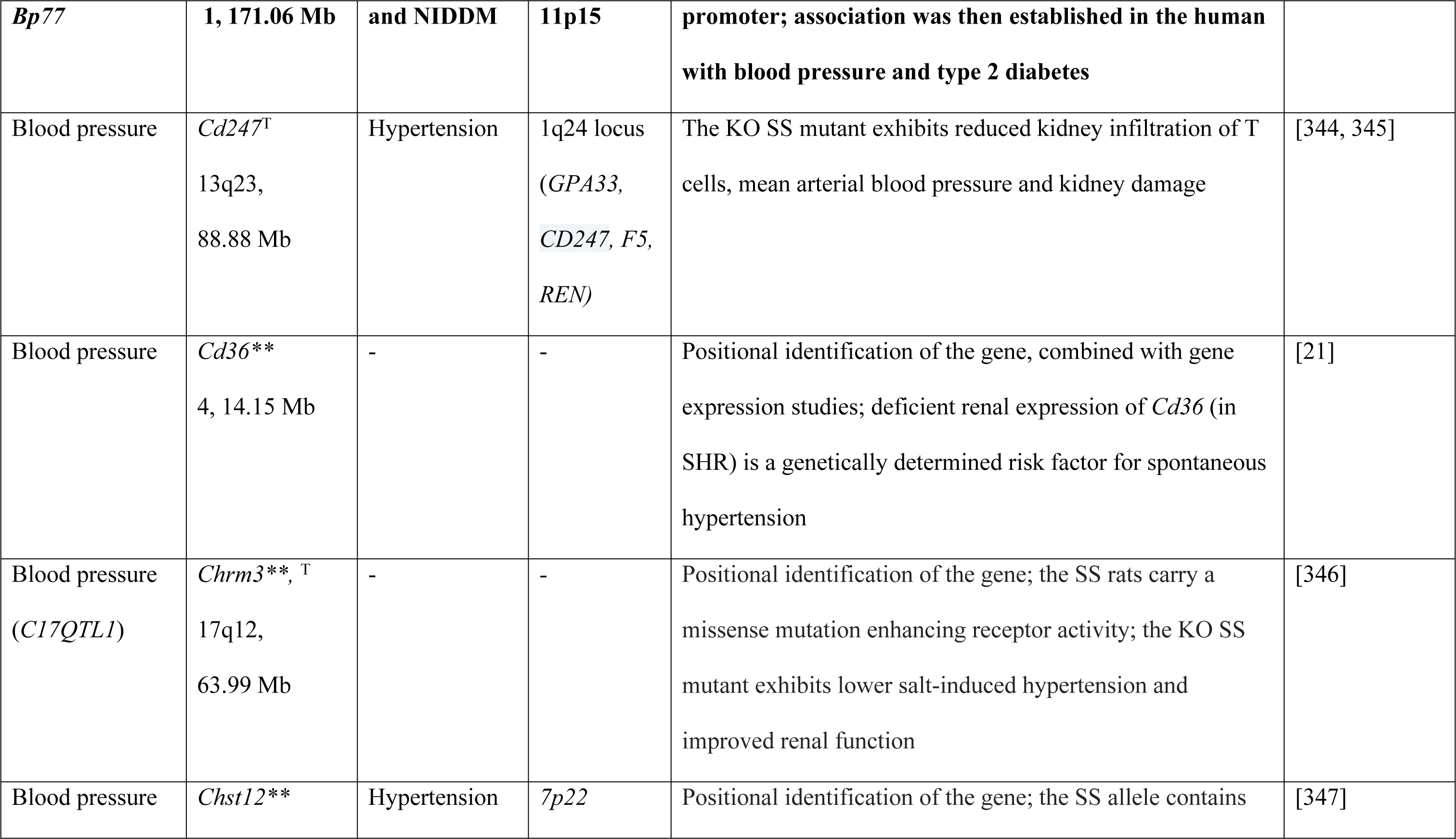
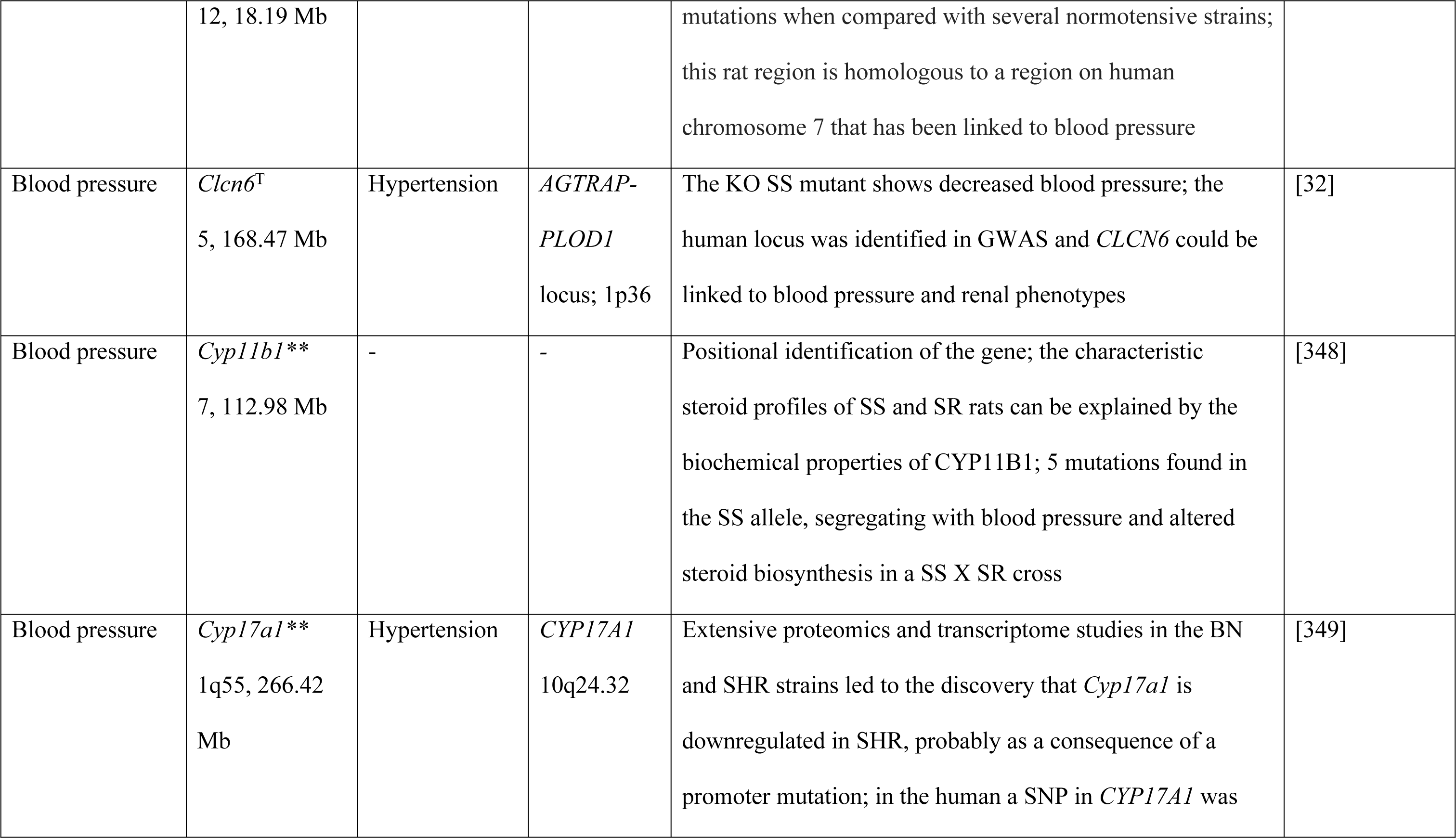
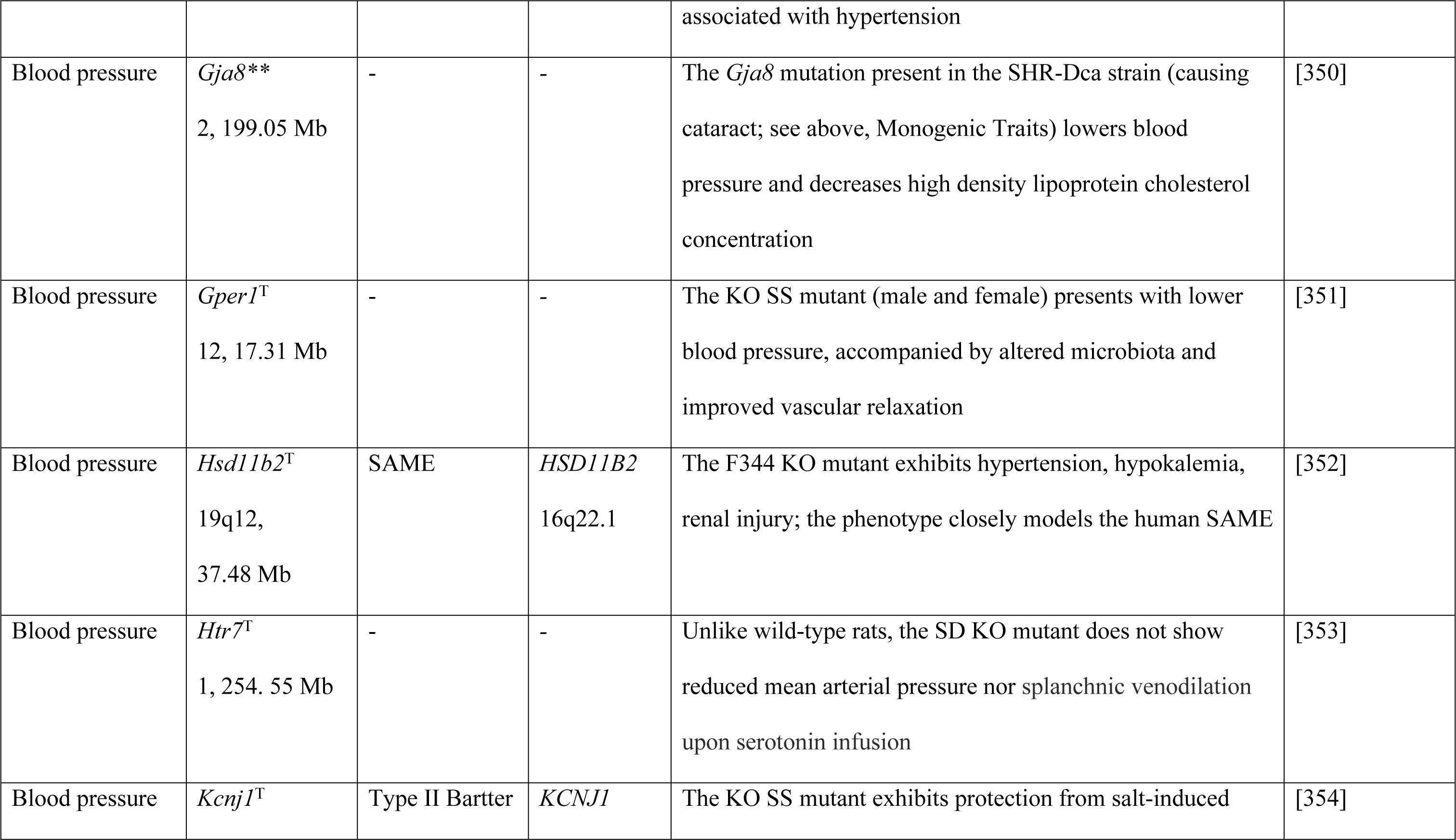
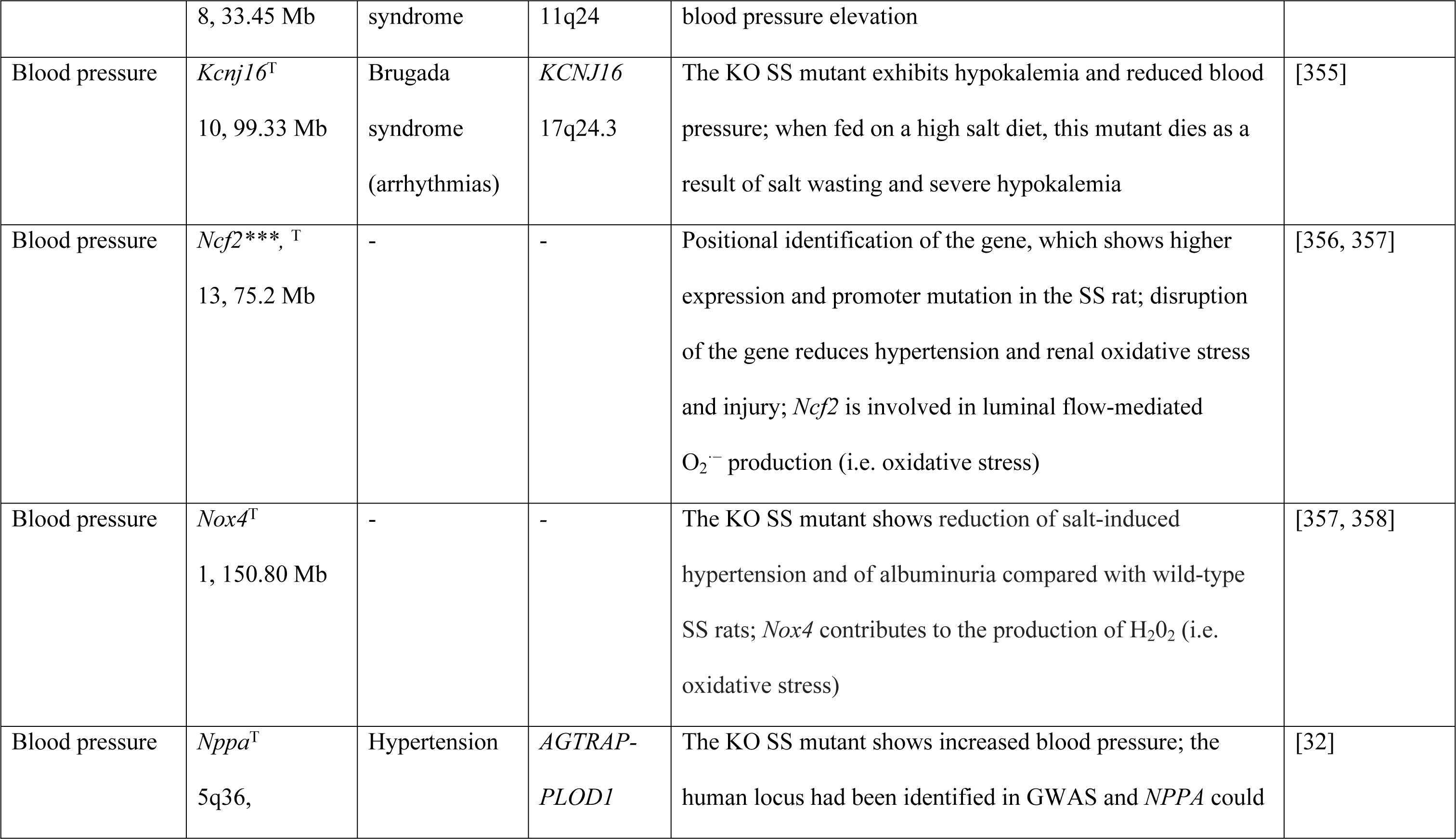
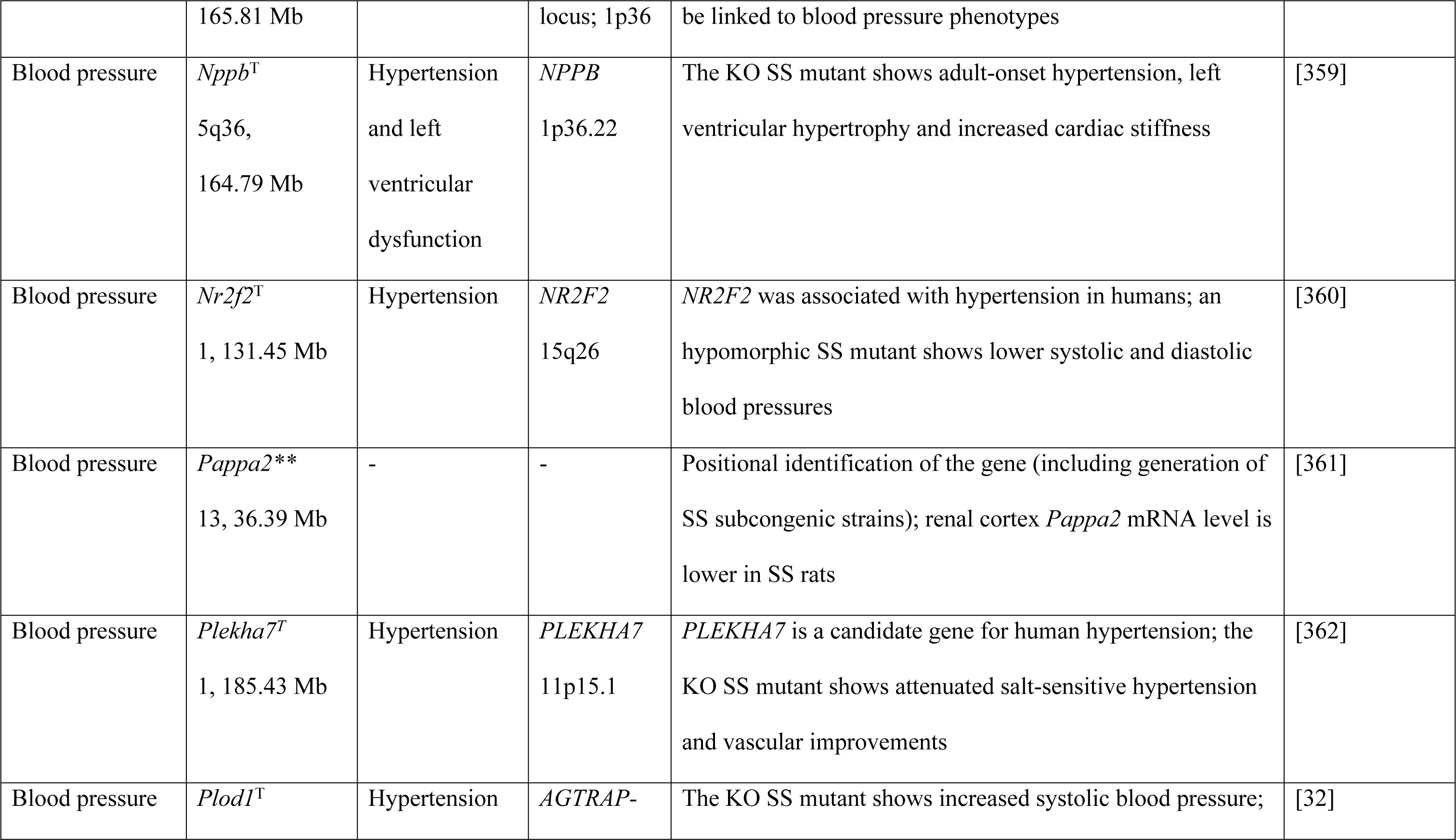
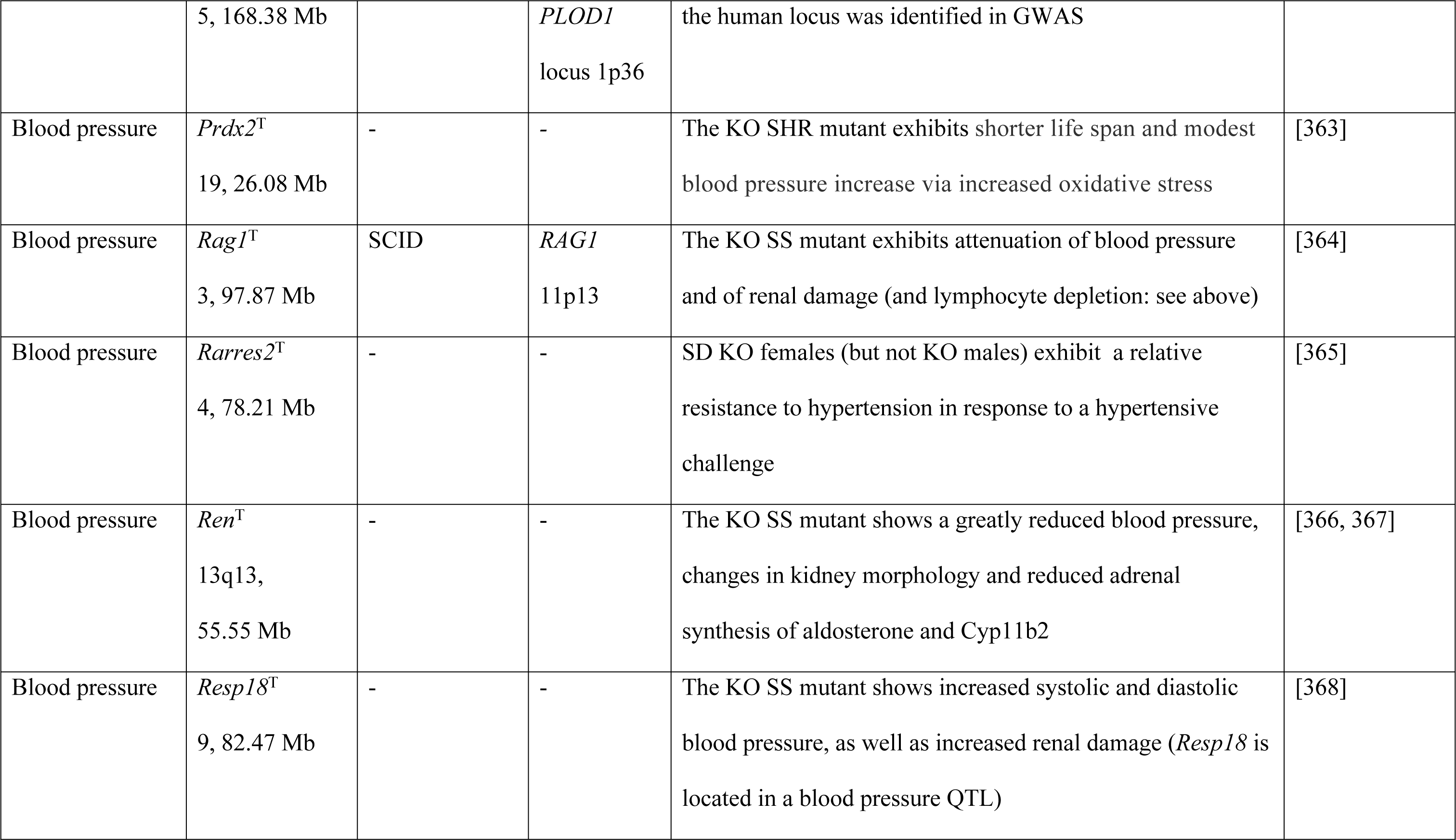
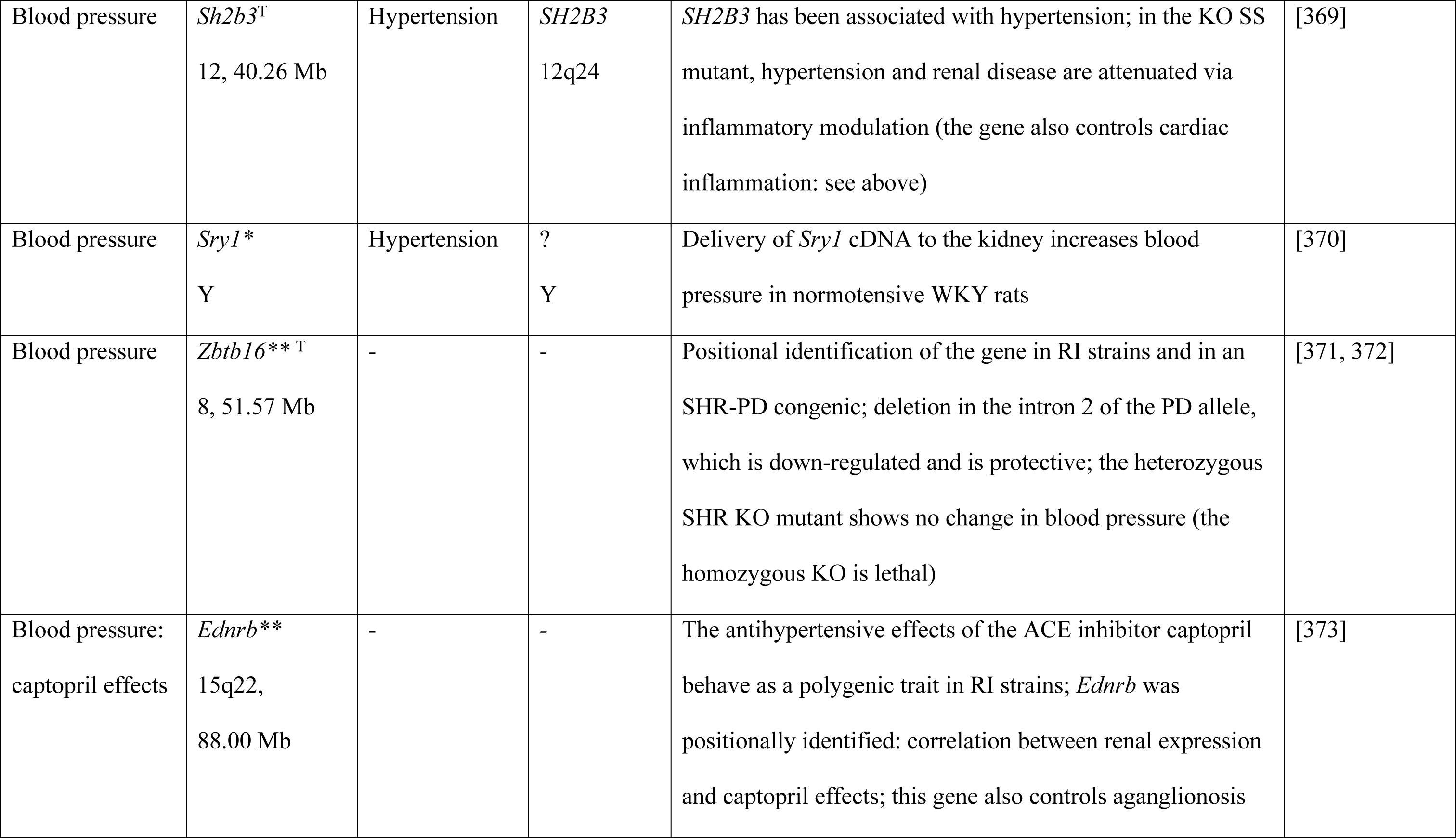
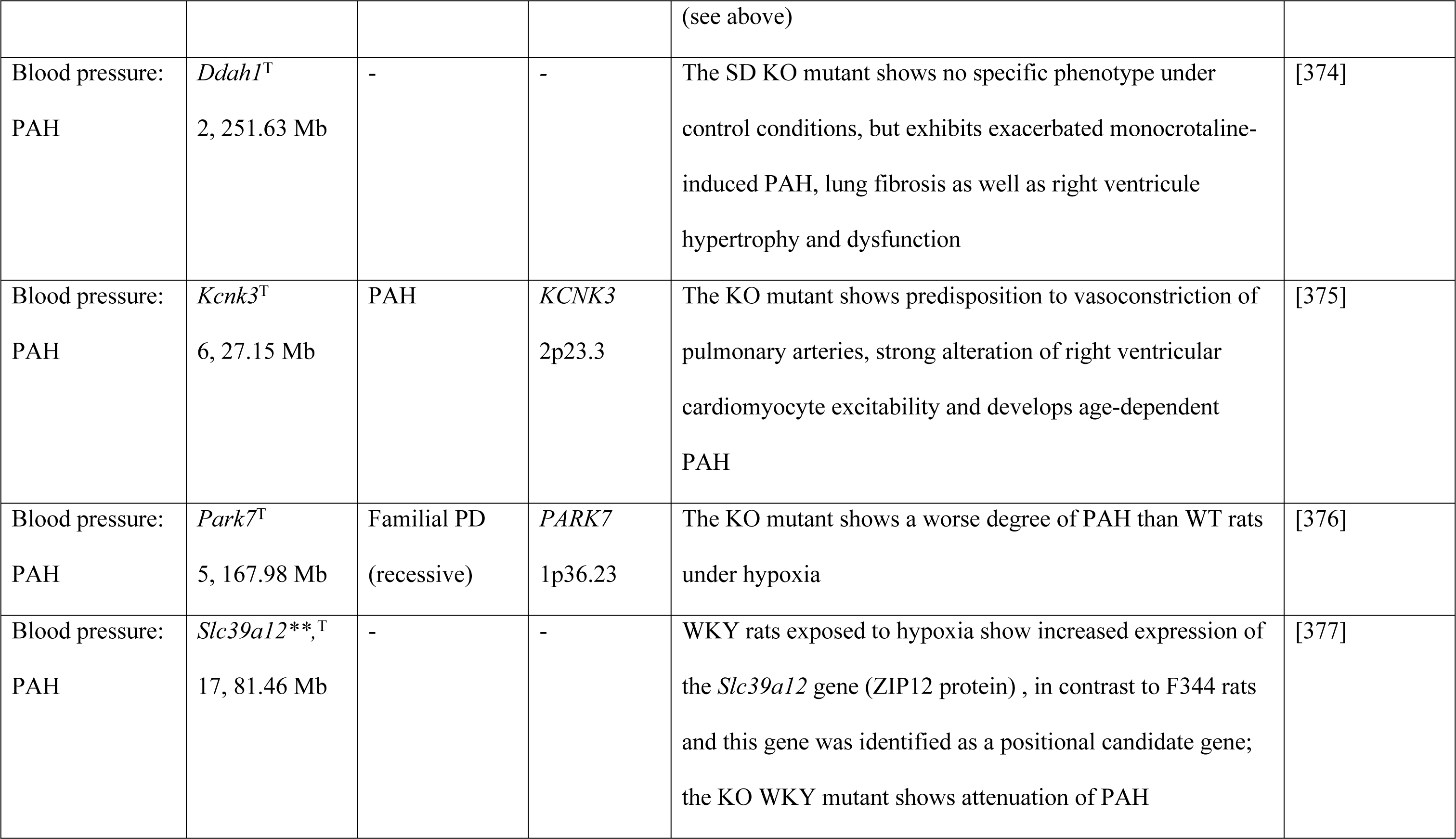
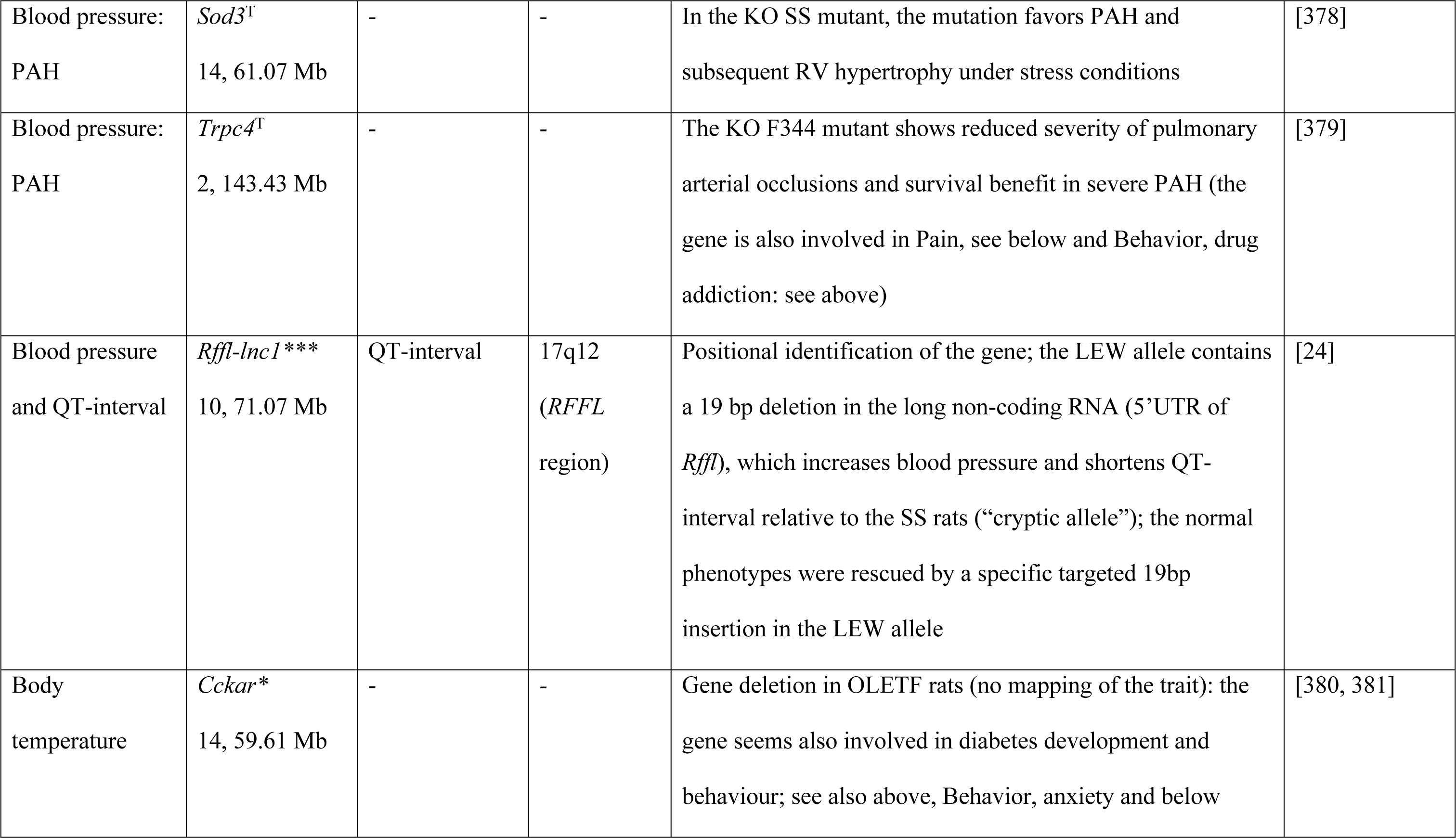
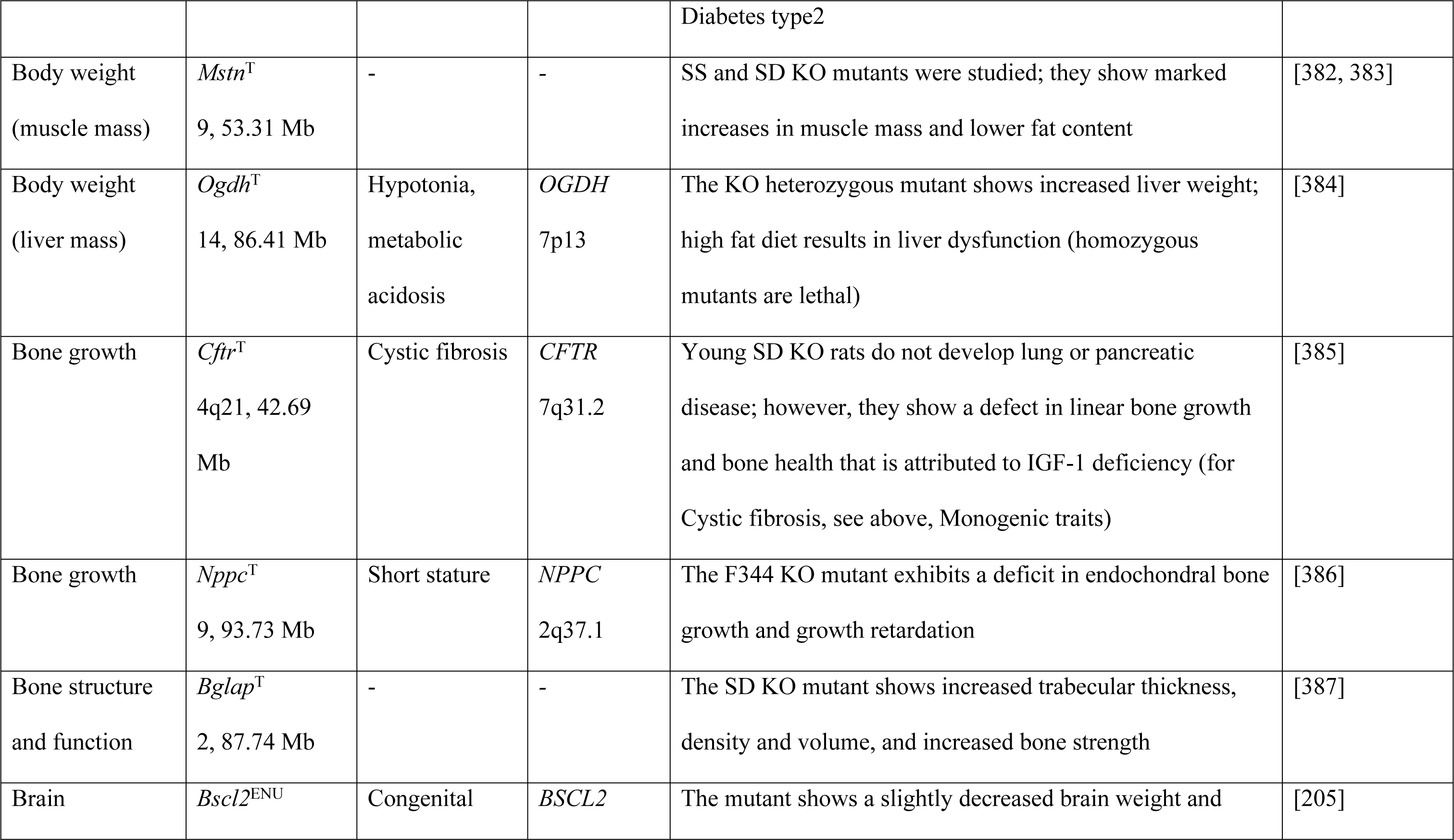
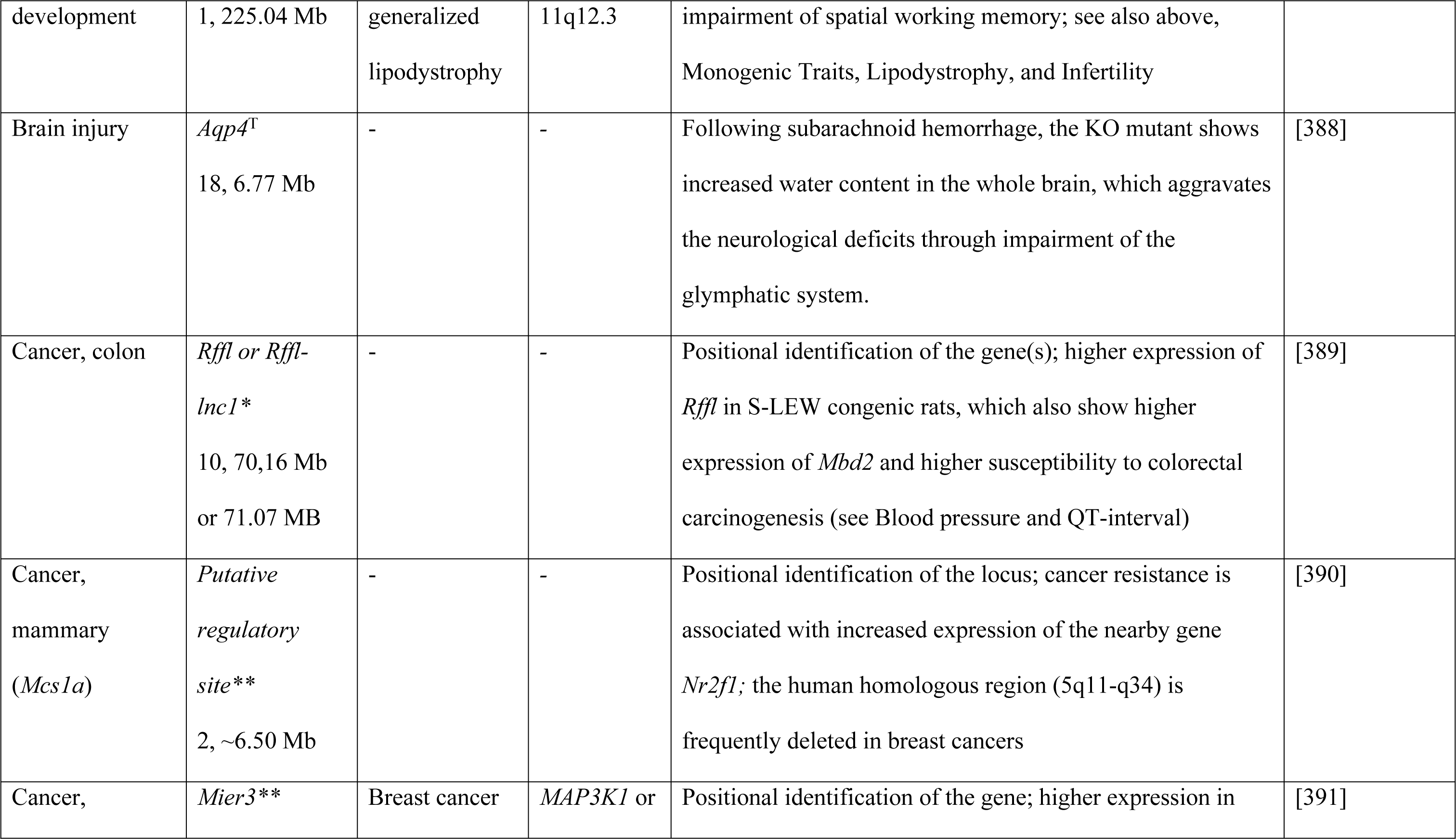
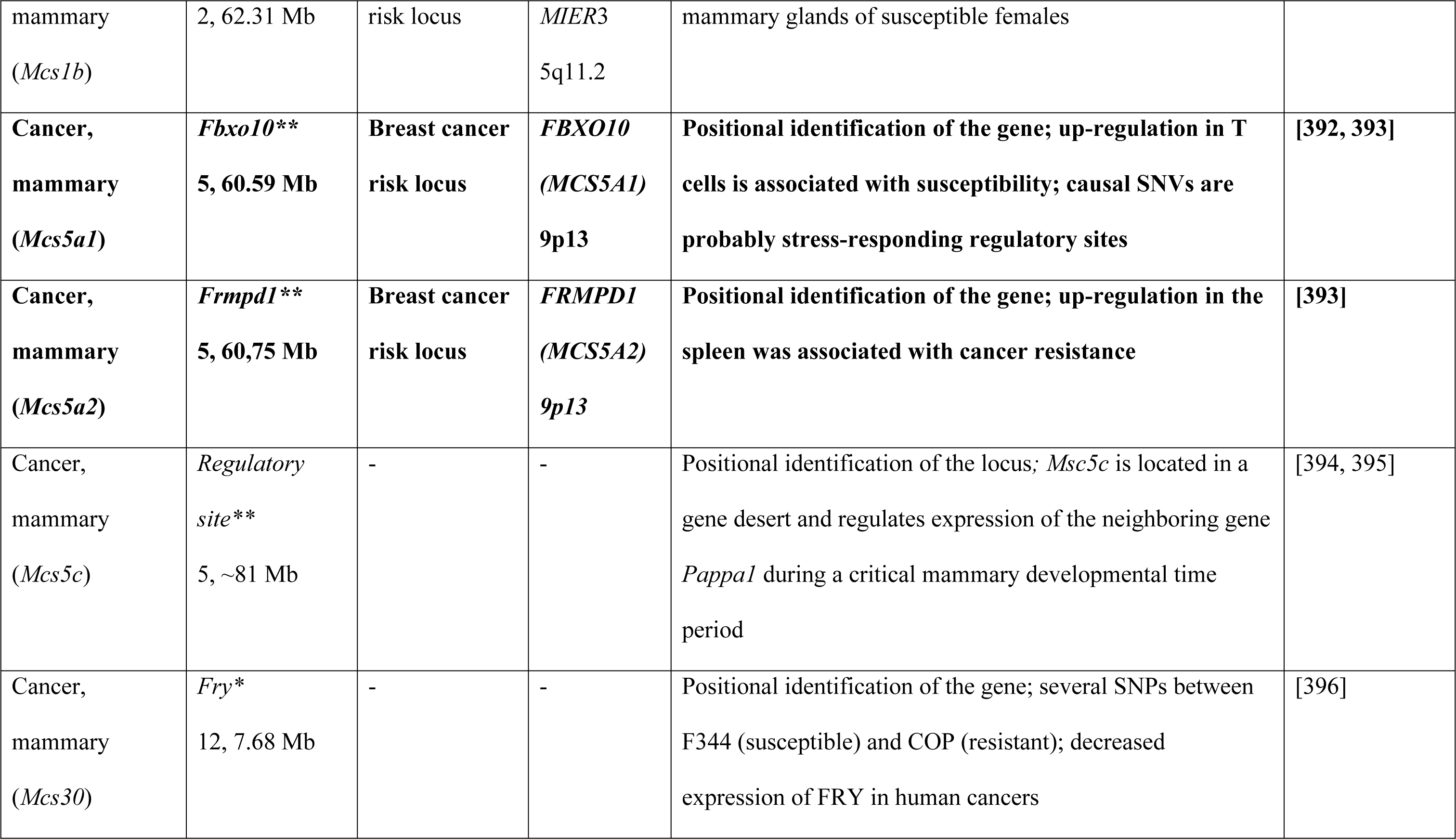
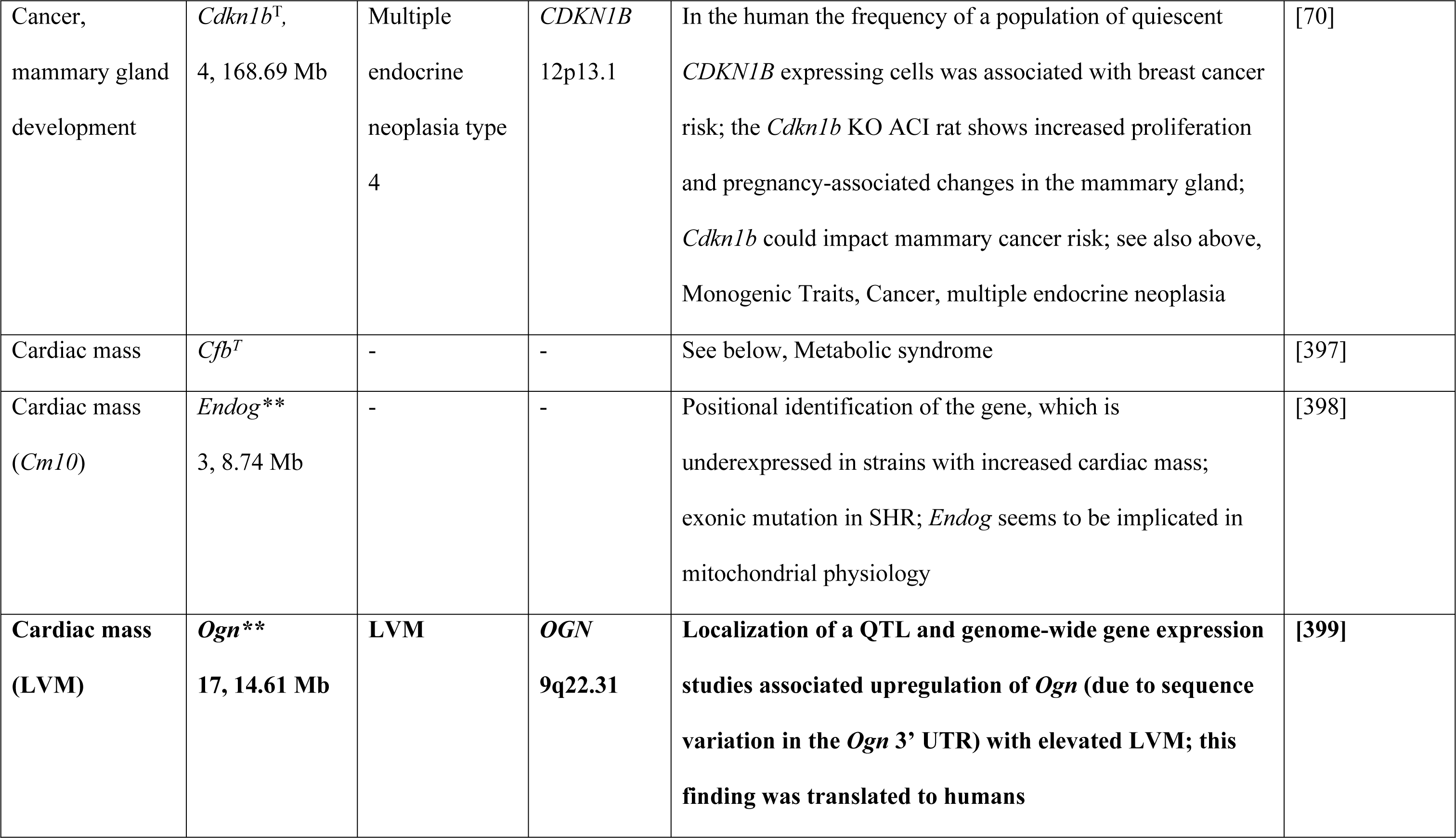
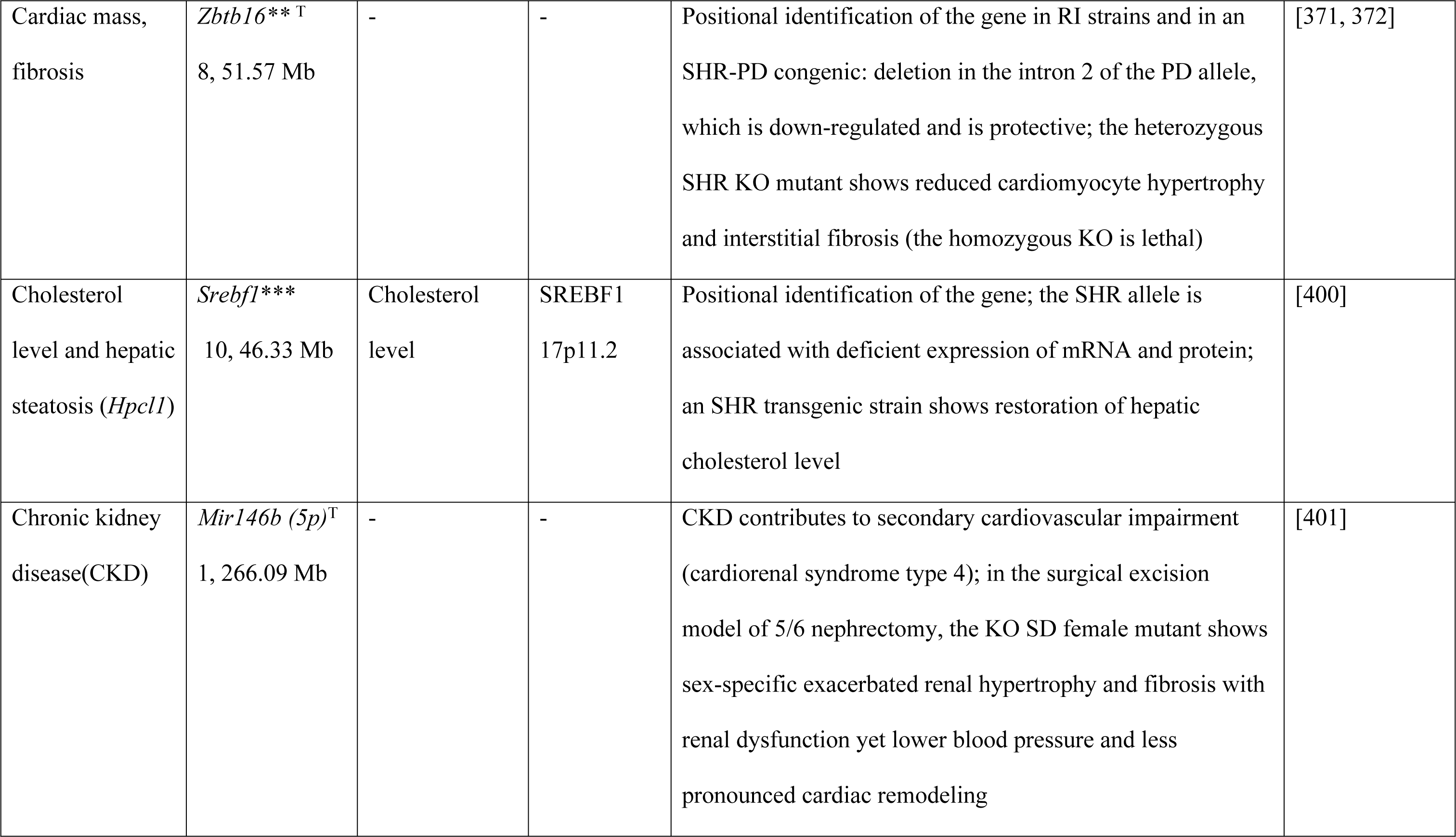
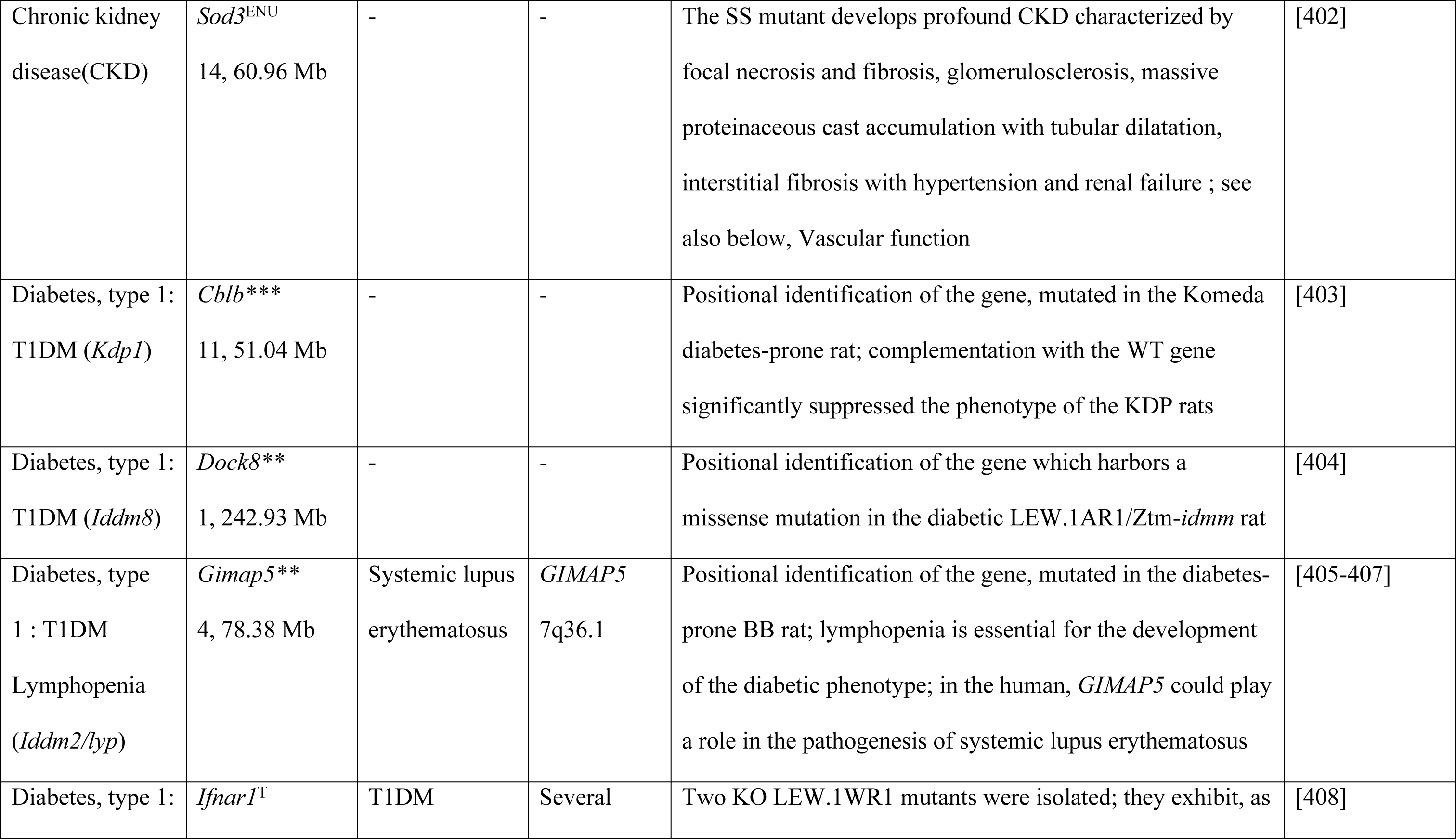
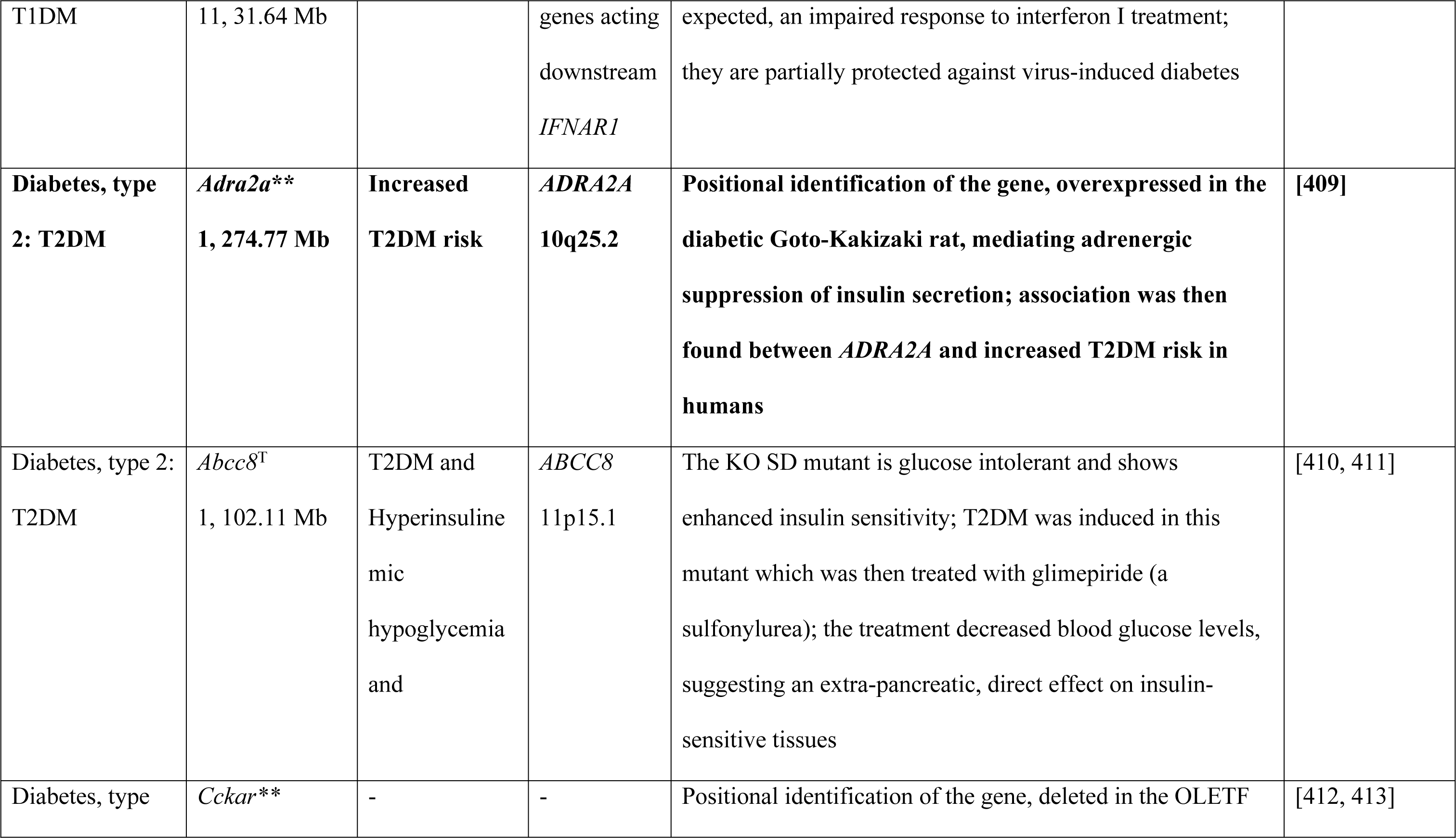
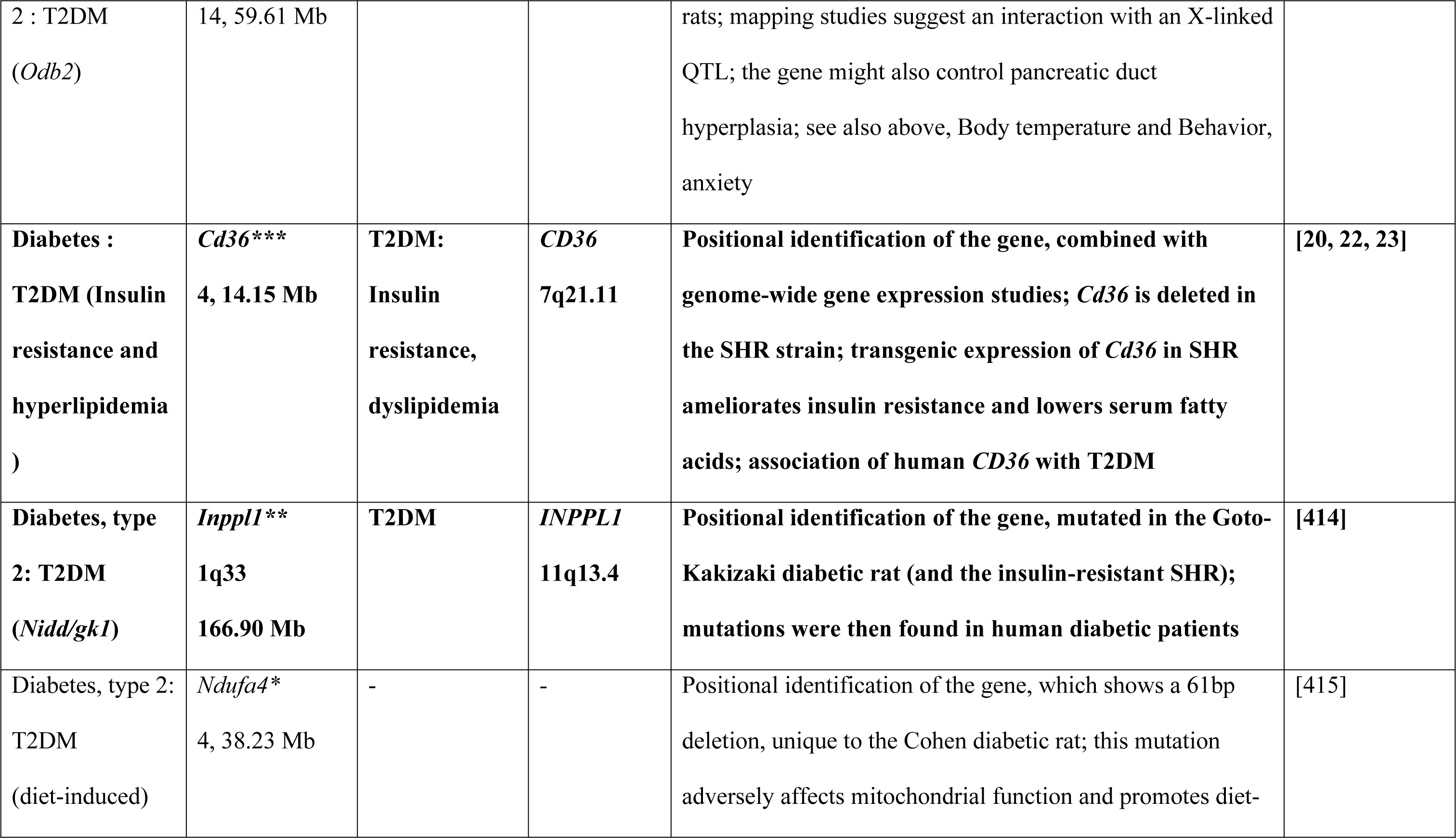
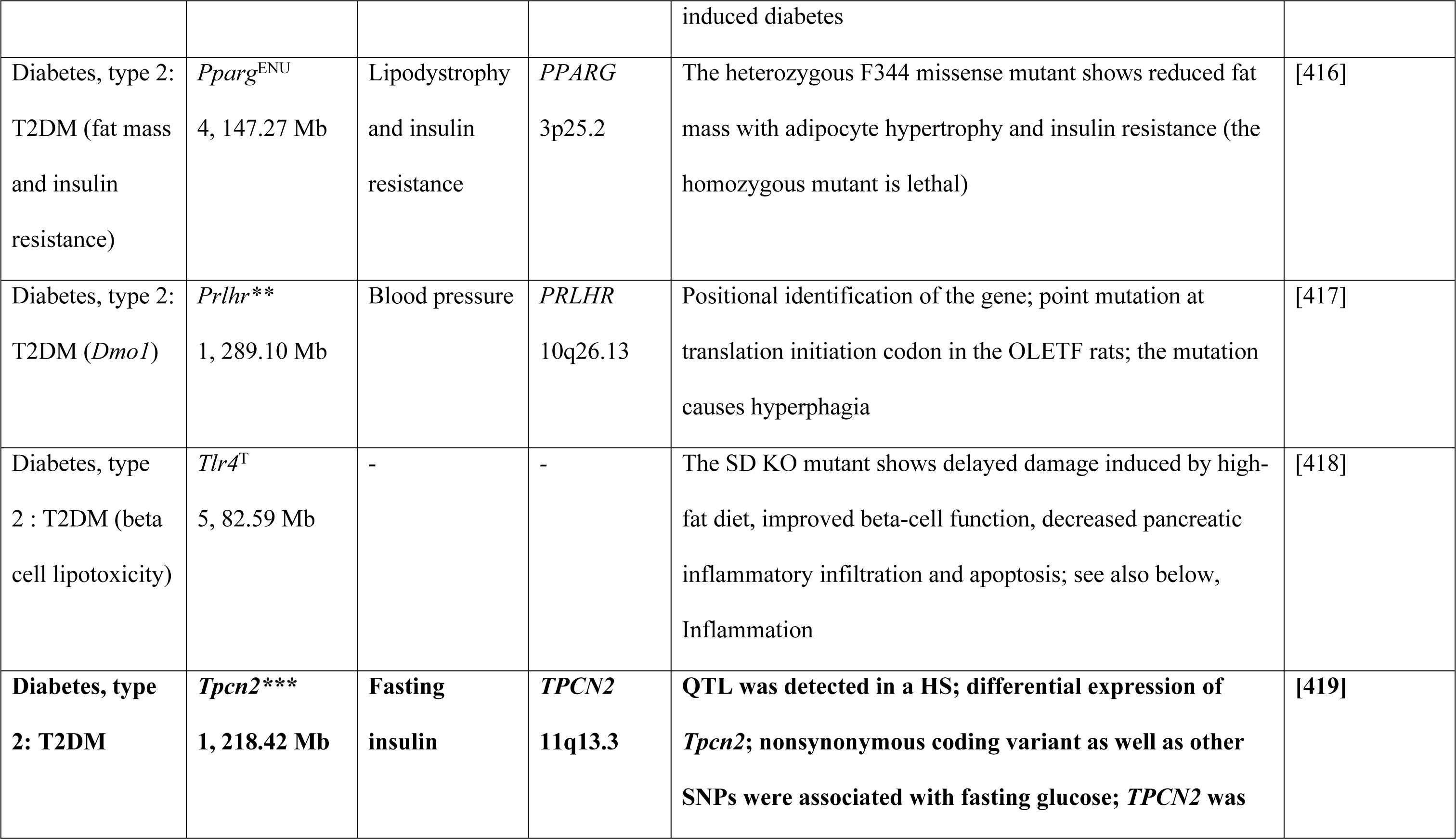
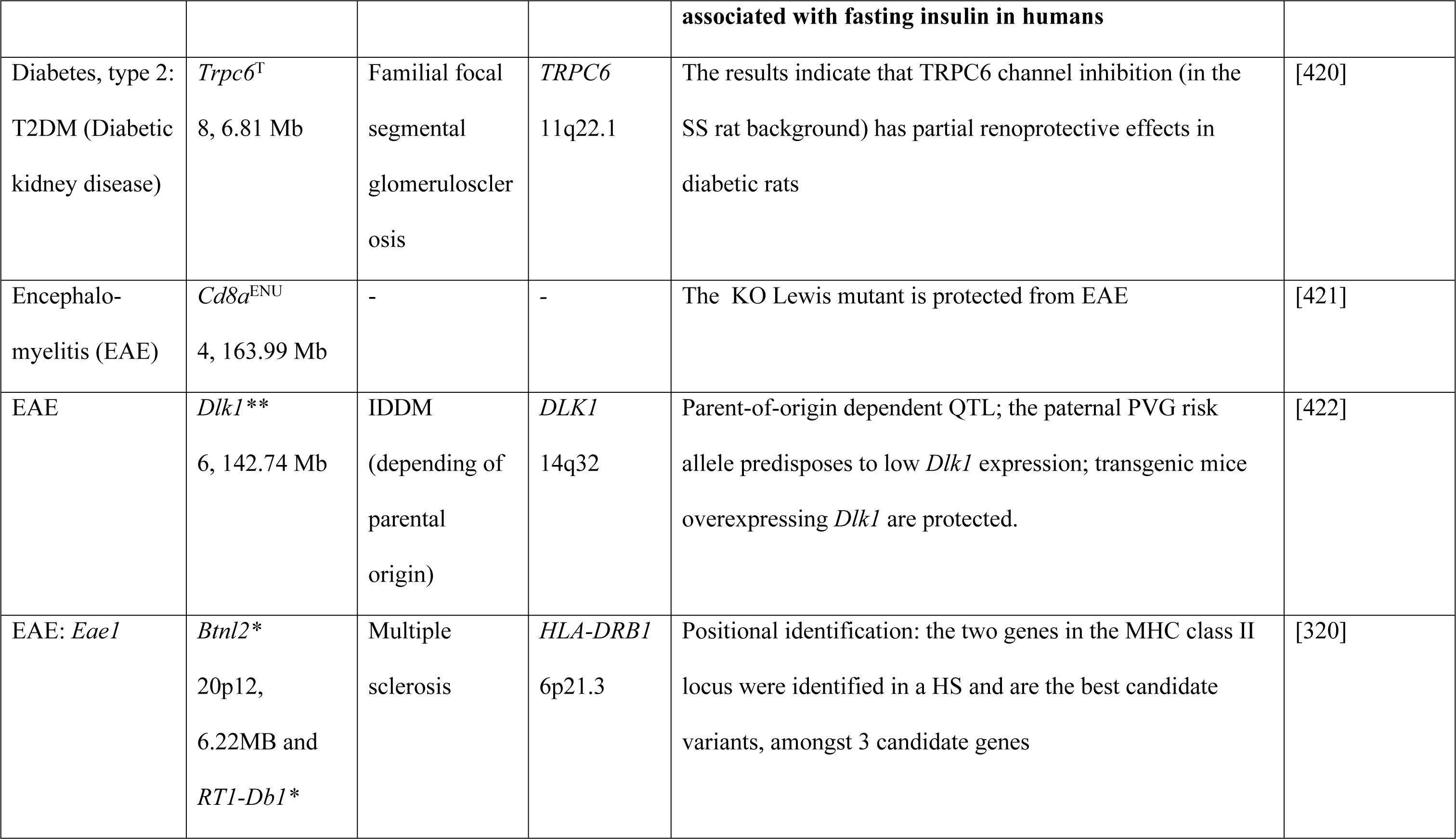
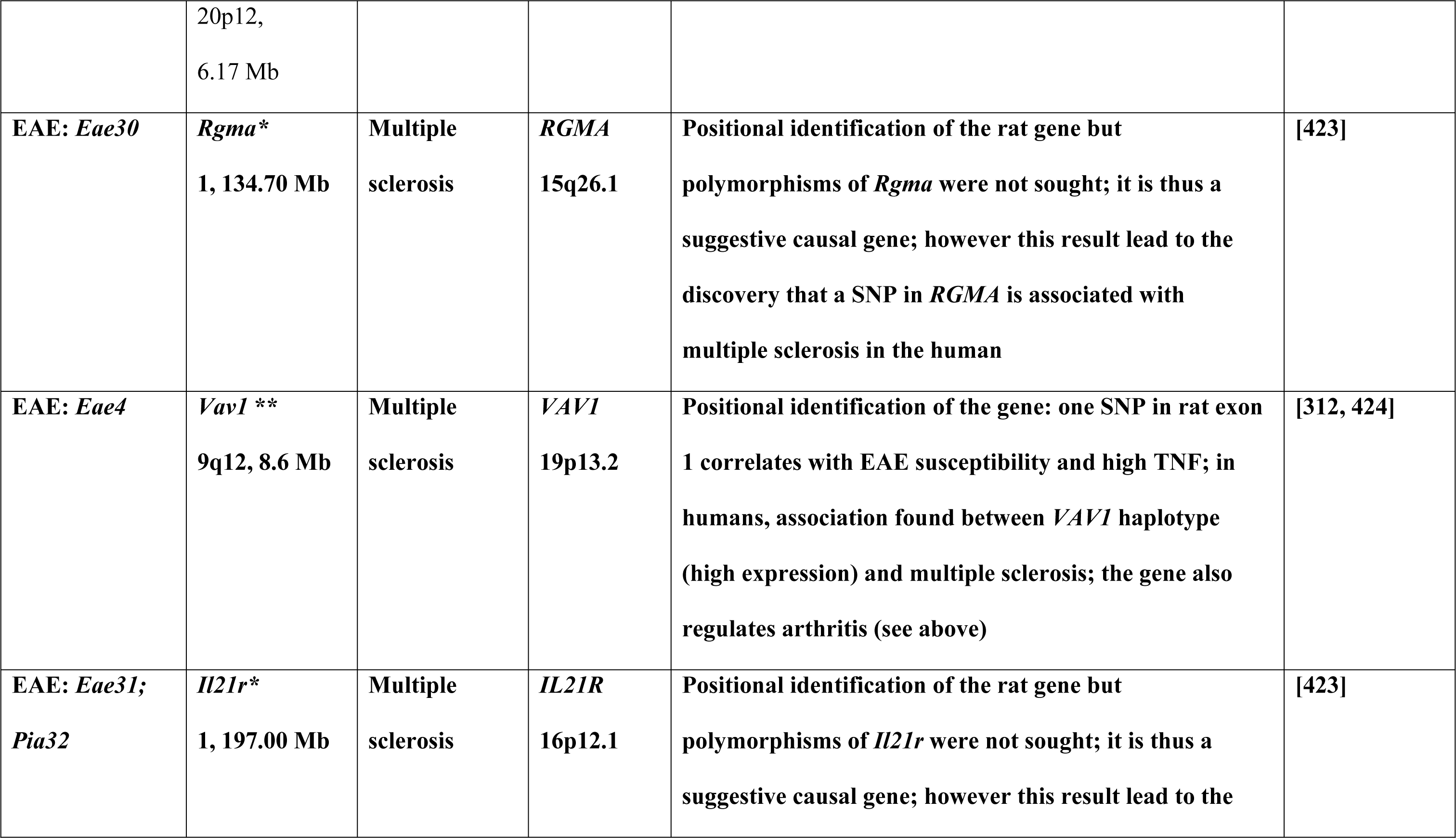
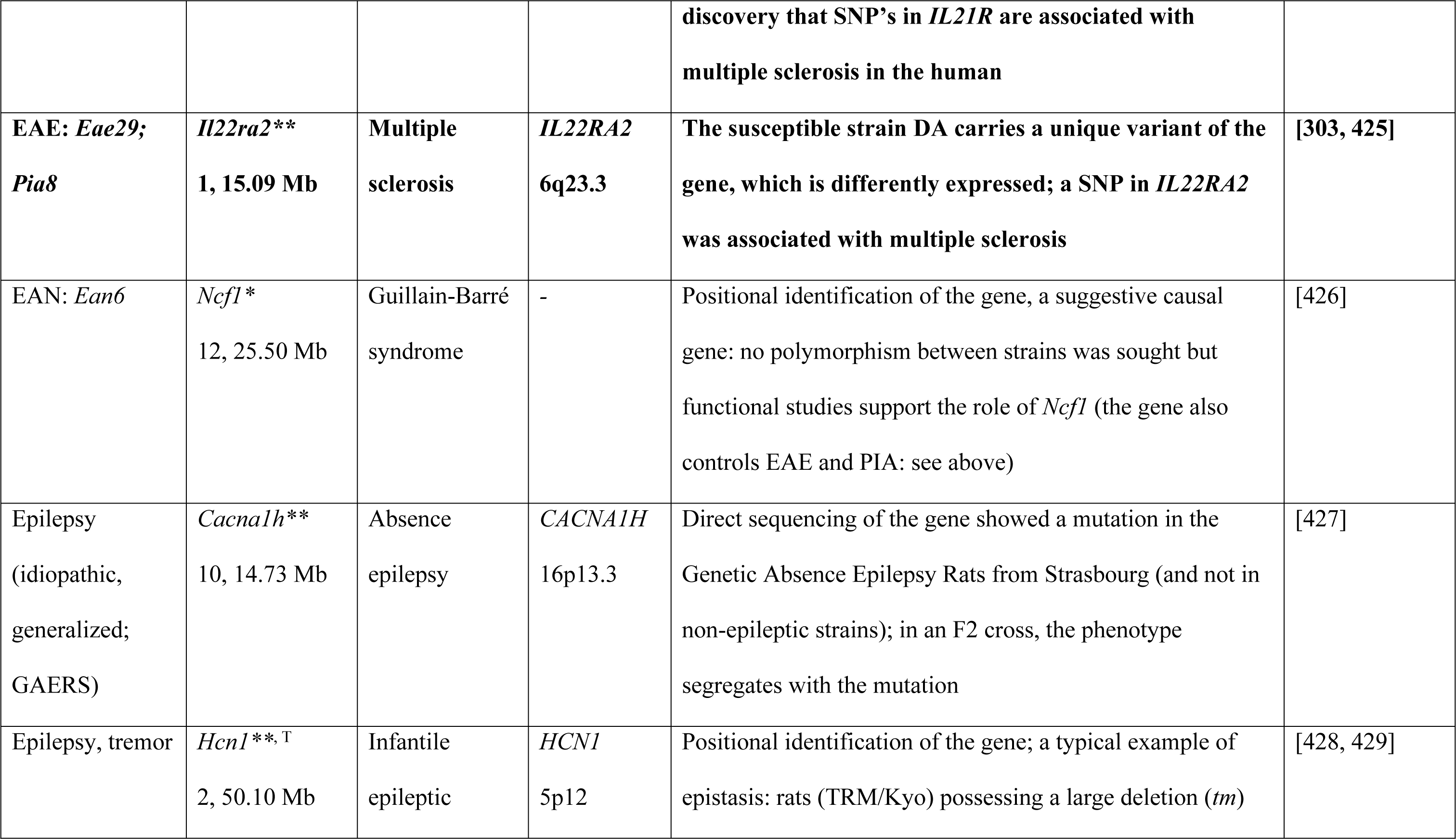
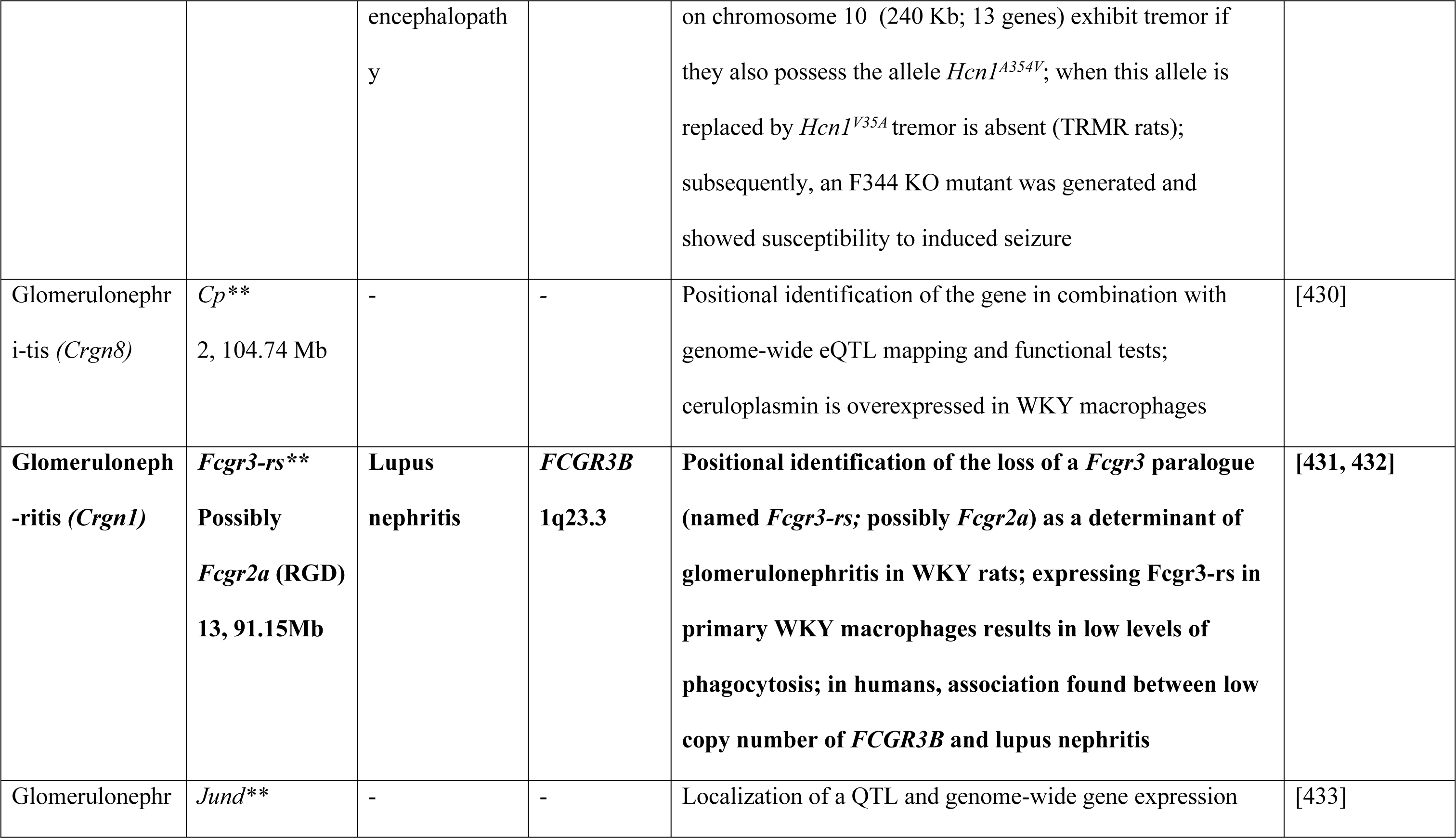
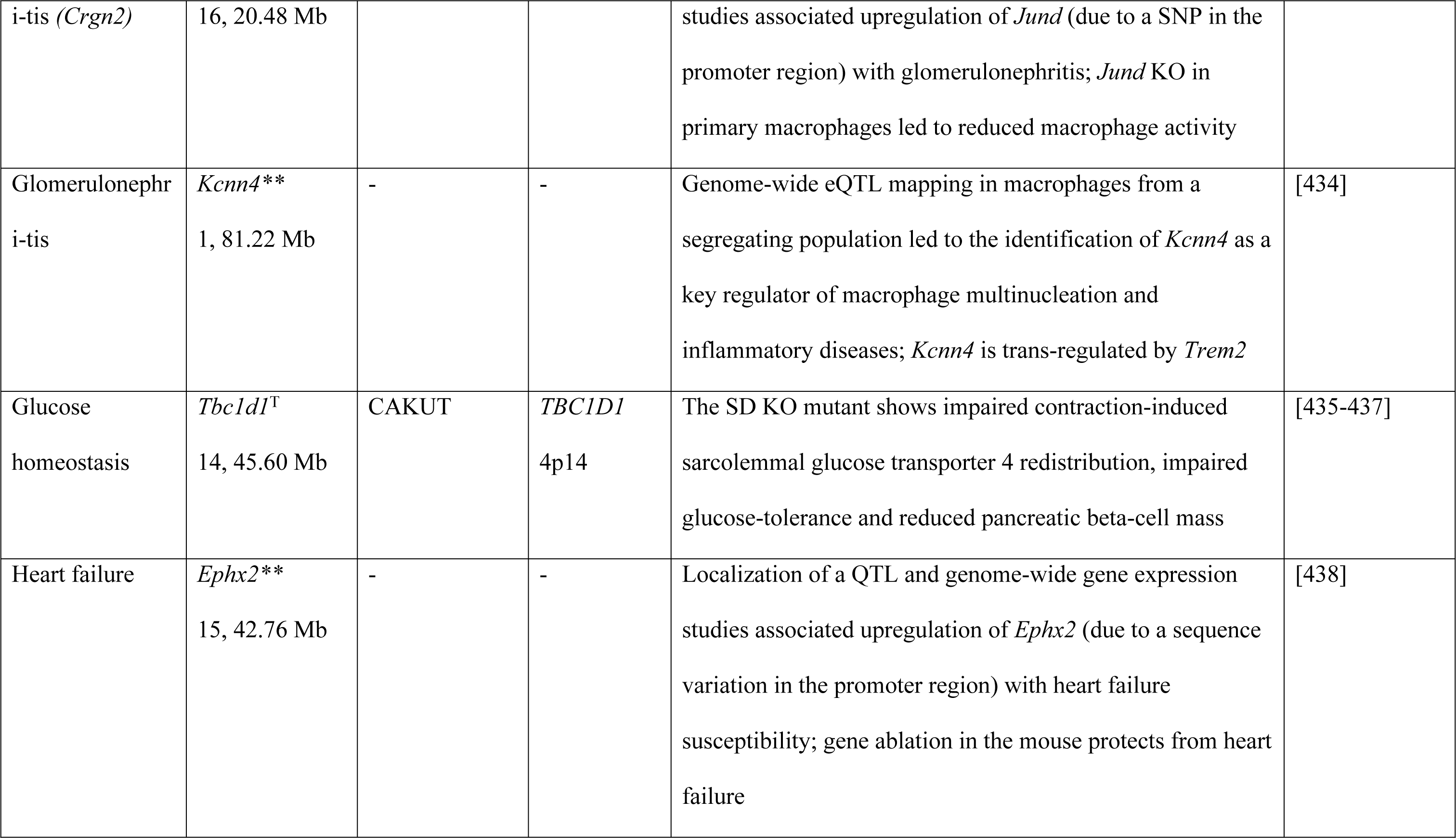
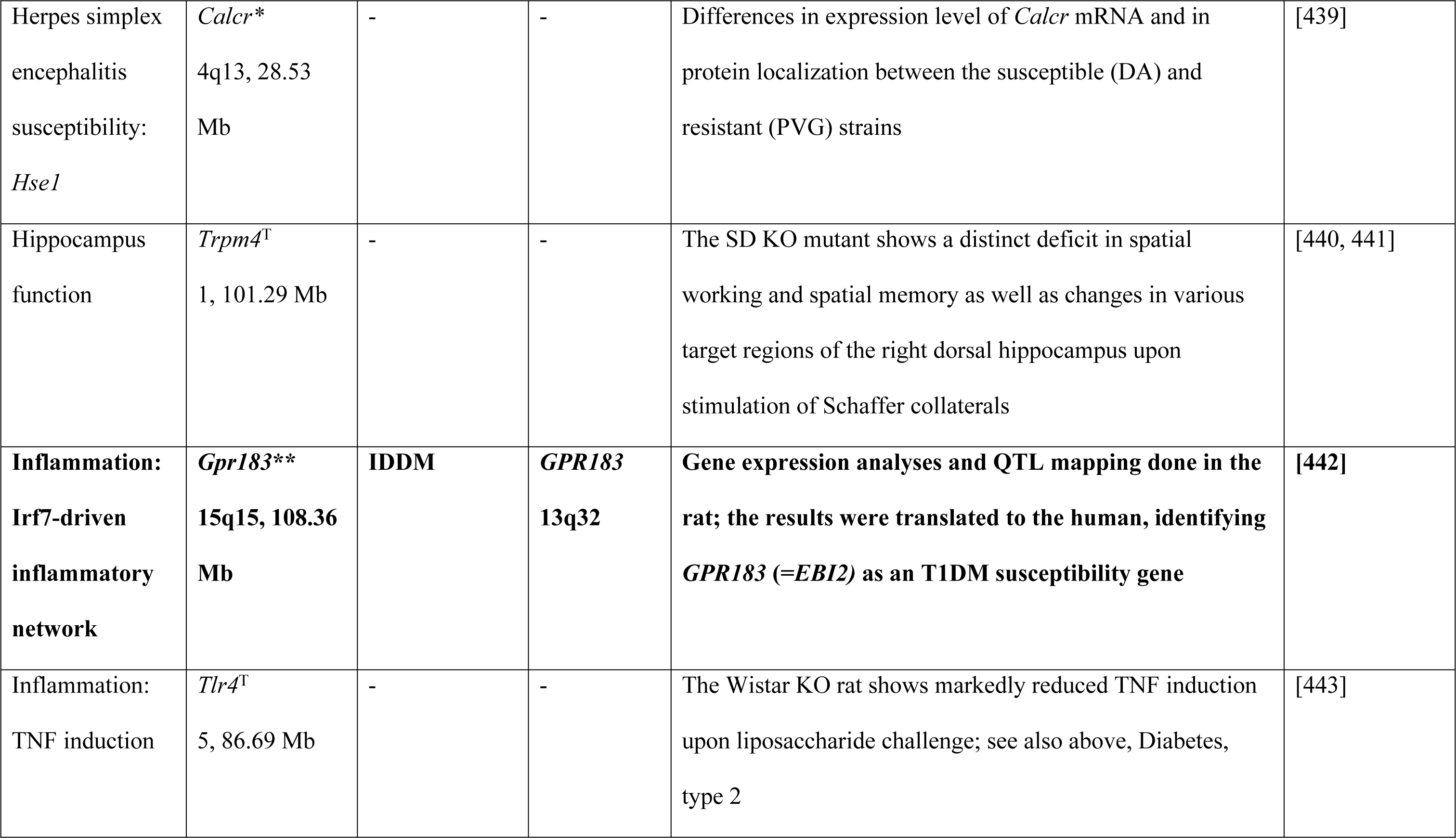
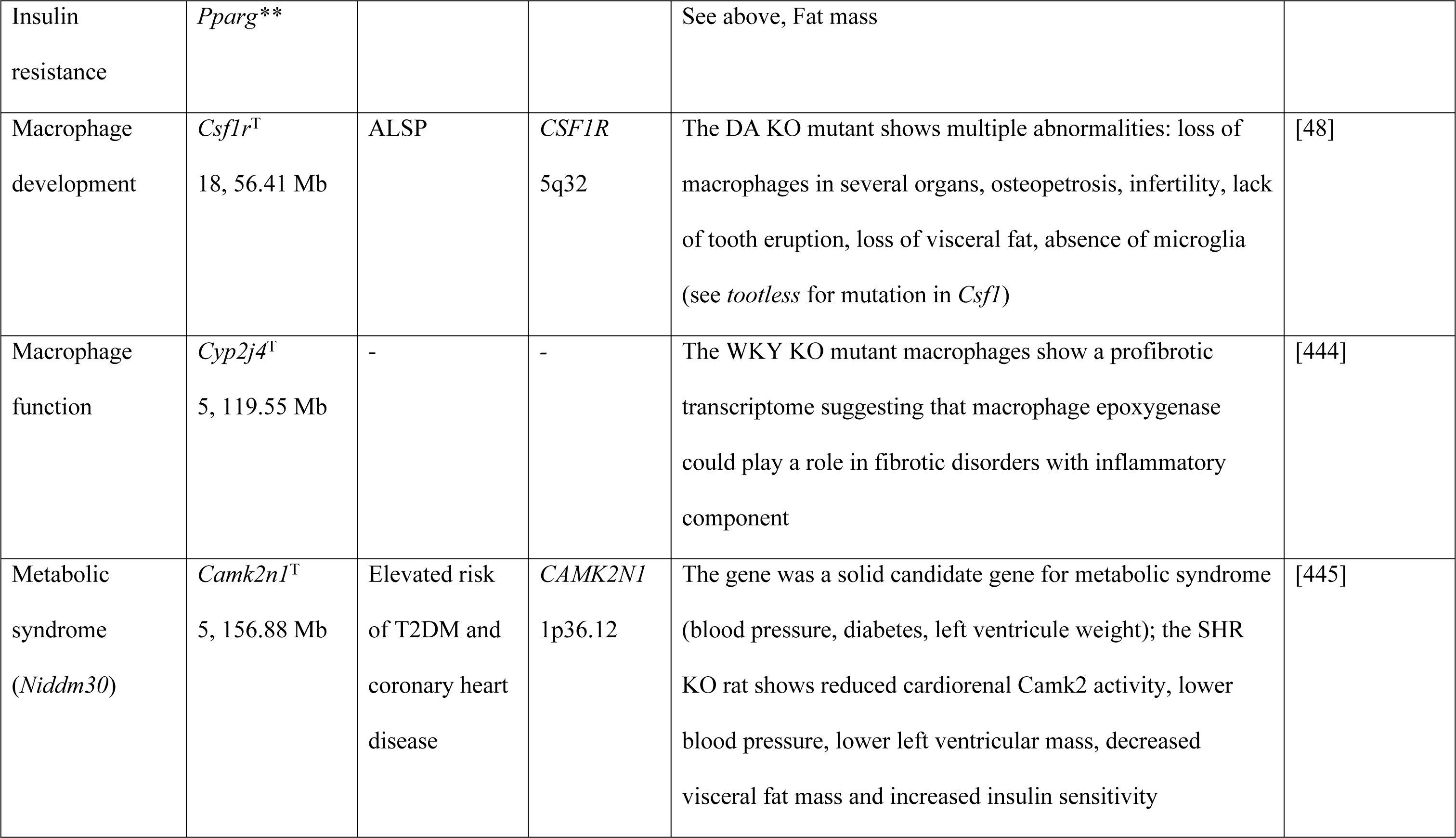
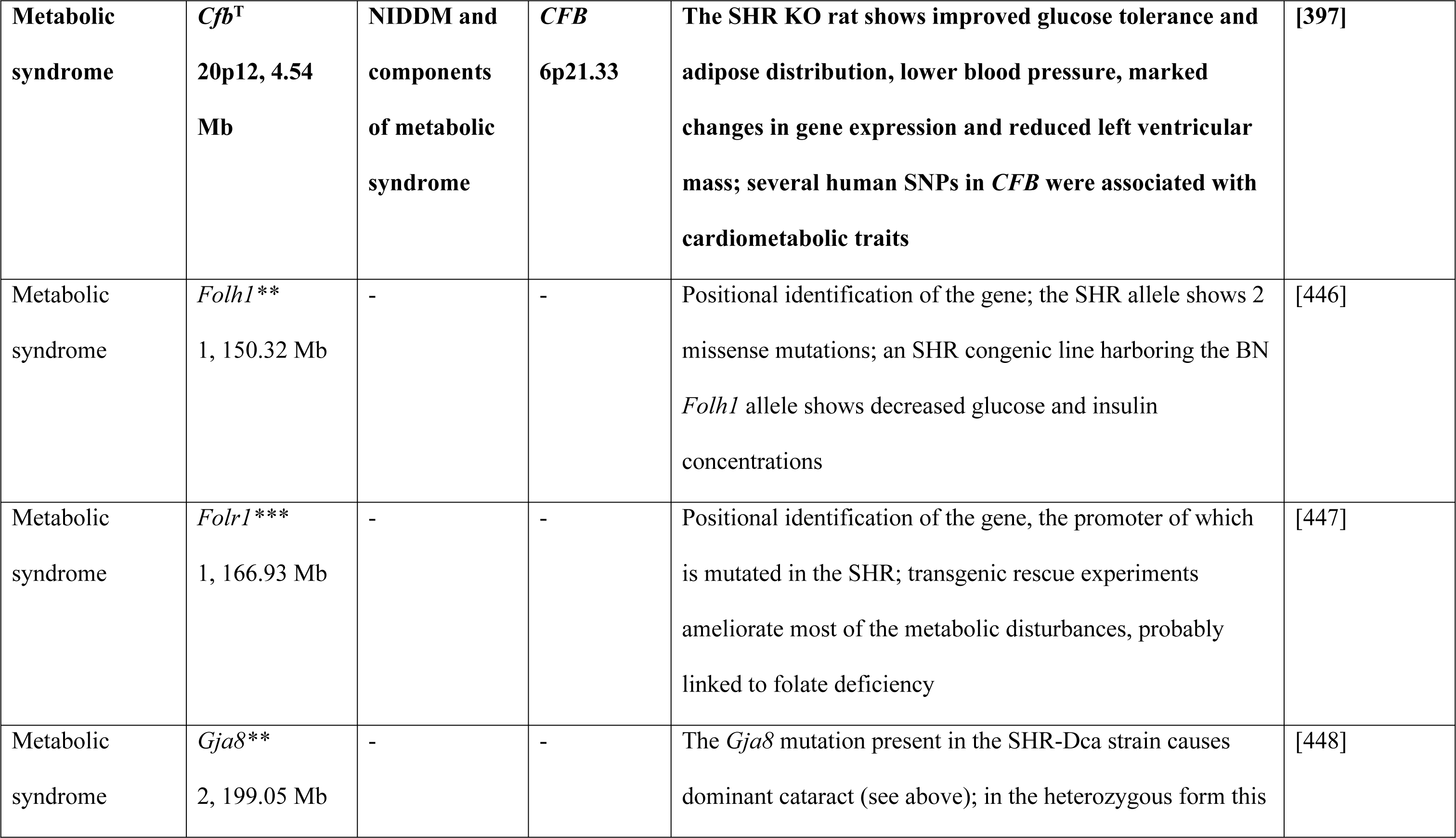
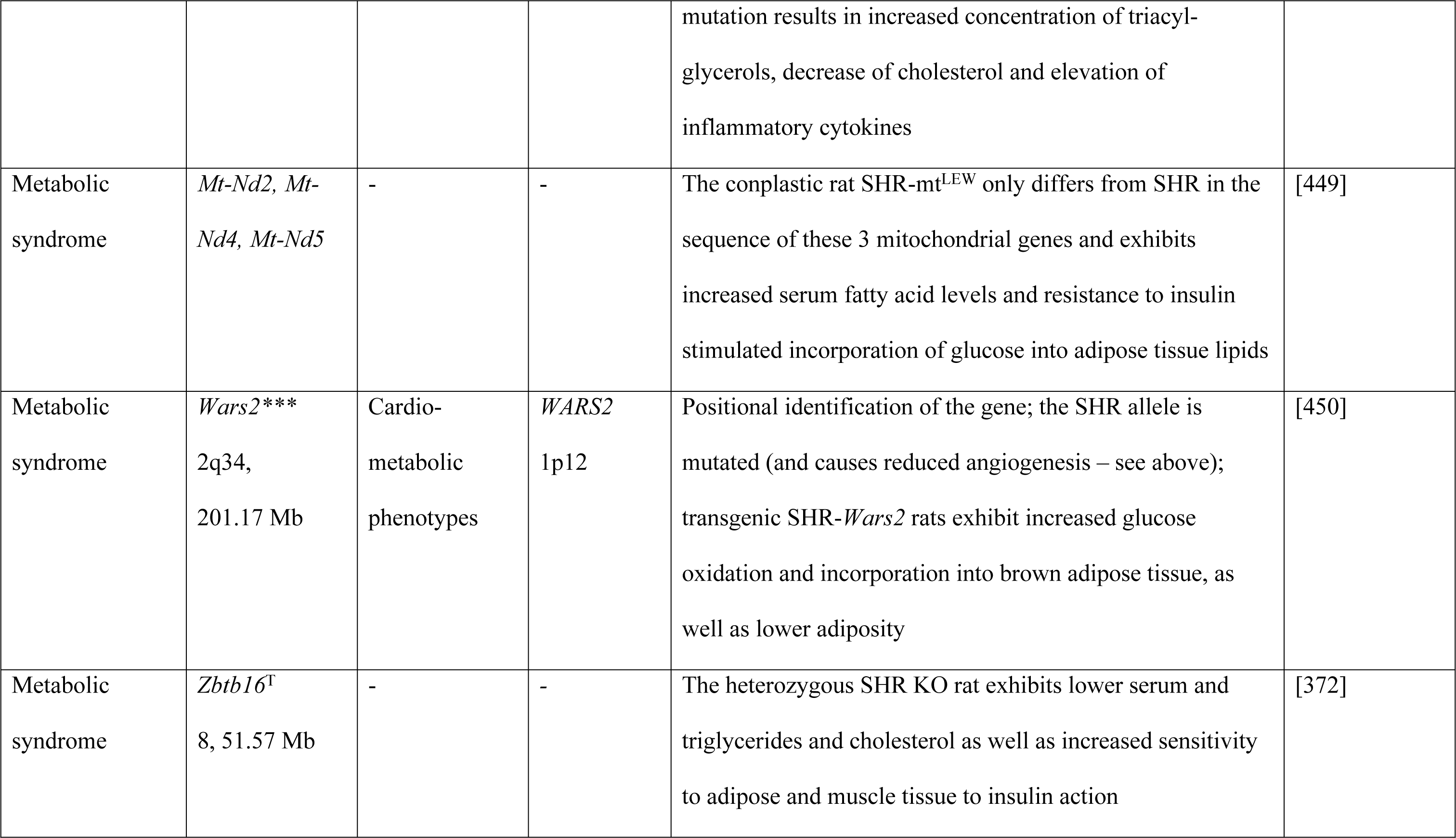
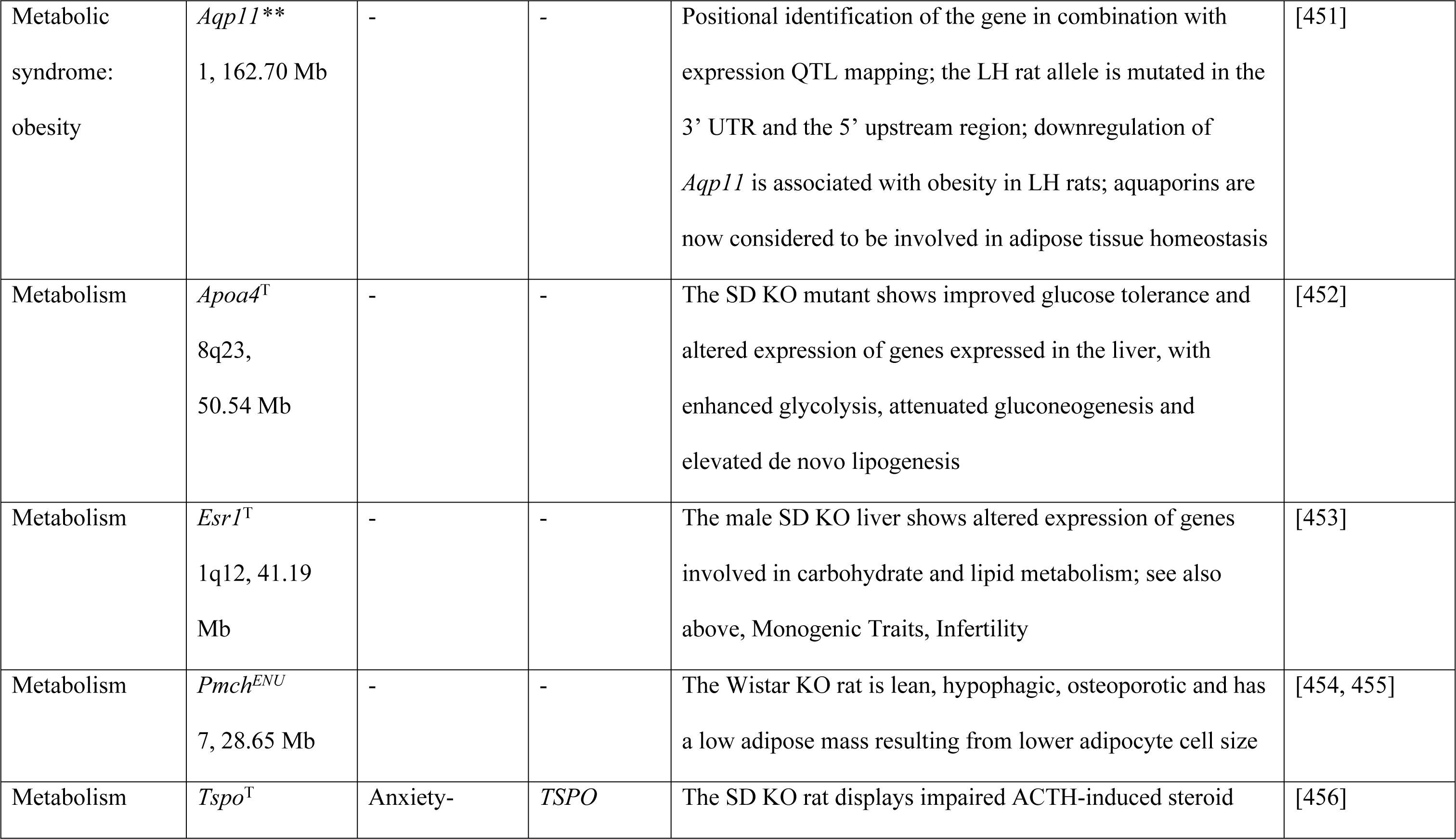
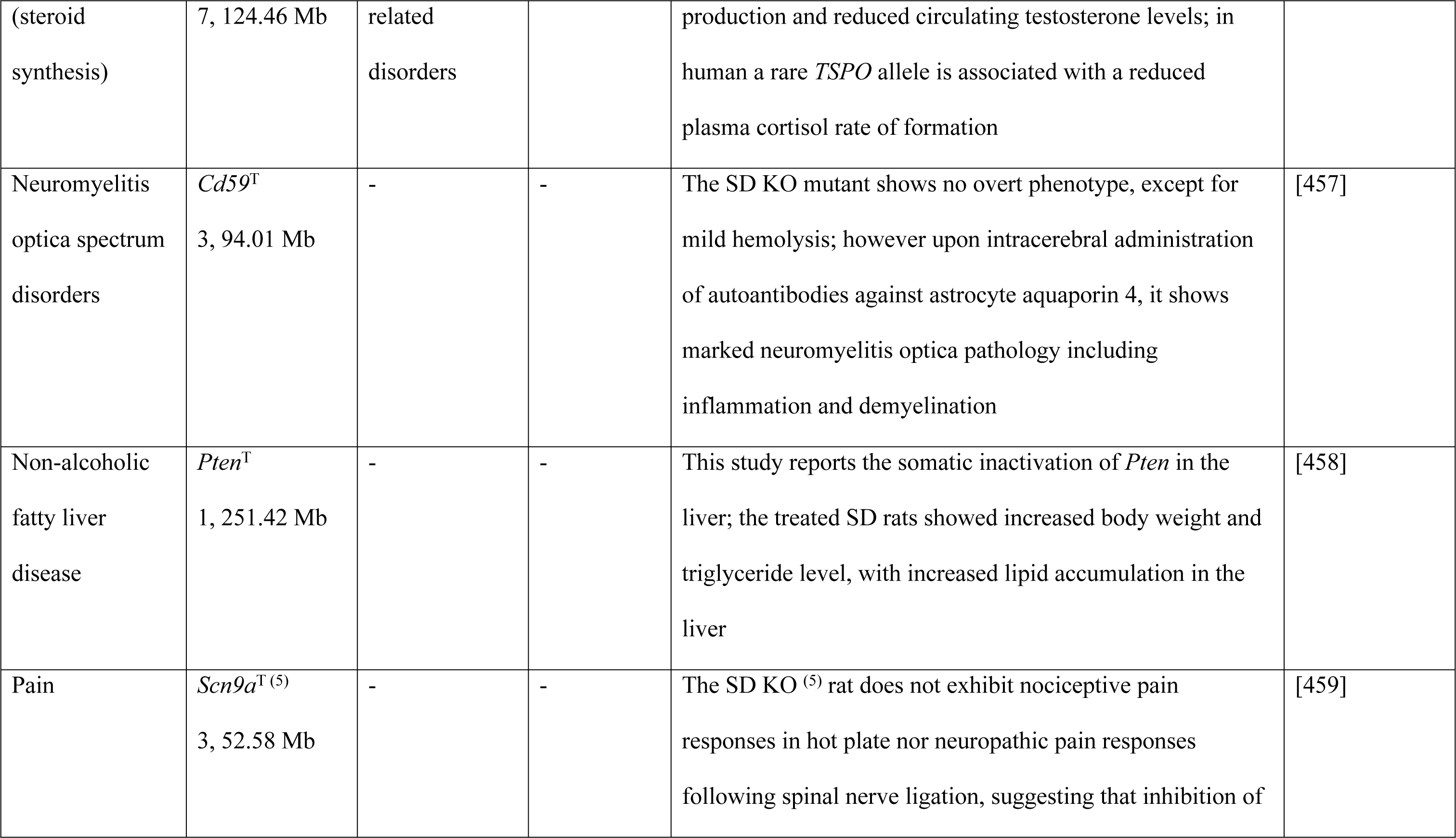
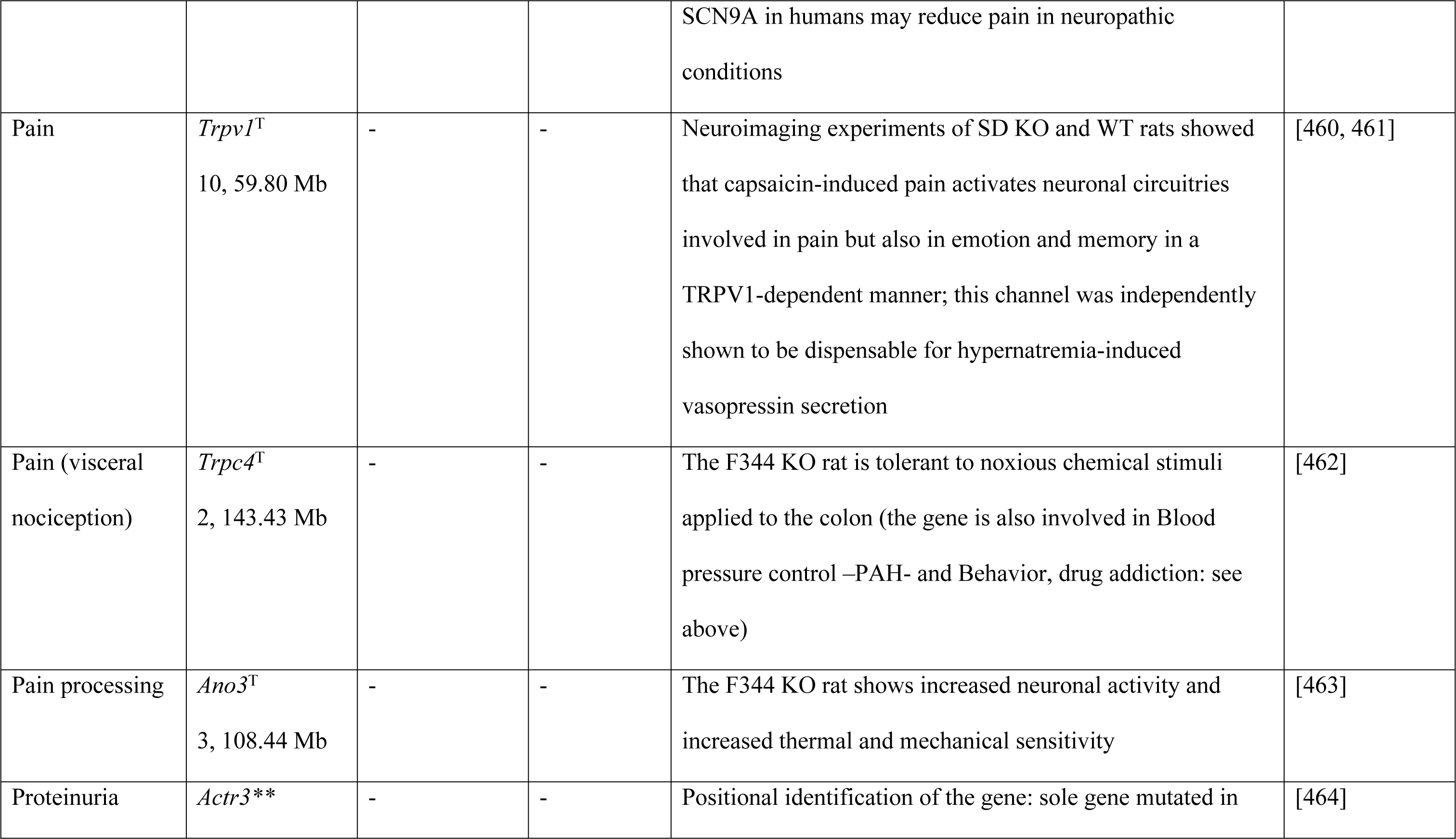
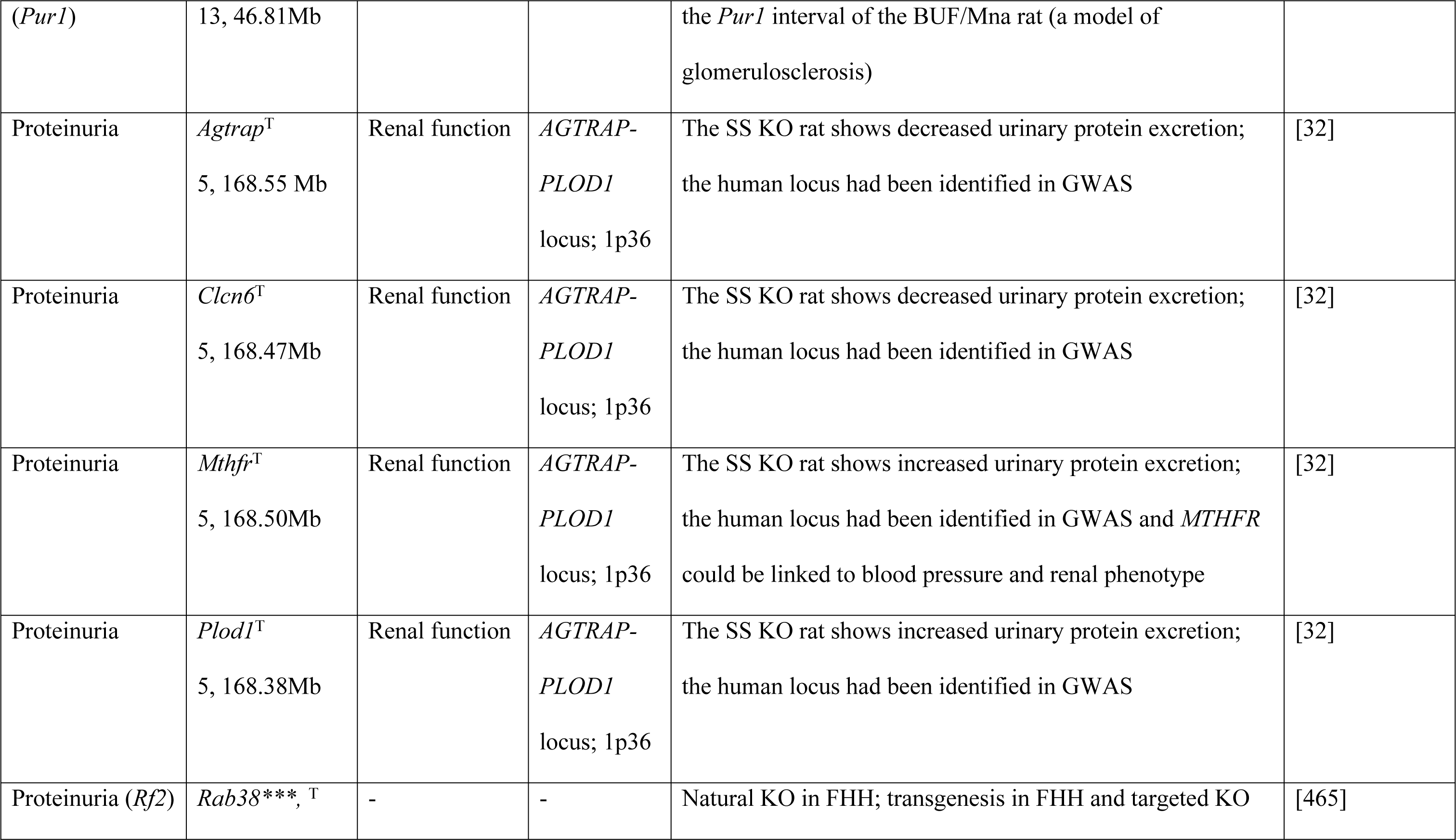
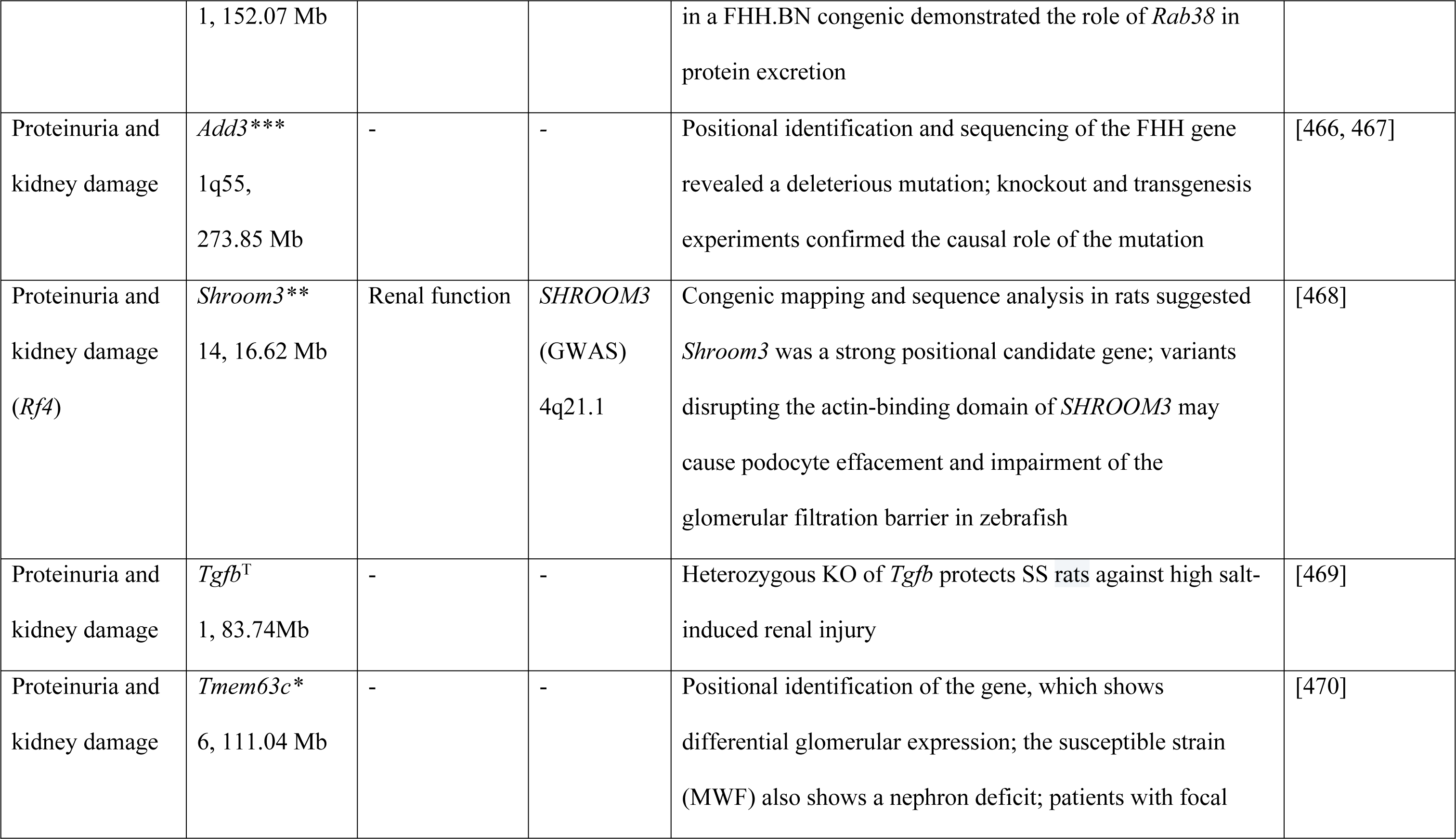
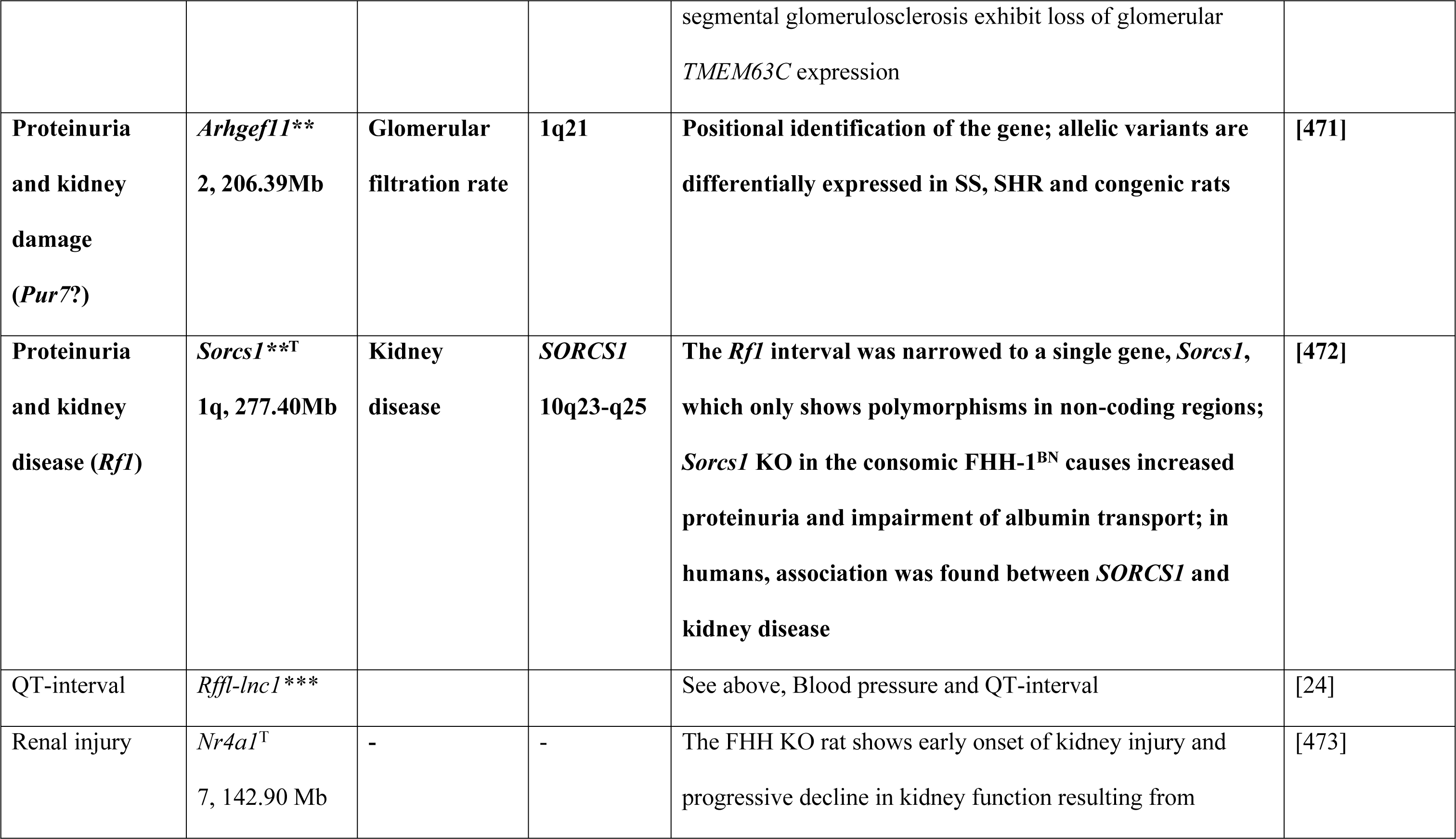
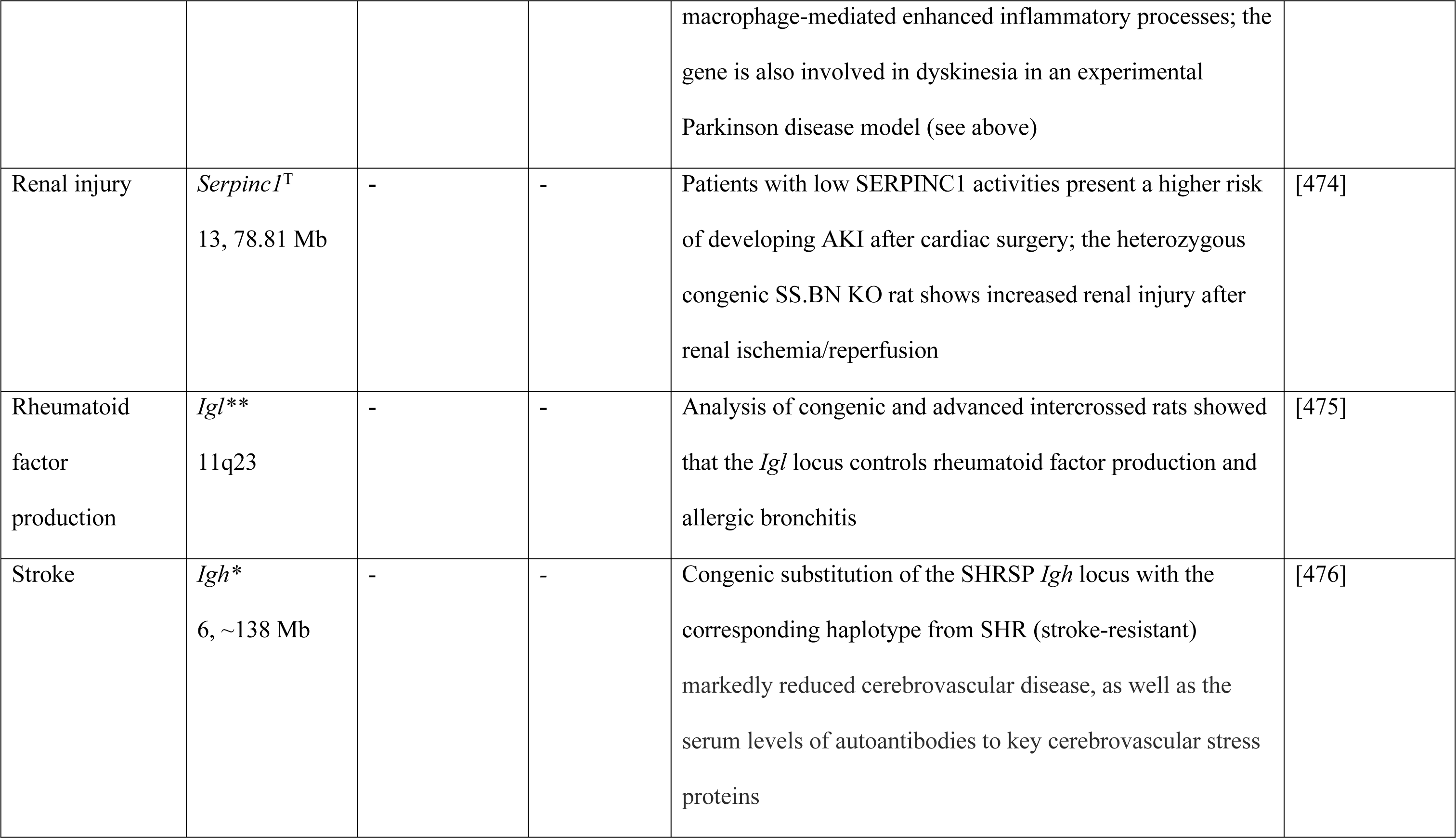
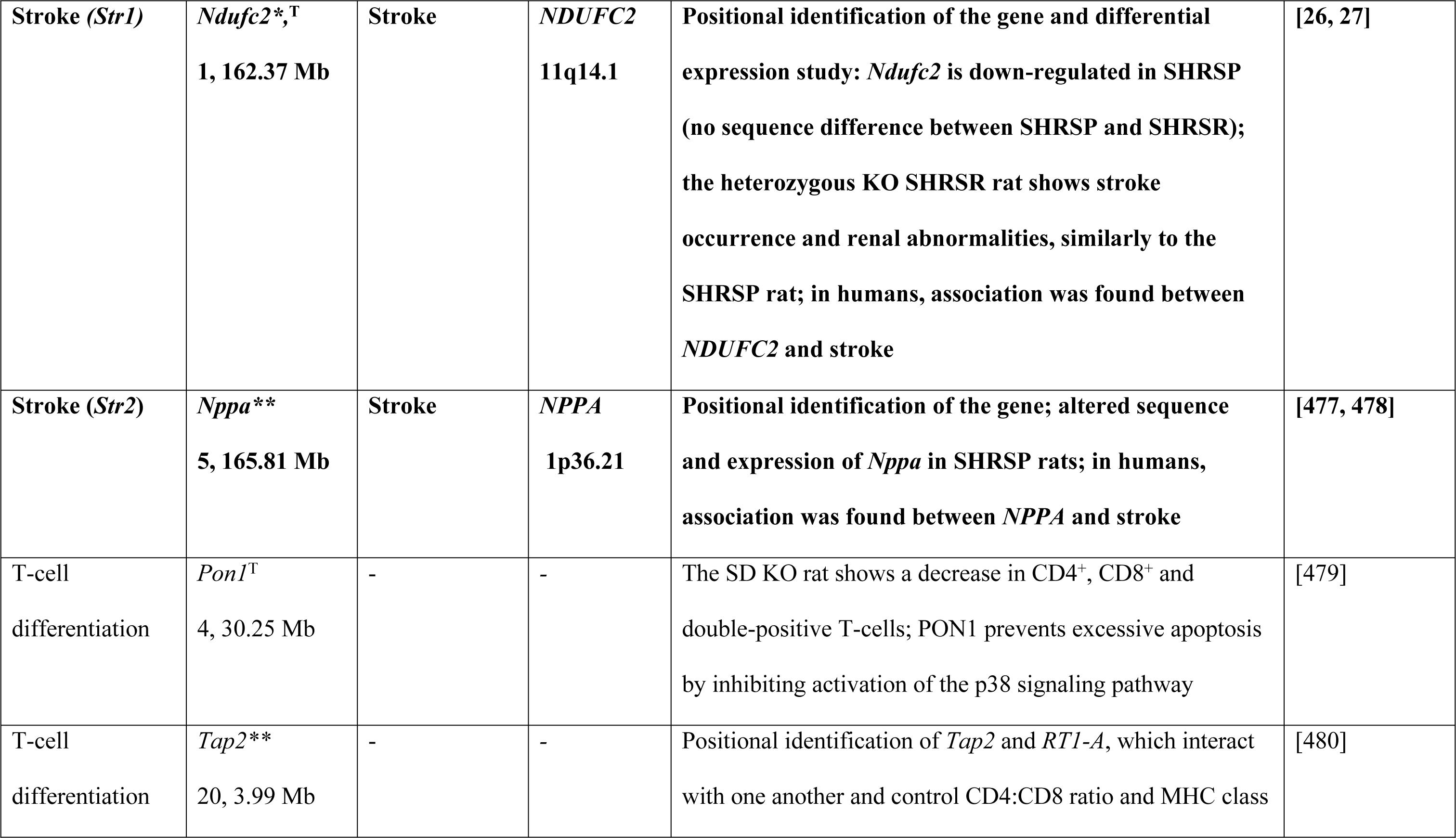
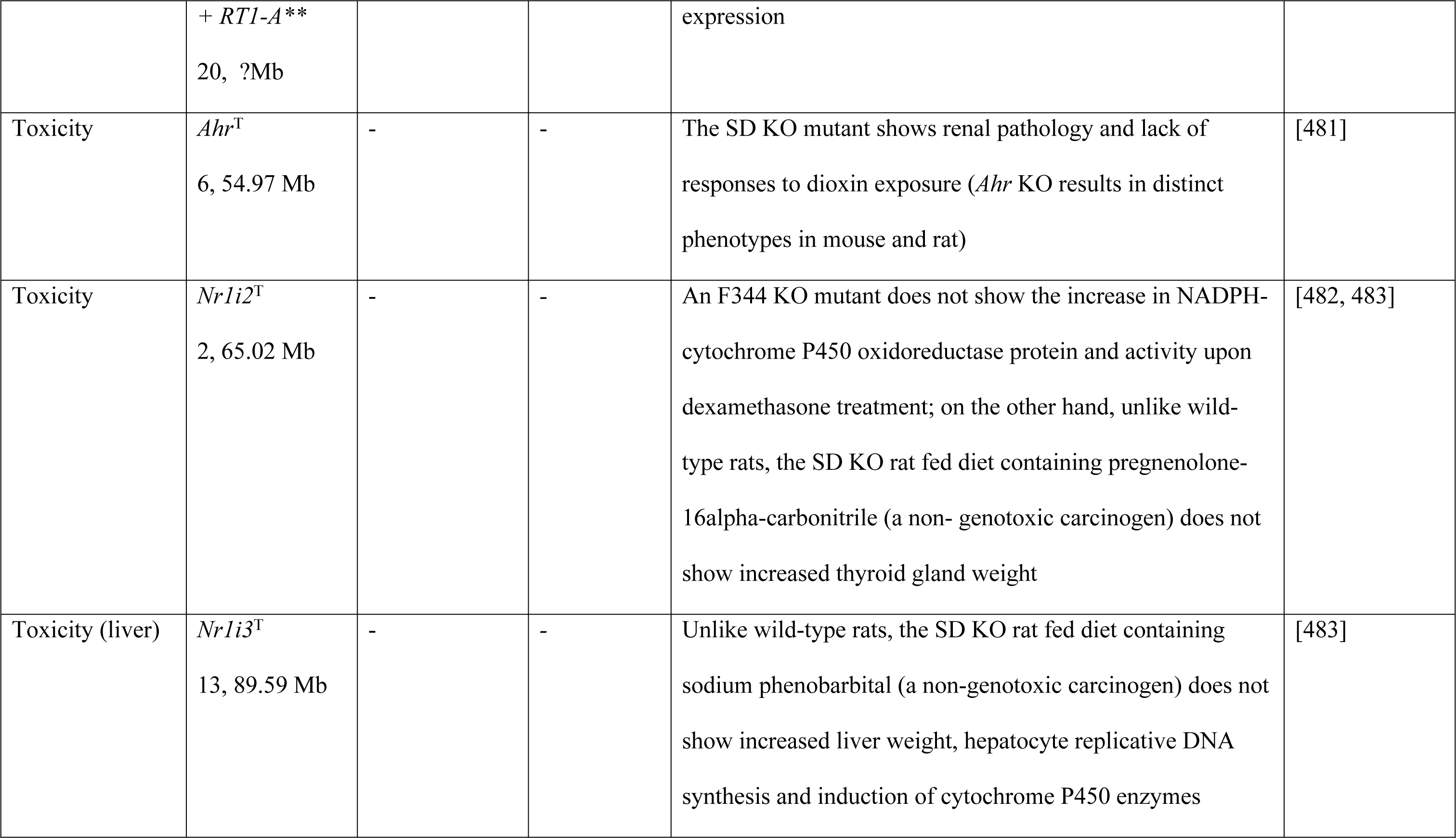
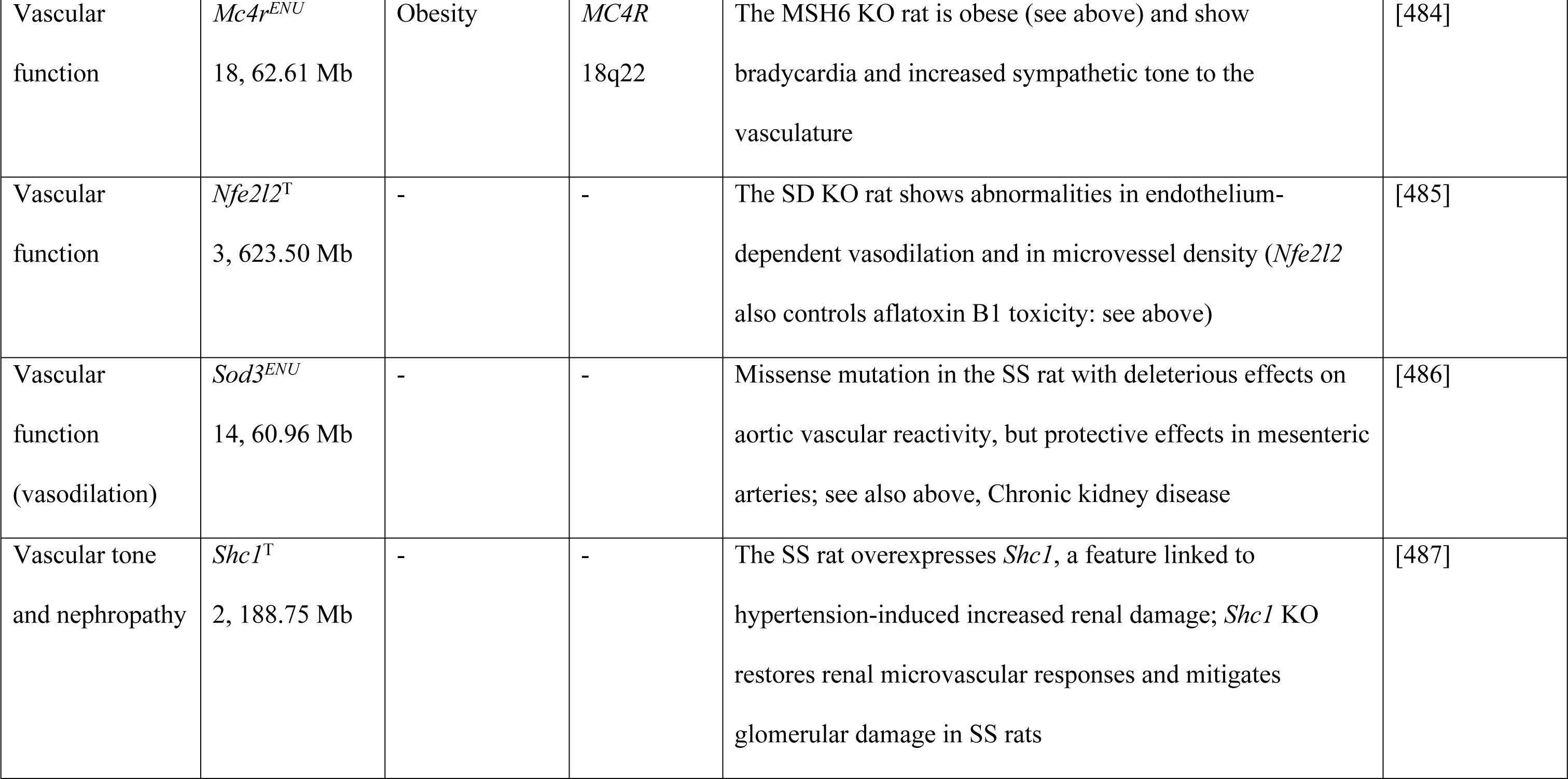
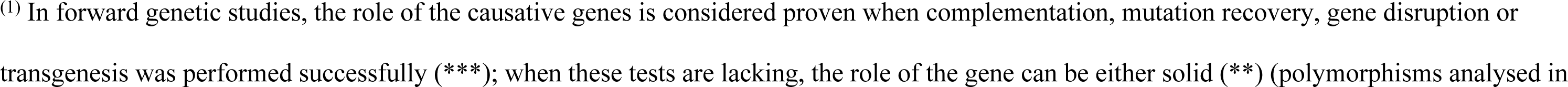
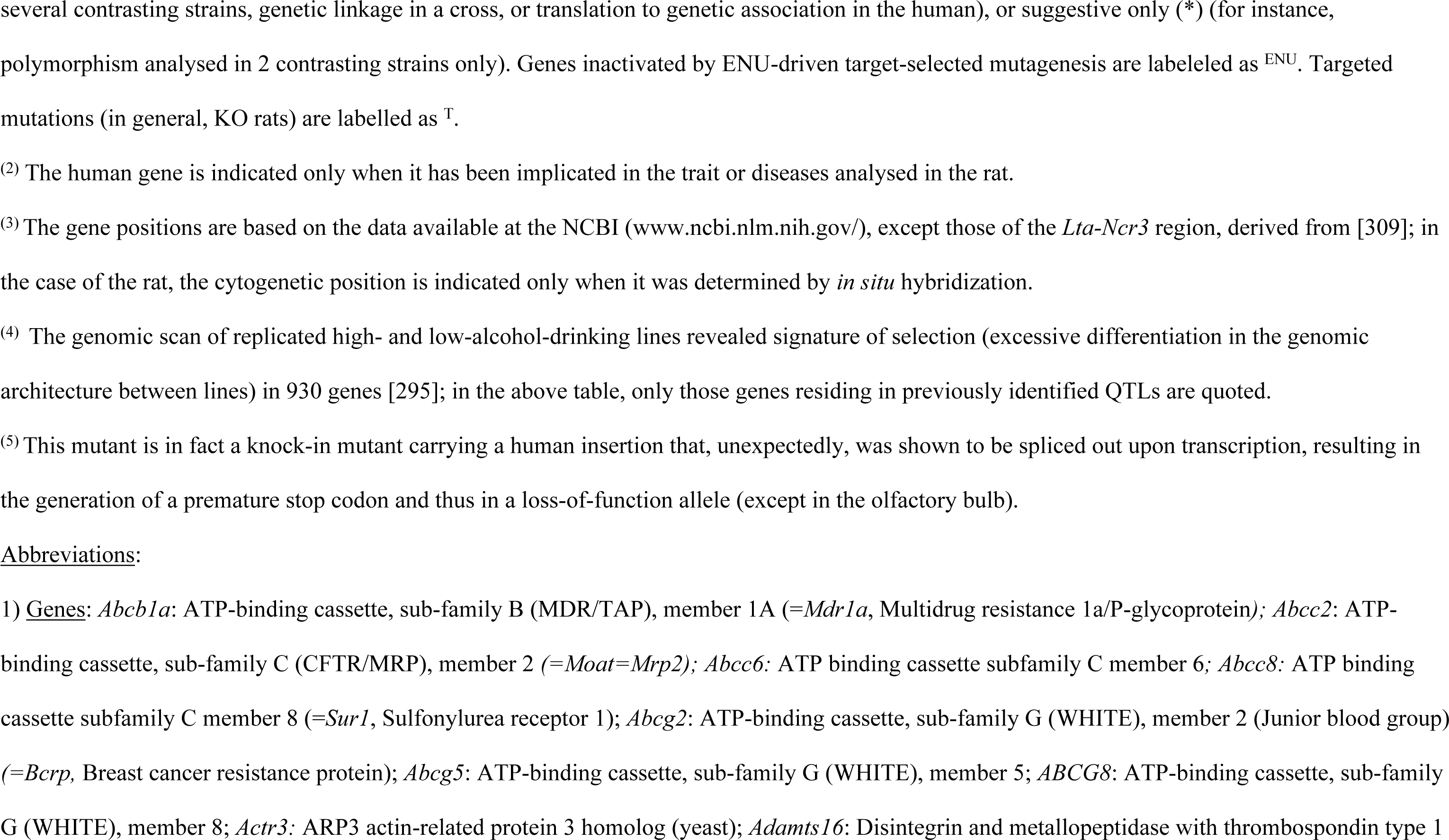
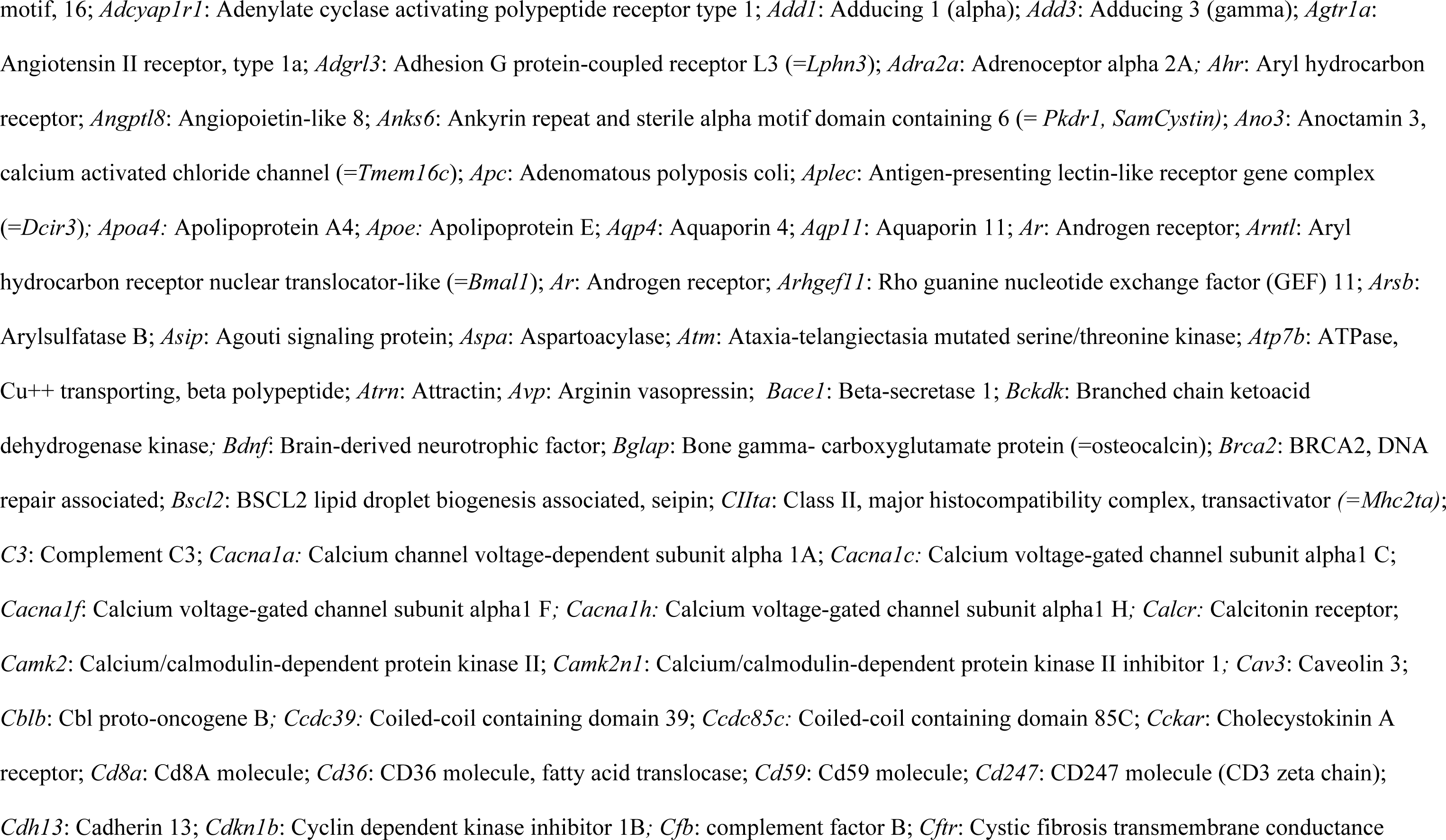
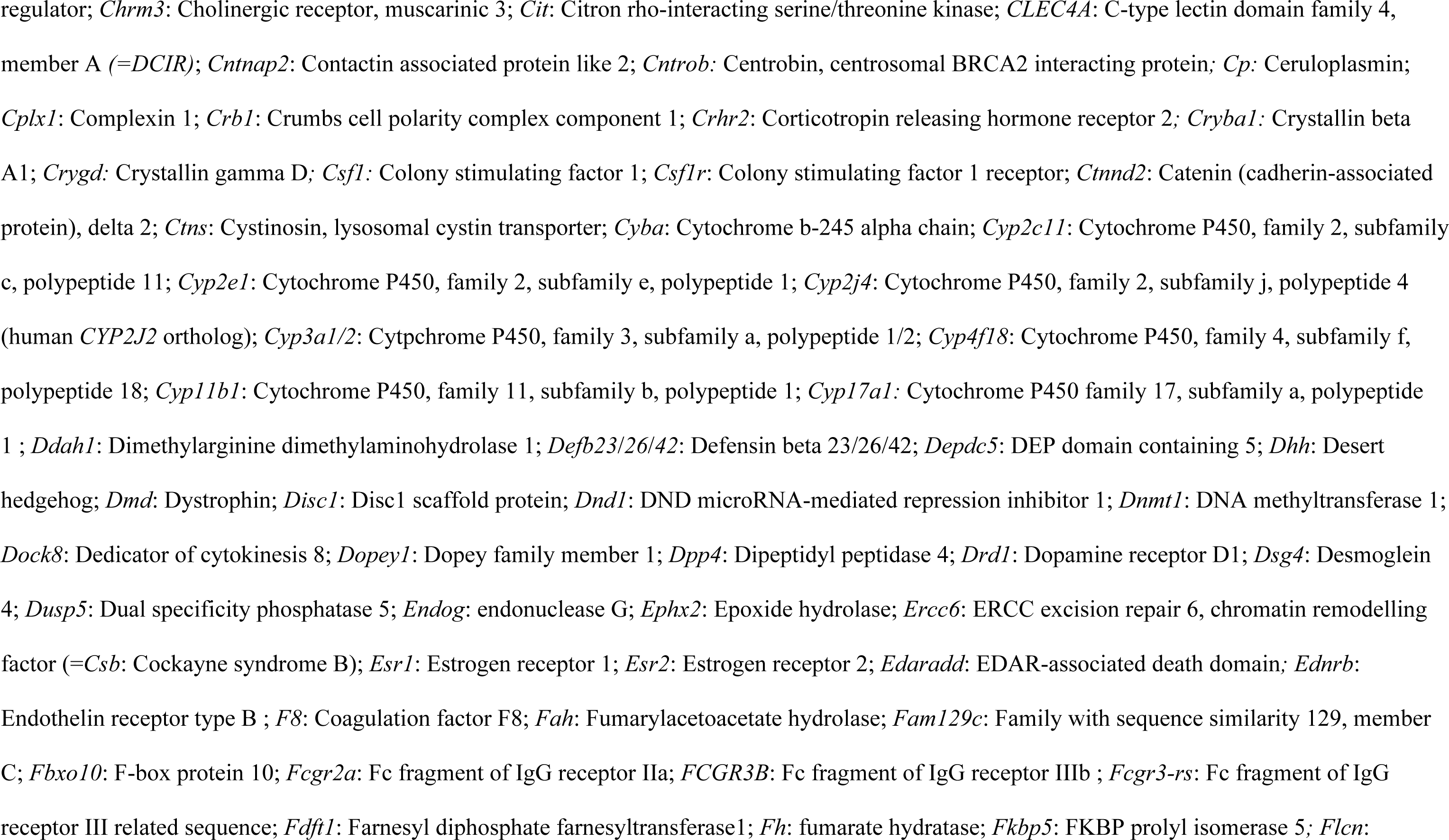
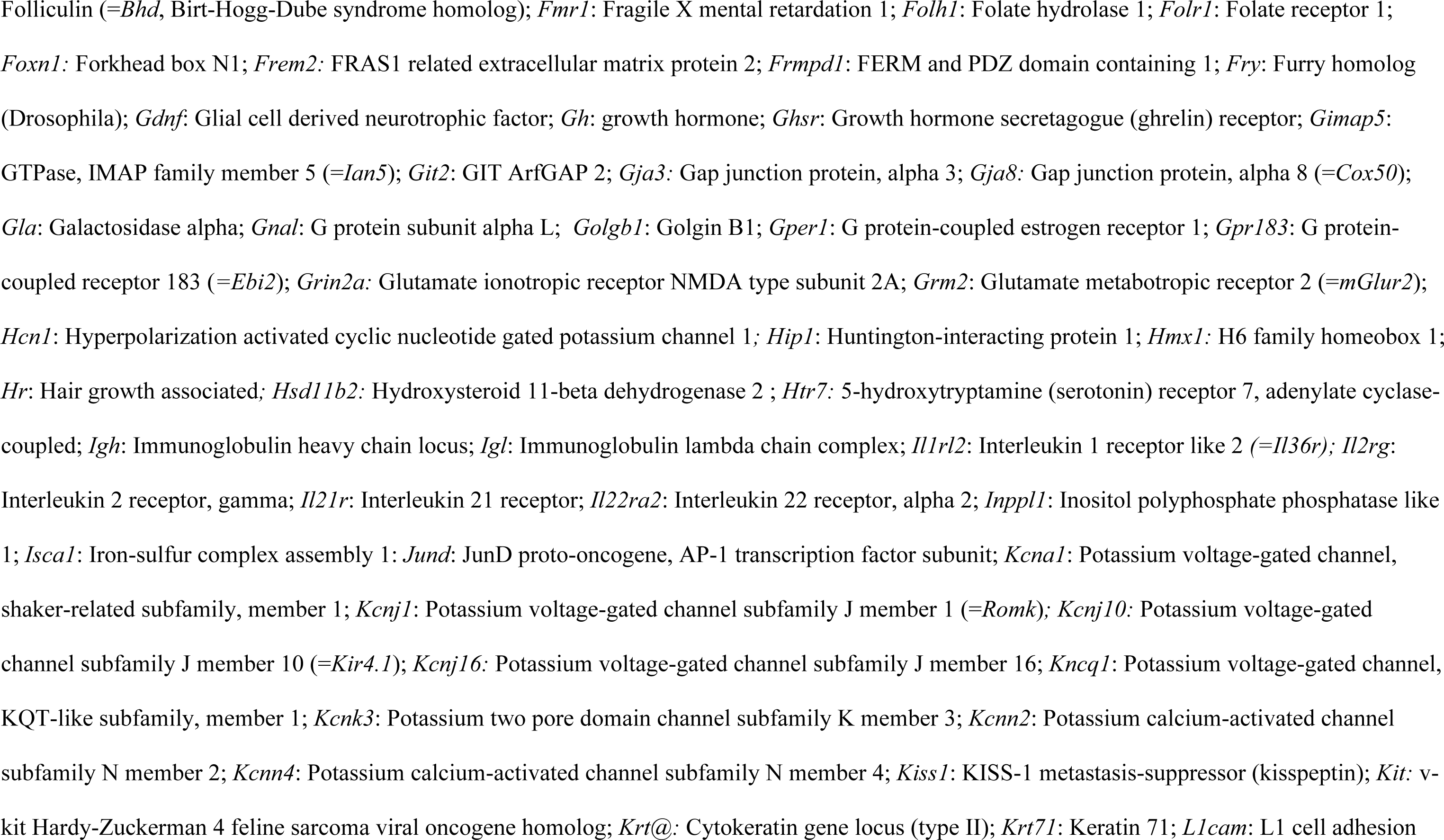
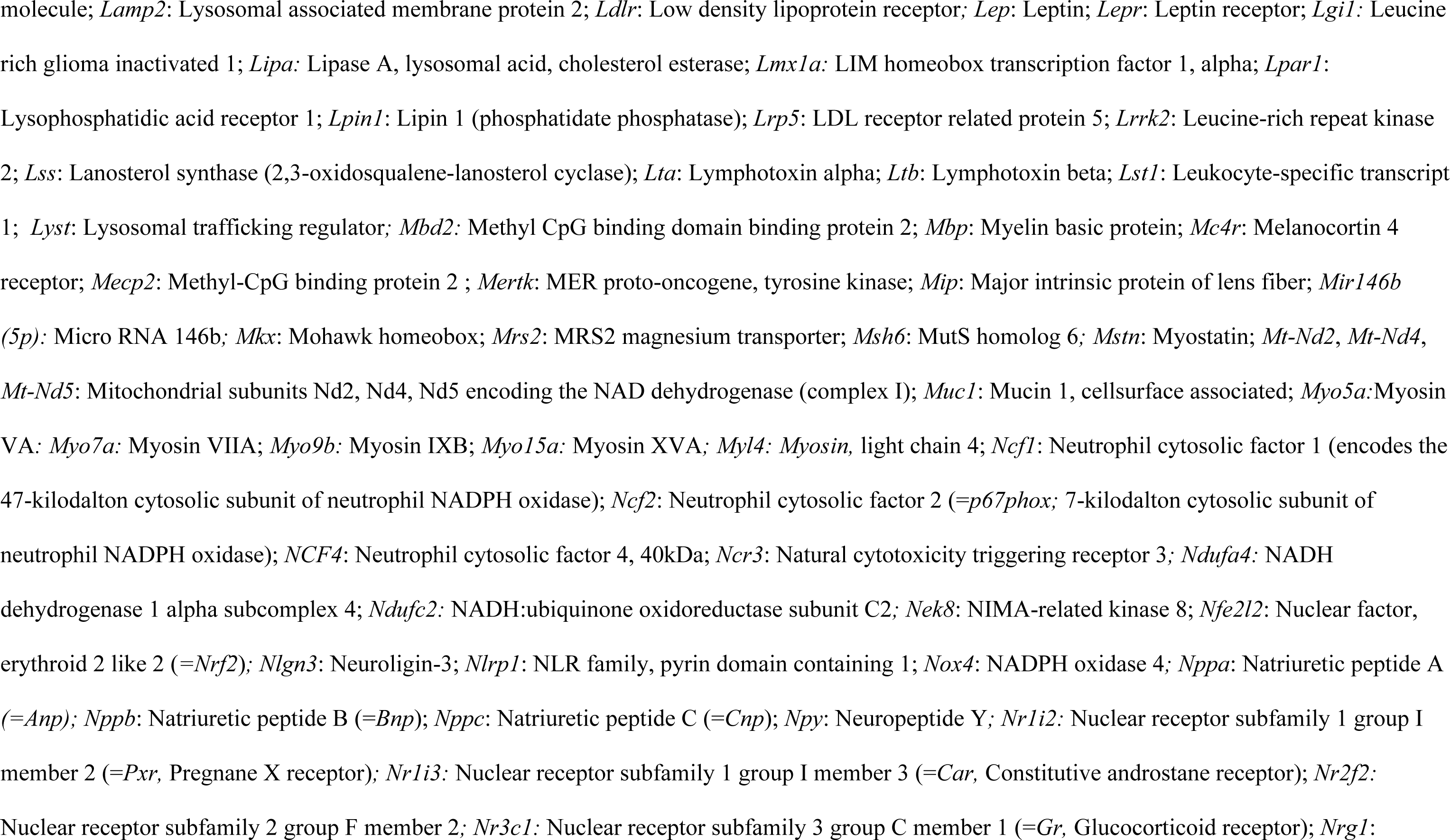
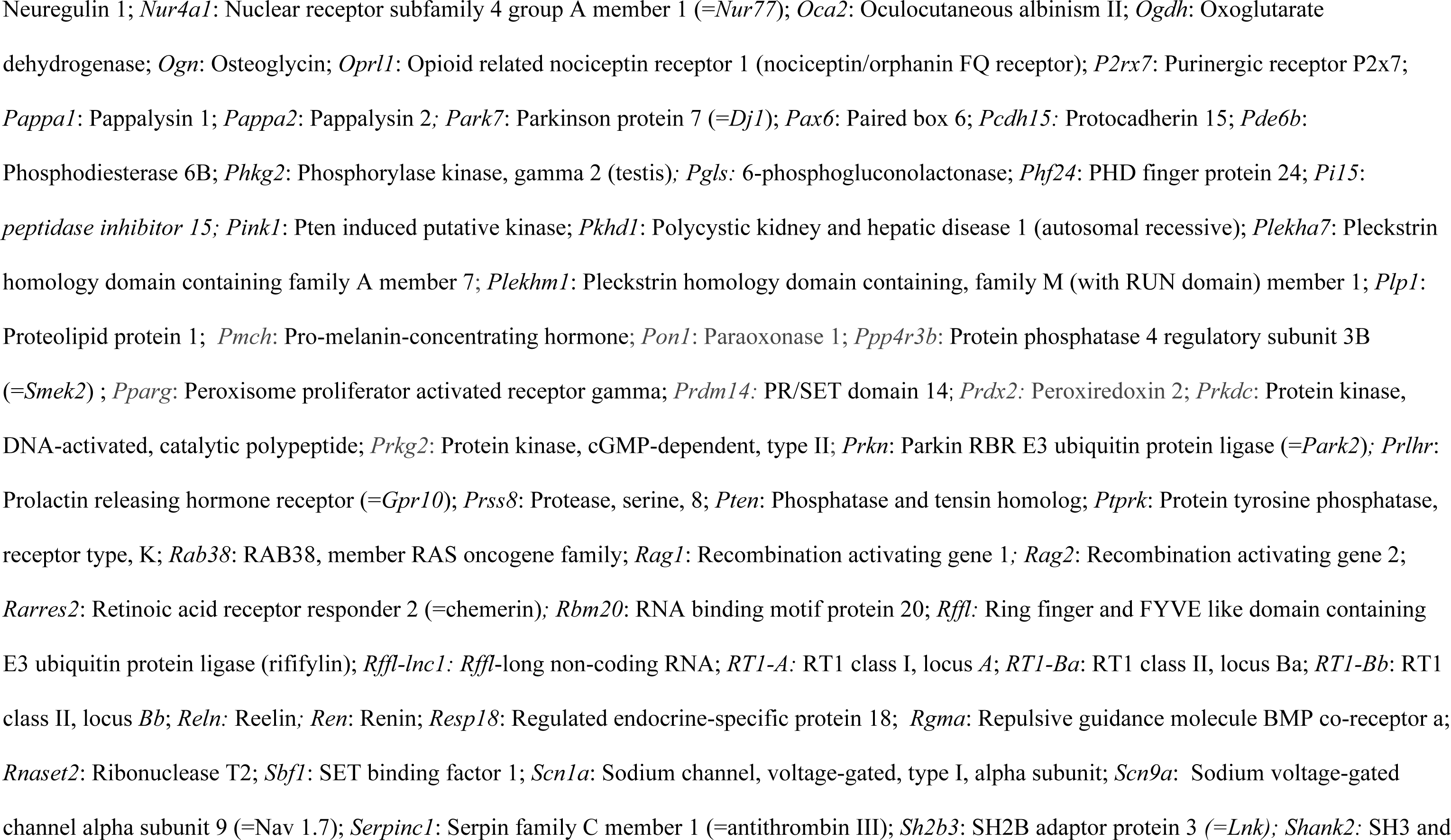
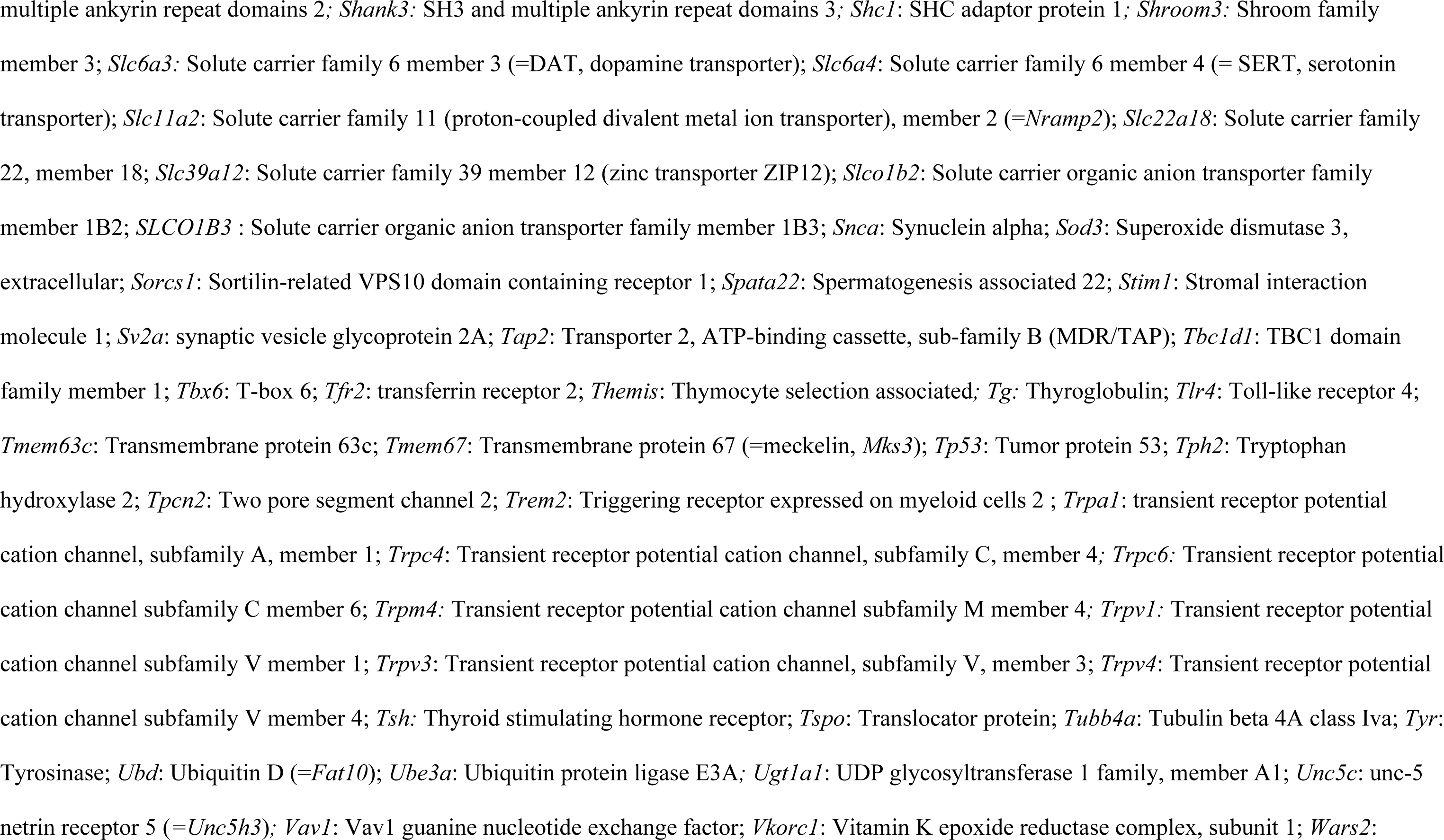
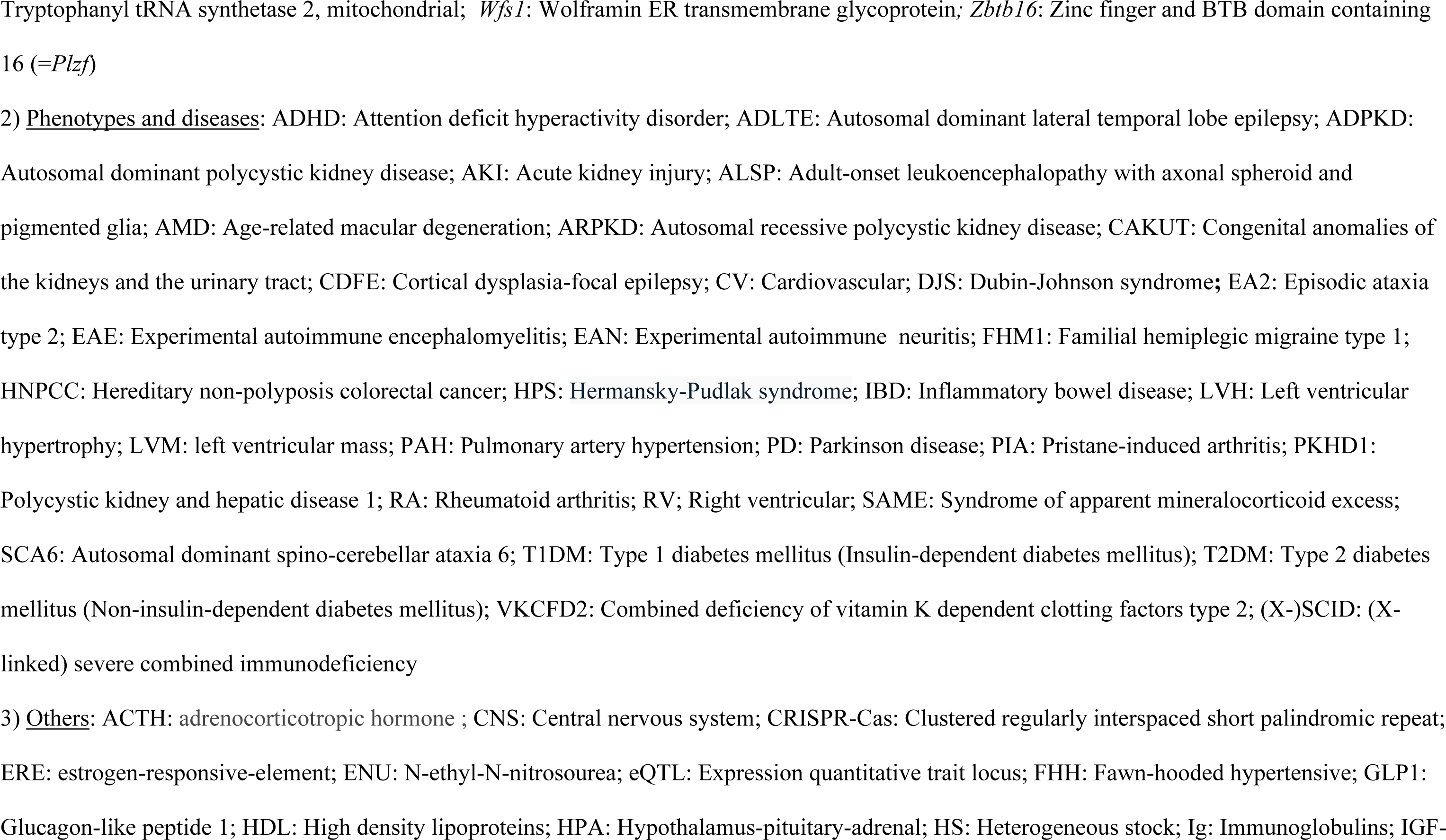
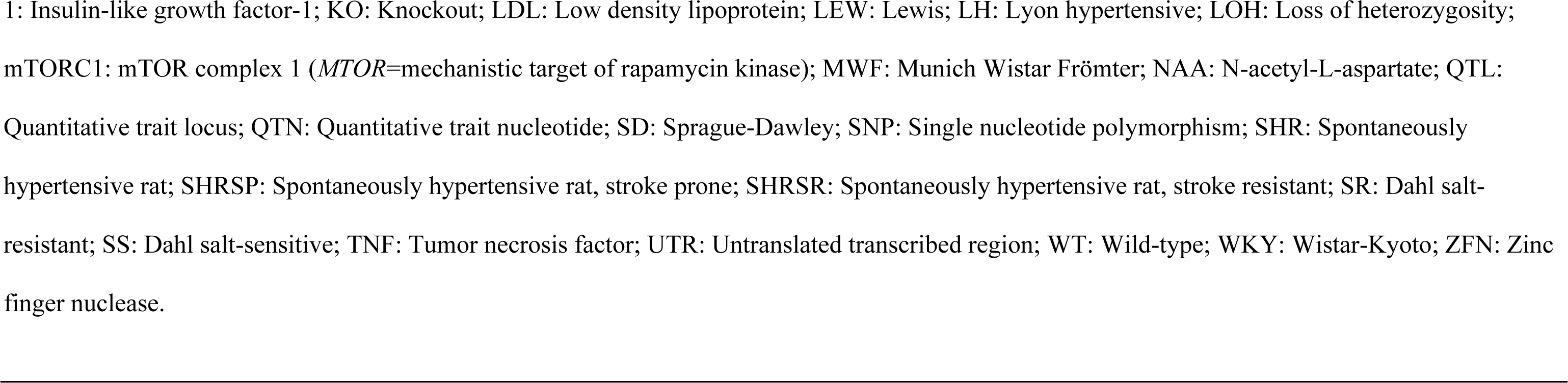
Alphabetical list of diseases and related traits with their causative rat genes and the human homologs

Table 1A shows the monogenic traits, and table 1B the complex traits (it a few cases this distinction is somewhat arbitrary, but in general this is a useful classification). Of note, when a gene was associated with several distinct phenotypes, an entry was created for each phenotype and the gene thus appears several times in the table. When the human homolog gene is known to be causal of the relevant disease or trait, it is also indicated in the table. Furthermore, entries in bold characters indicate that the human gene was found to be causal as a direct translation of the results obtained in the rat.

The identification of gene(s) underlying a given phenotype typically starts with the mapping of the trait by linkage analysis (backcrosses, intercrosses). In the case of monogenic traits, this approach is generally sufficient to identify the causative gene (positional identification, as illustrated in Table 1A). Identifying genes controlling complex traits is much more difficult [16]; indeed, linkage analyses of such traits lead to the localization of quantitative trait loci (QTLs), which are too large to allow the identification of the causative gene. Complementary strategies are thus required to narrow down the list of candidate genes, such as the generation of congenic lines or/and the use of integrative genomic approaches [as discussed in 17]. Alternative approaches rely on the use of panels of lines that show a higher level of recombinant events, as a result of crossing parental strains for multiple generations, such as recombinant inbred strains or heterogeneous stocks [as discussed in 18, for a striking harvest of results derived from the study of a heterogeneous stock, see 19]. The first complex-trait gene identified is the *Cd36* gene, which causes insulin resistance, hyperlipidemia and hypertension in the spontaneously hypertensive rat (SHR) [20, 21]. This identification was based on a combined gene expression micro-array and linkage approach and was definitively proven by in vivo complementation, i.e. transgenic expression of normal *Cd36* in the SHR [22]. Last but not least, association was then demonstrated between human *CD36* and insulin resistance [23]. Subsequently, the tools of forward genetic studies as well as gene expression and/or computational analysis (integrative genomics) led to the identification of numerous genes underlying rat polygenic traits or diseases, such as blood pressure, cardiac mass, diabetes, inflammation (in particular arthritis, encephalomyelitis), glomerulonephritis, mammary cancer, neurobehavioral traits, proteinuria. In several instances, the results were translated to the human, as illustrated in Table 1 by bold entries. Interestingly, a recently discovered complex trait gene is a long non-coding RNA, itself contained within the 5’ UTR of the *Rffl* gene (*Rffl-lnc1*); *Rffl-lnc1* shows a 19bp indel polymorphism which is the precise variation underlying regulation of blood pressure and QT-interval. This work was based on fine and systematic congenic mapping and is the first one to identify quantitative trait nucleotides in a long non-coding RNA [24]. The human homologous region, on chromosome 17, has multiple minor alleles that are associated with shorter QT-intervals and, is some cases, hypertension [25].

Identifying rat disease genes is not only useful to discover the homologous human disease genes but also helps in studying the mechanisms underlying the pathological abnormalities. After all, this is the essence of an animal model. For instance, the study of the genetic basis of stroke in the stroke-prone SHR strain (SHRSP) led to the conclusion that mitochondrial dysfunction contributes to stroke susceptibility and to hypertensive target organ damage (such as vascular damage); this better understanding of the etiology of the disease can open the door to novel therapies [26, 27]. Another example is provided by the identification of *Ncf1* as a causative gene of arthritis [28] which led to the discovery that reactive oxygen species are important regulators of several chronic inflammatory disorders and more generally of immune and inflammatory pathways; surprisingly, they have a protective role in autoimmune diseases [29].

The rat is also a useful model to decipher the biological significance of QTLs identified in human genome-wide association studies (GWAS) aimed at understanding the aetiology of common human diseases [30, 31]. These studies pint-point human genomic regions controlling a complex trait, and generally contain several genes; the current methods lack the statistical power to pinpoint the human causative gene. Animal model such as the rat provides one with the possibility to knockout or to mutate in more subtle manner each of the rat genes homolog to the human genes contained in a given GWAS locus. In this way, the possible role of each gene can be evaluated. For instance, Flister and c-corkers [32], studying a multigene GWAS locus controlling blood pressure and renal phenotypes (*AGTRAP*-*PLOD1* locus) used gene targeting in a rat model to test each of the genes contained in this locus. In this way these authors could show that several genes impact hypertension and that multiple causative gene variants cosegregate at this locus; several linked genes thus control blood pressure (*Agtrap*, *Clcn6*, *Mthfr*, *Nppa*, *Plod1*). Furthermore, each of the KO rat models so generated can be used to dissect the biological effects of the gene loss of function.

The genetic basis of human diseases is also actively analyzed by whole genome sequencing; such studies have uncovered several genes underlying diseases or related phenotypes [33, 34] and one can thus questioned the importance of genetic analyses in an animal model. As argued and illustrated above, animal models and the rat in particular, remain valuable tools to analyze the biological mechanisms underlying a phenotype. In addition, transgenesis or gene substitution can also be carried out, in which a human allele can be introduced in the relevant KO rat, in order to verify the role of the human mutation. Alternatively, the rat genome can be directly modified to specifically introduce a mutation similar to the one causing the human trait [34, 35]. If the modified rats exhibit defects similar to those observed in the human patients, it can be concluded that the tested human mutation indeed plays a causal role. In addition, similarly to examples mentioned above, such specifically modified rats provide one with models suitable to study the mechanisms responsible for the abnormalities generated by the mutation and also to carry out pharmacological tests and look for possible new therapies [35].

The need of relevant animal models is also illustrated by the fact that even when the human gene causing a disease is identified, mutated rat strains (in particular KO strains) are created to analyze the gene function and the disease pathogenesis (see numerous examples of such gene targeting in Table 1). In 2008, Aitman and coworkers [2] reported a list of 21 rat disease genes that had been identified by positional cloning since 1999. Here I included all genes, independently of the date of their identification. This inventory added a few disease genes identified before 1999 but mainly numerous genes identified (or deliberately mutated) after 2008. The total rat gene number listed here is over 300, illustrating the vigor of the rat biomedical research which led to enrichment of numerous disease models, with the translation to humans of disease gene discoveries in rats.

## Acknowledgments

The author thanks Jennifer Smith (RGD) for advice in extracting relevant data from RGD, in particular using the ontology browser for disease (https://rgd.mcw.edu/rgdweb/ontology/view.html?acc_id=DOID:4). The author is an Honorary Research Director of the FNRS (Belgium).

